# Self-inspired learning to denoise for live-cell super-resolution microscopy

**DOI:** 10.1101/2024.01.23.576521

**Authors:** Liying Qu, Shiqun Zhao, Yuanyuan Huang, Xianxin Ye, Kunhao Wang, Yuzhen Liu, Xianming Liu, Heng Mao, Guangwei Hu, Wei Chen, Changliang Guo, Jiaye He, Jiubin Tan, Haoyu Li, Liangyi Chen, Weisong Zhao

## Abstract

Every collected photon is precious in live-cell super-resolution (SR) fluorescence microscopy for contributing to breaking the diffraction limit with the preservation of temporal resolvability. Here, to maximize the utilization of accumulated photons, we propose SN2N, a <u>S</u>elf-inspired <u>N</u>oise<u>2N</u>oise engine with self-supervised data generation and self-constrained learning process, which is an effective and data-efficient learning-based denoising solution for high-quality SR imaging in general. Through simulations and experiments, we show that the SN2N’s performance is fully competitive to the supervised learning methods but circumventing the need for large training-set and clean ground-truth, in which a single noisy frame is feasible for training. By one-to-two orders of magnitude increased photon efficiency, the direct applications on various confocal-based SR systems highlight the versatility of SN2N for allowing fast and gentle 5D SR imaging. We also integrated SN2N into the prevailing SR reconstructions for artifacts removal, enabling efficient reconstructions from limited photons. Together, we anticipate our SN2N and its integrations could inspire further advances in the rapidly developing field of fluorescence imaging and benefit subsequent precise structure segmentation irrespective of noise conditions.

## INTRODUCTION

Fluorescence microscopy of live cells requires gentle imaging conditions and adequate spatiotemporal resolution to record authentic biological information, thus the photon budget is usually limited. By encoding super-resolution (SR) information via specific optics and fluorescent on-off indicators, the SR techniques have further strengthened the spatial resolution^1^ and enabled previously unappreciated, intricate structures to be observed^2–5^. However, for a finite number of fluorophores within the cell volume, the increase in spatial resolution leads to the rise in illumination intensity or exposure time by order of magnitudes to maintain the signal-to-noise ratio (SNR)^6^. Furthermore, any increase in spatial resolution must be matched with an increase in temporal resolution to prevent motion artifacts^7, 8^. This is particularly challenging for live-cell SR imaging to accumulate sufficient photons.

Because of the hardware-limited photon budget, computationally boosting the SNR is essential to maximize the utilization of sensor-collected photons. By modeling image-formation process, classical denoising algorithms based on numerical filtering and mathematical optimization can remove the noise from fluorescence images to a certain extent^9–12^. However, the corresponding carried assumptions are not dependent on the specific content of the imaging data and cannot fulfill the optimal performance. Therefore, to capture the full statistical complexity of data, the field has witnessed a sudden surge of data-driven methods, producing the unprecedented restoration results^13^. After iterative training on a dataset with ground truth (GT) labels, deep neural networks (DNNs) can learn the mapping between noisy images and their clean counterparts^14–16^. Intuitively, for supervised learning, the collection of abundant content-matched clean images is crucial, which is of great challenge to live-cell applications, especially under the SR scale.

To denoise images without clean ones, Noise2Noise (N2N)^17^ learns a mapping between pairs of independently degraded versions of the same image, and its performance can approach to that of supervised learning methods. Nevertheless, the need for the twin noisy pairs is still against the live-cell SR applications. By leveraging the pixel-wise independence of noise, several approximated forms of N2N have been developed to denoising without paired data^18–21^. However, these approximations may lead to downgraded performances. To realize the full form of N2N configuration, the DeepCAD^22^ used a temporally interleaved self-supervised data generation process to create the required noisy data pairs, by assuming that the two adjacent frames in a video of continuous imaging can be considered with the same underlying content. Unfortunately, although this assumption is routinely satisfied in calcium imaging^22, 23^ or other few fast-imaging applications^24^, it is still hard to accomplish in live-cell SR techniques considering the commonly compromised temporal resolution and increased spatial resolution.

On the other hand, SR images are usually with more than sufficient sampling rate, at least over the Nyquist sampling theorem to protect the enriched spatial information. Thus, we create a self-supervised data generation strategy based on this spatial redundancy, using a diagonal resampling step followed by a Fourier interpolation. Compared to the previous temporal resampling^22–24^, this spatially interleaved data generation is more universal and stable, and it produces almost no bias in our tests. Beyond this spatially self-supervised N2N realization, we also develop a self-constrained learning process to further enhance the denoising-performance and data-efficiency. Conceptually, the two network predictions of the generated noisy pair will be constrained to one identical expectation, and this process shrinks the learning and predictive uncertainty, inherently increasing the efficiency and effectiveness. Together, our self-inspired N2N (SN2N) reaches or even outperforms the supervised learning to denoise methods with less training data, in which only one single frame is sufficient for training.

To showcase the broad applicability, we apply our SN2N directly to two commercial spinning-disk confocal-based structured illumination microscopes (SD-SIM)^25, 26^ and two commercial stimulated emission depletion (STED)^27, 28^ microscopes, as well as expansion microscope (ExM)^29^ and high-resolution confocal microscope, enabling high-quality long-term multi-color 3D live-cell SR imaging (5D in *xyz*-*c*-*t*). Beyond that, we also integrate our SN2N framework into the existing SR reconstruction procedures, including the iterative deconvolution on SD-SIM and STED, SR optical fluctuation imaging (SOFI) reconstruction^30, 31^, and SIM^10^, effectively mitigating the artifacts embedded in the SR images and enhancing the photon efficiencies. These organic integrations deliver the fact that our SN2N can serve as a practical framework for further strengthening the performance of live-cell SR microscopy. With fully open-sourced codes, we also expect our SN2N will be applicable to other fields for random noise removal.

## RESULTS

### Principle of SN2N

#### Self-supervised data generation by considering the inherent characters of SR microscopy

Before designing the data generation strategy, we intend to list several physical properties (**p**) of SR microscopy^32^: **p1)** To fulfill the increased spatial resolution, the sample rate is routinely finer than the Nyquist theorem. **p2)** The point spread function (PSF) represents the highest frequency of the system and is usually with spatial symmetry inside 2 × 2 pixels; **p3)** Commonly, each pixel of the camera is sampled independently; **p4)** The imaging system can be regarded as an equivalent low-pass filter. In this case, each SR image is composed of many PSFs multiplied with the corresponding fluorophore brightness at different spatial positions, and each PSF occupies at least larger than an area of 2 × 2 pixels^12^. Considering this spatial redundancy (**p1** and **p2**), we can directly resample one SR image as two subimages by an unusual form of binning we regard as ‘diagonal re-sampling’ (**Fig. 1a**). In specific, we consider every 4 adjacent pixels (2 × 2) as one unit, and every two diagonal pixels (top left and bottom right, magenta color in **Fig. 1a**; top right and bottom left, green color in **Fig. 1a**, **Extended Data Fig. 1a**) are averaged as one new pixel. Because of the independency of each pixel and the spatial symmetry on the two diagonal directions (**p2** and **p3**), we can generate statistically independent image pairs that share identical details but different noise realizations for the N2N configuration (**Methods**). Fundamentally, assuming the identical contents inside one smallest unit, our method can be seen as a relaxation form of the temporal resampling approach^22–24^, which requires the entire contents of two frames being identical. Compared to other spatial generation methods^22, 33^, this resampling in diagonal axes is more stable as requiring no additional registration or calibration to create perfectly matched image content. Notably, although originated from the spatial redundancy of SR images, our approach is not sensitive to the changes of pixel sizes and can produce stable denoising results for even undersampled images (until 260 nm pixel size with 150 nm resolution, ↓8 in **Extended Data Fig. 2a-2b**).

**Fig. 1.**
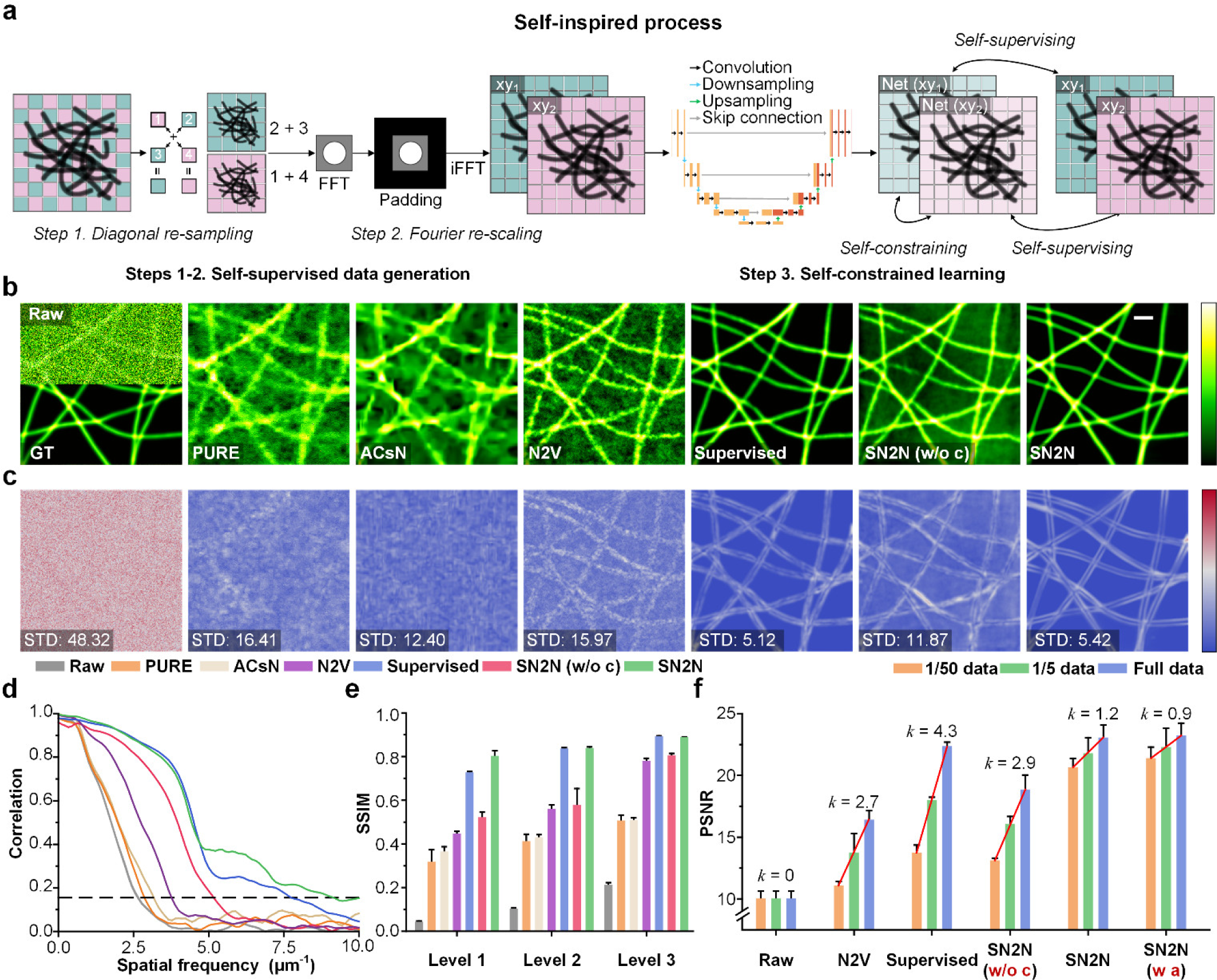
Workflow and simulation validation of SN2N. **a**, Overview of SN2N. Steps 1-2, self-supervised data generation. The single-frame input image of H × W pixels is diagonally re-sampled to two images of H/2 × W/2 pixels. Then, the two resulting images are re-scaled back to two images of H × W pixels with a Fourier interpolation. Step 3, self-constrained learning process, i.e., the two re-scaled images serve as both inputs and labels, and the corresponding two predicted images from a classical U-Net architecture are constrained to minimize the difference between them (**Methods**). **b**, Validation of SN2N using synthetic microtubule structures (**Methods**). The synthetic structures were convoluted with a 150 nm PSF and down-sampled 2 times (pixel size 32.5 nm) as ground truth. The noisy images were created by further injection of Poisson noise and 40% Gaussian noise. From left to right: noisy (top) and ground-truth images (bottom), PURE, ACsN, N2V, supervised, SN2N without self-constrained loss (SN2N w/o c), and SN2N denoising results. **c,** Data uncertainty results of **b**. Ten independent frames of identical contents under the same imaging conditions were fed to the trained SN2N network, and the standard deviation (STD) of the ten resulting predictions was calculated as the data uncertainty. Marked numbers are the average values of STD maps. **d**, FRC analysis of the denoising images. The dashed line represents the FRC threshold. **e,** SSIM values of various denoising methods under different noise levels (*n* = 10). Images of Levels 1-3 conditions were injected with 40%, 20%, and 10% Gaussian noise as well as the corresponding Poisson noise, respectively. **f**, PSNR values of networks trained by different amounts of data under the Level-1 condition (*n* = 10). Full data dimensions: 2048 × 2048 × 50. *k* denotes the slope (red lines) of PSNR value along the data increment. Error bars, s.e.m. Experiments were repeated ten times independently with similar results; scale bar, 1 µm (**b**).

Because these produced two subimages are twice smaller, we further adapt an interpolation method to rescale them to the original structural scale (**Methods**). Different from the conventional spatial methods, we apply a Fourier interpolation to the resulting two subimages, which is based on the fact that the optical transfer function (OTF) of an SR microscope has only finite support (**p4**). Padding the Fourier-transformed image out of its OTF support with zeros does not alter its information content^34^, and after back-transforming, this padded image will have doubled pixel number in both *xy* axes, identical to the original SR image (**Fig. 1a**, **Extended Data Fig. 1a**). Without this operation, the network may produce structural artifacts for the scale difference between the training and test datasets (**Extended Data Fig. 2c**). On the other hand, the spatial methods (such as bilinear interpolation) create more pixels according to the noise-corrupted pixels and it will be problematic when meeting the background areas without fluorescence signal. Because these smoothly created pixels do not conform to the randomness of noise, which may influence the learning process, potentially producing background artifacts (**Extended Data Fig. 2c**).

#### Self-constrained learning process

Beyond the full form of N2N, we further design a self-constrained learning process to constrain the training process and denoising variance (**Fig. 1a**). This is based on a simple intuition that the denoising predictions of the same underlying signal should be identical. In our case, the generated data pair has matched content but different noise components, thus ideally, the corresponding two denoised results should have no differences. In specific, the two generated images are successively and individually fed into the network, and the two resulting predictions can be calculated by the following loss function to execute the training stage.

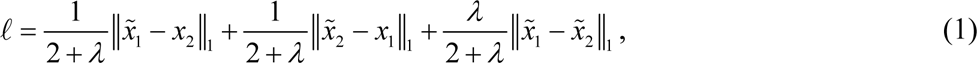

where 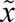 represents the network outcome of input *x*, and ‖ ‖ refers to some kind of distance calculations. We used the *l*_1_ norm as distance calculation (**Methods**). The first two terms are the conventional N2N losses, and the last term is our self-constrained loss with a constraint weight *λ*. By enforcing this constraint, we found that the low-frequency components manifest a greater propensity of being stable than the high-frequency ones (**Extended Data Fig. 2e-2f**). Thus, to avoid over-smoothed results, we routinely set the constraint weight as 1. Notably, our SN2N is model-independent, thus we employed the commonly used U-Net^35^ to highlight its ability (**Fig. 1a**, **Extended Data Fig. 1a**, **Methods**).

#### Data augmentation for low-level tasks

Although there are plenty of strategies to increase the training data amount, they are designed for high-level classification tasks and not suitable for low-level denoising applications^36^. Accordingly, we develop random patch transformations in multiple dimensions (Patch2Patch) to further improve the data efficiency (**Extended Data Fig. 1b**). Specifically, the random selected ROIs (regions of interest) on each image will be rotated, flipped, or remained still and subsequently interchanged with other ROIs from the same frame or a different frame from different time or experiments (**Methods**, **Supplementary Video 1**). This Patch2Patch can create more imaging results without changing the inherent noise properties and hence it can effectively reduce the required data bulk.

### Benchmarking with known structures

#### Simulation validation

To quantitatively test the SN2N’s performance, we first validated it on synthetic microtubule imaging data with 150 nm resolution and three different Poisson-Gaussian noise levels (**Methods**, **Supplementary Fig. 1**). Our SN2N solution is superior to existing methods in two aspects. First, SN2N is denoising effective, especially for ultra-low SNR conditions. Under the lowest SNR condition (Level 1), classical denoising methods, e.g., PURE (Poisson unbiased risk estimate)^9^ and ACsN (automatic correction of sCMOS-related noise)^11^, failed to eliminate the noise and left overly blurred underlying structures (**Fig. 1b**). The approximated form of N2N, Noise2Void (N2V)^19^, can reduce the noise, but the predicted images from ten repetitive acquisitions are still with large standard deviation (STD as 15.97 in **Fig. 1b**), reflecting the limited performances. Removing the self-constrained learning process (SN2N w/o c), the improvement of our method against N2V is relatively limited and cannot be competitive with the supervised learning method using sufficient training data. The full SN2N truly approaches the performance of supervised learning with similar STD and even better FRC (Fourier ring correlation)^37^, SSIM (structural similarity)^38^, PSNR (peak SNR), and RMSE (root-mean-square error) metrics (**Fig. 1b-1e**, **Extended Data Fig. 3b**). On the other hand, when examining the higher SNR conditions (Level 2 and Level 3), all methods can achieve acceptable denoising results (**Fig. 1e**, **Extended Data Fig. 3a**) and the improvements are less impressive, in which the results of N2V and SN2N without constraint (SN2N w/o c) reach closer to the full SN2N (**Supplementary Table 1**).

Second, our SN2N is data efficient and can be trained even on one single image. The requirement of large datasets for DNNs to capture the accurate data distribution of high noise-level is moderated by our self-constrained learning process. To test this, we measured the slopes of SSIM, PSNR, and RMSE metrics of different learning-based methods by decreasing the training data size by a factor of 5 and 50 (1 frame only) (**Fig. 1f**, **Extended Data Fig. 3c-3d, Supplementary Table 2**). As the data pool shrinking, the denoising quality of the supervised learning method dropped quickly, e.g., *k* = 4.3 for PSNR (**Fig. 1f**), and this variation also dramatically effected the performances of N2V (*k* = 2.7) and SN2N w/o c (*k* = 2.9). In contrast, it is clearly observed that the full SN2N moderated the influence (*k* = 1.2). Our self-constrained loss helps the network to learn the denoising process more efficiently, which is further strengthened by the developed Patch2Patch (*k* = 0.9, SN2N with augmentation, SN2N w a). Fundamentally, these two aspects are highly correlated to the data and model uncertainties of the DNNs^39^, in which the denoising-effectiveness reflects the data uncertainty, and data-efficiency represents the model uncertainty. Using ten acquisitions’ predictions and ten repetitively trained models’ predictions, we measured the data and model uncertainties, respectively. Consistently, it can be seen that our SN2N outperforms other methods (**Extended Data Fig. 3e**, see also **Supplementary Figs. 2-3**).

#### Experimental evaluation using standard sample under SD-SIM

SR confocal microscopy can double the spatial resolution by narrowing its pinhole size, but correspondingly the number of photons reaching the detector is severely restricted^25^. Although the photon reassignment concept has mitigated this inherent low SNR condition^25^, the photon efficiency of SR confocal microscopy still needs to be improved for capturing fast long-term sub-organelle dynamics in multiple dimensions (5D in *xyz*-*color*-*time*), especially for its paralleled version, i.e., the spinning-disk confocal-based structured illumination microscopy (SD-SIM)^26^. Next, we experimentally evaluated our SN2N with ground-truth under a commercial SD-SIM system (Olympus SpinSR10 with an sCMOS camera, **Methods**). Compared to the diffraction-limited confocal mode (**Extended Data Fig. 4d**), the integrated optical reassignment module and 3.2 × second-stage magnification system of SpinSR10 bring heavily reduced SNR conditions. Using a commercial Argo-SIM slide (**Methods**), we measured the variance across the vertical straight line, and only our SN2N can draw the expected brightness profile with minimum fluctuations, even smaller than that of image under 100× exposure (**Fig. 2a**). Furthermore, the double-line structures are highlighted by SN2N from noise with the highest contrast (**Fig. 2a**, **Supplementary Video 2**). Based on several metrics including LRQ (line restoration quality, **Methods**)^12^ according to the known line structures, SSIM against the 100× exposure generated ground-truth, FRC, and STD from repetitive acquisitions, as well as visual inspection, we found SN2N successfully restored real-world collected images with superior stability and quality (**Fig. 2b-2e, Extended Data Fig. 4a, Supplementary Table 3**).

**Fig. 2.**
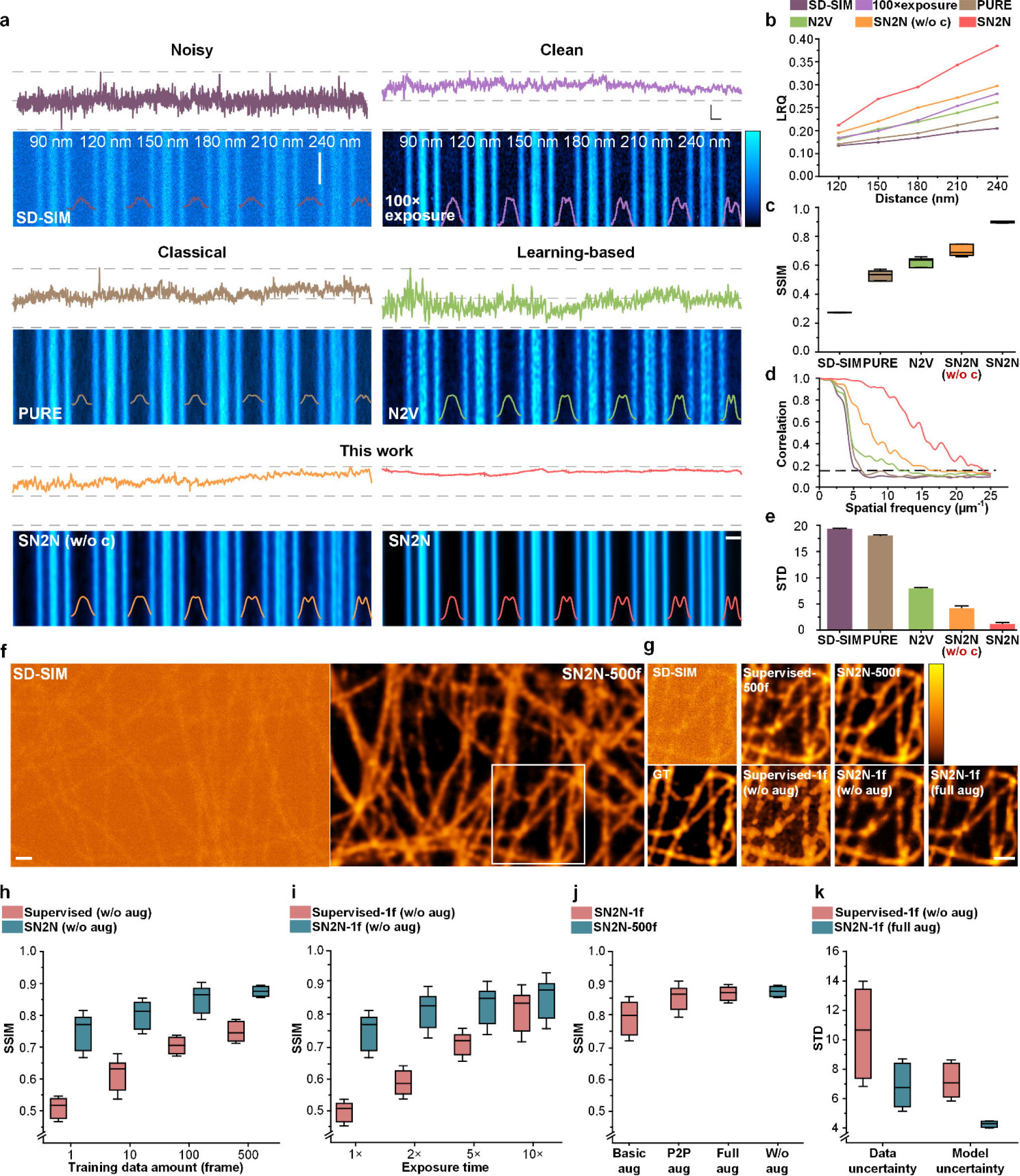
Systematical evaluations in SD-SIM experiments using known structures. **a**, Benchmarking of different denoising algorithms on the commercial Argo-SIM slide under SpinSR10 SD-SIM system. Representative images from different methods are presented below the corresponding intensity profiles indicated by the white line. Top row: Low SNR (left) and high SNR (with 100× exposure, right) SD-SIM images; middle row: PURE and N2V denoising results; bottom row: our SN2N without self-constrained loss (w/o c) and full SN2N denoising results. The intensity profiles of the corresponding double-line pairs (90 nm, 120 nm, 150 nm, 180 nm, 210 nm, and 240 nm distances) are displayed on their right. **b**, The line restoration quality (LRQ) values of images in **a**. **c**, SSIM values of different denoising methods (*n* = 10). The ground-truth image was created from the high SNR images by subtracting a constant background value and subsequently filtering a small Gaussian kernel. **d**, FRC analysis of the denoising images. The black dashed line denotes the 1/7 FRC threshold. **e,** The standard deviation (STD) of the denoising images predicted from ten repetitively collected noisy images. **f**, QD_525_ labeled microtubules in fixed COS-7 cells imaged by SD-SIM (left) and the corresponding SN2N results (right) trained with 500 frames (’SN2N-500f’). **g**, Enlarged regions enclosed by the white boxes in **f**. Top row: Raw image under SD-SIM, and supervised learning method (’Supervised-500f’) and SN2N results trained with 500 frames (’SN2N-500f’); bottom row: ground truth (GT) image, supervised learning (’Supervised-1f (w/o aug)’) and SN2N (’SN2N-1f (w/o aug)’) trained with 1 frame and no augmentation, and SN2N trained with 1 frame and full augmentation (’SN2N-1f (full aug)’). The ground truth image was created by the same procedure used in **b**. **h**-**j**, Average SSIM values of different training data amounts (*n* = 10) (**h**), different models trained by images under different exposure time (*n* = 10) (**i**), and different data augmentation strategies (*n* = 10) (**j**). In **h** and **i**, both the supervised learning method and SN2N were trained without data augmentation. In **j**, the basic augmentation (Basic aug) represents the random flip/rotate of training sets, and the full augmentation (Full aug) includes both our Patch2Patch augmentation (P2P aug) and basic augmentation. **k**, Data and model uncertainties quantified by the STD of ten independent frames and ten independently trained models, respectively. Centerline, medians; limits, 75%, and 25%; whiskers, maximum and minimum; error bars, s.e.m. Experiments were repeated ten times independently with similar results; scale bars, 1 µm (**a** and **f**), and 500 nm (**g**).

#### Experimental evaluation using biological samples under SD-SIM

To examine the broad applicability of SN2N, we applied it on microtubules in fixed COS-7 cells under another commercial SD-SIM system (GATACA Systems, Live-SR with an sCMOS camera, **Methods**) (**Fig. 2f**). The ultimate goal of unsupervised learning-to-denoise is to approach the similar performance of supervised learning without requiring the data pairs and large training set. In this case, we focused on comparing our SN2N against the supervised learning method in the denoising performance and data efficiency (**Fig. 2f**), in which the 500× exposure result is considered as the ground-truth. When only using one frame, the supervised learning method produces obscure structures (**Fig. 2g**), indicating an underfitting configuration, and the performance grows dramatically when increasing the data amount from 1 to 500 frames (**Fig. 2g**-**2h**). In contrast, we found the expansion of the data pool has little effect on the denoising performance of SN2N (**Fig. 2g**-**2h**), and the integration of our Patch2Patch (P2P) data augmentation (**Methods**) helps the one-frame-trained model to approach the performance of the 500-frame-trained model (**Fig. 2j**, **Supplementary Table 4**). Furthermore, by taking advantage of its intrinsic data efficiency and further reasonable data augmentation, SN2N exploits the full potential of available data, producing superior model and data uncertainties (**Fig. 2k**, **Supplementary Table 5**). Interestingly, we found that the one-frame-trained SN2N is less affected by the SNR degradations (**Fig. 2i**, **Supplementary Table 6**), prompting us to explore the trickier denoising task of multi-color live-cell SD-SIM data.

### RL-SN2N on SD-SIM unlocks fast long-term imaging across 5D

#### Multi-color live-cell SR imaging

The Richardson-Lucy (RL) deconvolution^40, 41^ has been routinely applied on SD-SIM to enhance the contrast and resolution, but it is prone to artifacts under live-cell imaging conditions (**Fig. 3b**). Thus, beyond denoising, we integrate the RL deconvolution with our SN2N (RL-SN2N, **Methods**) for simultaneously achieving the artifacts removal and resolution enhancement (**Fig. 3a-3b**). To avoid breaking the pixel-wise noise independency, the deconvolution is executed after the self-supervised data generation step during the training stage, and at the inference phase, the trained SN2N model is fed with the directly deconvolved images. We first validated the performance of RL-SN2N using the Argo-SIM slide, and its LRQ contrast substantially outperformed SN2N (**Extended Data Fig. 4b-4c**).

**Fig. 3.**
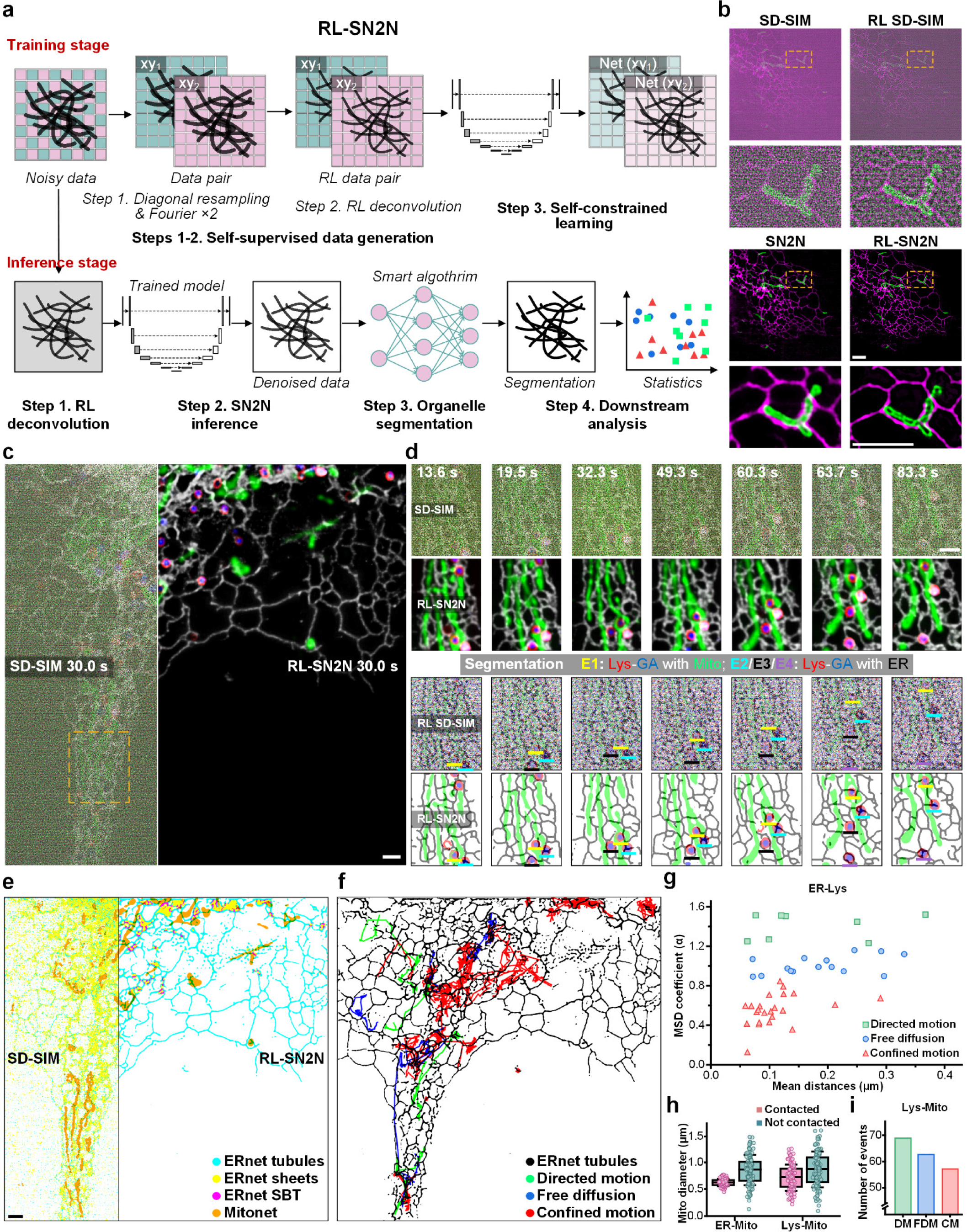
Multi-color live-cell SR imaging enabled by RL-SN2N on SD-SIM. **a**, Workflow of RL-SN2N. Training stage: Step 1, the data pairs are created with diagonal resampling followed by Fourier upsampling 2× (Fourier 2×); Step 2, the resulting data pairs are processed by RL deconvolution as RL-SN2N training set; Step 3, Self-constrained learning is executed using the generated RL data pairs. Inference stage: Step 1, the RL deconvolution is applied on the raw data; Step 2, the RL deconvolved data is fed into the trained RL-SN2N network and the outcomes are the final SR results; Step 3, the denoised SR images are input to the selected smart algorithms for segmentation; Step 4, Performing downstream analysis using the segmentation. **b**, A representative example for dual-color SR imaging of mitochondria (Mito, green) and ER (magenta) labeled with Tom20-mCherry and Sec61β-EGFP in live COS-7 cells under SD-SIM (top left), SD-SIM after RL deconvolution (top right, RL SD-SIM), SN2N result (bottom left), and RL-SN2N result (bottom right), alongside the enlarged region of yellow dashed box. **c**, A representative example for four-color imaging of Mito (green), ER (gray), lysosome (Lys, red), and Golgi apparatus (GA, blue) labeled with MitoTracker® Deep Red FM, Sec61β-EGFP, Lamp1-mCherry, and Golgi-BFP in live COS-7 cells under raw SD-SIM (right) and RL-SN2N (left)**. d**, The yellow dashed box in **c** is enlarged and shown at seven time points under different configurations. From top to bottom: Raw SD-SIM, RL-SN2N, RL SD-SIM segmentation (by Otsu hard threshold), and RL-SN2N segmentation (by Otsu hard threshold) results. The lines in different colors indicate different interaction events (E1-E4). **e**, Segmentation results of the Mito (orange, by Mitonet) and ER (by ERnet) under SD-SIM (left), and RL-SN2N (right). The ERnet segmentations contain tubules (cyan), sheets (yellow), and sheet-based tubules (magenta, SBT). **f**, Trajectories of Lys exhibiting directed motion (green), free diffusion motion (blue), and confined motion (red), and ER tubules (black). **g**, Distribution of the estimated *α* values of Lys versus their temporal average distances to ER tubules. **h**, Average Mito diameters (*n* = 100) (**Methods**). **i**, Average numbers of events surpassing the MOC threshold (>0.26) of Lys-Mito (*n* = 46). DM (green): directed motion; FDM (blue): free diffusion motion; CM (red): confined motion. Centerline, medians; limits, 75% and 25%; whiskers, maximum and minimum; error bars, s.e.m. Experiments were repeated ten times independently with similar results; scale bars, 2 µm (**c**-**e**), and 5 µm (**b**).

Also, we expect the resulting high-quality SR images can empower precise segmentation, facilitating automated analysis at a suborganelle-level precision from massive data (see the prototype in **Fig. 3a** and **Extended Data Fig. 5a**). The first representative case is dual-color imaging of endoplasmic reticulum (ER) and outer mitochondrial membrane (OMM) labeled live COS-7 cells (**Supplementary Video 3**). We can observe that our RL-SN2N effectively extracts the fluorescence signal from the noisy background with further enhanced contrast (**Extended Data Fig. 5b-5c**), hence many relative movements between OMM and ER can be dissected. Predictably, the direct hard threshold segmentation (**Methods**) on the low-SNR data produced highly broken structures and strong false positives. Our RL-SN2N minimized such noise-induced segmentation artifacts allowing inspection of fast mitochondrial fission events near the ER-Mito contact sites (arrows, **Extended Data Fig. 5c**).

Next, we extended application to four-color live-cell imaging of lysosomes (Lys), Golgi apparatus (GA), mitochondrial matrix protein (Mito), and ER (**Fig. 3c**, **Supplementary Video 4**). Consistently, our RL-SN2N provided plausible reconstructions for monitoring multiple organelle interactions from obscure captures. The increases in SNR and contrast were stable during time-lapse SR imaging (**Fig. 3d**), which led to distinct observations of Lys-GA (GA being surrounded by Lys) with Mito (Event 1, E1, yellow lines) and Lys-GA with ER (E2-E4, cyan, black, and purple lines) interactions. In these events, we found that Mito or ER settled in a circle could anchor onto a moving Lys-GA and be pulled out for spatial movements as it followed the trajectory of Lys-GA (segmentations in **Fig. 3d**)^42, 43^. Furthermore, we applied two recently developed learning-based segmentation methods, i.e., the ERnet^44^ and Mitonet^45^, on our ER and Mito data (**Methods**). Although better than the hard threshold segmentations, we found these well-trained networks were still influenced by the noise conditions (left in **Fig. 3e**, **Extended Data Fig. 5d-5g**). On the other hand, empowered by our RL-SN2N, ERnet could precisely draw the ER topology of tubules, sheets, and sheet-based tubules (SBT) (right, **Fig. 3e**). The trajectories of Lys and spatial masks of ER tubules (Methods, **Fig. 3f**) enabled us to examine the correlation between motions of lysosomes and the ER network. By calculating the MSD (mean square displacement) of Lys and their distances to ER, the Lys with confined motion behaviors were mostly located adjacent to ER^43^ (**Fig. 3g**, see also **Extended Data Fig. 5i, 5h, 5k**). Mitonet-generated masks allowed us to calculate the Mander’s overlap coefficient (MOC) of Lys with the nearest Mito for identifying potential functional sites of Lys-Mito contact sites^24^, i.e., MOC > 0.26 as a potential event (**Methods**, **Extended Data Fig. 5j**), in which the Lys with directed motion behaviors exhibited relatively larger number of events compared to the free diffusion and confined motions (**Fig. 3i**). We also quantified the mitochondrial diameters at the ER-Mito or Lys-Mito contact sites (**Methods**, **Fig. 3h**), reflecting the fact that the constricted loci in Mito preferentially associated with the ER-Mito contact^43^. Together, in the absence of tedious manual processes, our RL-SN2N on SD-SIM system simplified the dissection of the synergy of different organelles at a suborganelle scale (**Extended Data Fig. 5a**). Comparatively, N2V^19^, DeepCAD^22^, DeepSeMi^21^, and SRDTrans^33^ cannot provide valid denoising results from data under ultralow SNR conditions (**Supplementary Fig. 4**).

#### 3D-network and -deconvolution extension for volumetric SR imaging across 5D

With roots in the confocal microscopy, SD-SIM can perform 3D SR imaging with further enhanced axial contrast by the 3D deconvolution. To fully utilize the axial information, we extend the RL deconvolution and U-Net from 2D^35^ version to its 3D mode (**Fig. 4a**, **Extended Data Fig. 4f-4g**). By our RL-SN2N, 3D mitochondrial networks were visualized, and as expected, it revealed with tori cross-section of OMM genuinely, which were ambiguous in the raw SD-SIM results (**Fig. 4b**, **Supplementary Video 5**). Various types of OMM structures, i.e., from tubular to a series of different structures including fragments, small vesicles and spheroids, can be resolved by our RL-SN2N (**Fig. 4d**). The segmentation also helps to highlight the hollow structures of OMM networks, which are hardly distinguishable in the raw images (**Fig. 4c**). Furthermore, benefiting from SNR and contrast reinforcements, we can capture the 4D OMM network dynamics across hours (**Fig. 4e**, **Supplementary Fig. 5, Supplementary Video 5**). In our RL-SN2N results, the kiss-and-run events happening in 3D are correctly identified (arrows, **Fig. 4f**), which might be misinterpreted as the fissions in 2D slice under low SNR and contrast conditions. Finally, we extended our test to the challenging 5D SR imaging, revealing the dynamics of ER, Mito, and nucleus during the entire cell mitosis process in all three dimensions over long duration of 3 hours (**Fig. 4g**, **Supplementary Fig. 6, Supplementary Video 6**). Empowered by RL-SN2N, we clearly monitored the segregation process (**Fig. 4h**) and the interactions (**Fig. 4i**) of dense ER-Mito networks.

**Fig. 4.**
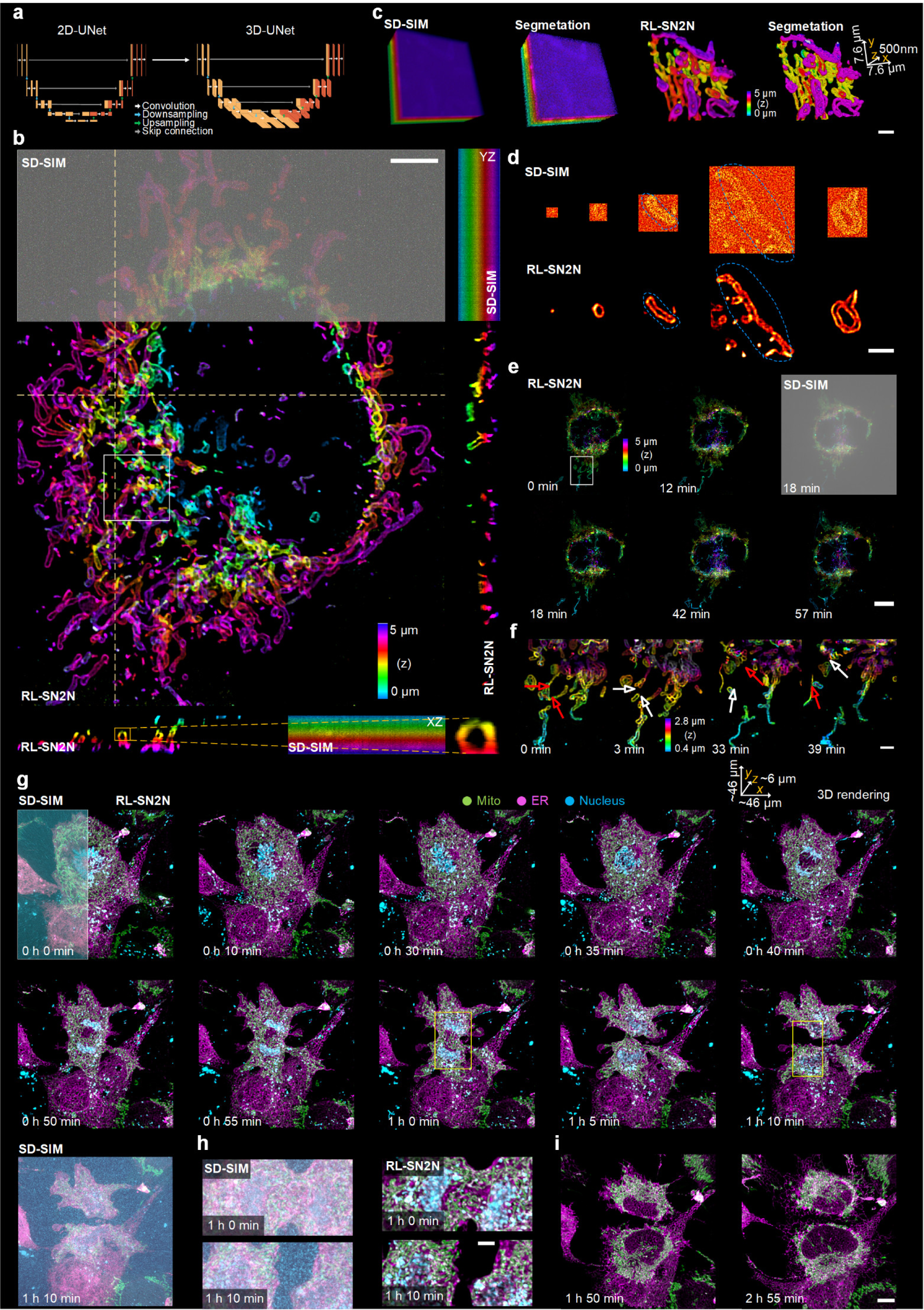
3D RL-SN2N on SD-SIM unlocks fast long-term imaging across 5D. **a**, The extension of 2D U-Net (top) to its 3D mode (bottom). **b**-**d**, 3D OMM network imaging of Tom20–mCherry labeled live COS-7 cells. **b**, Color-coded volumes of raw SD-SIM (top) and RL-SN2N (bottom). The color-coded axial views (*yz* and *xz* planes) indicated by the yellow dashed lines are provided alongside. Magnified view of yellow boxed regions on *xz* section is shown at the bottom left of the images. **d**, 3D rendering views of the white boxed region in **b** under raw SD-SIM (1^st^ column), segmentation of SD-SIM (2^nd^ column), SN2N (3^rd^ column), and segmentation of SN2N (4^th^ column). **c**, 2D slices under raw SD-SIM (top) and RL-SN2N (bottom). **e**-**f**, 4D imaging of OMM network (Tom20–mCherry) in live COS-7 cells. **e**, Representative color-coded volumes at six time points. **f**, Magnified views of the white boxed region in **e** under RL-SN2N at four time points. The red and white arrows indicate the mitochondrial fission and before fission, respectively. **g**, 5D imaging of mitochondria (green, mGold-Mito-N-7), ER (magenta, DsRed-ER), and nucleus (blue, SPY650-DNA) in live COS-7 cells. Representative 3D rendering views of the cell mitosis process at ten time points under RL-SN2N, except for the first (left part) and the last views being under raw SD-SIM. **h**, Magnified views from yellow boxes in **g** under raw SD-SIM (left) and RL-SN2N (right). **i**, Two representative time points of mitochondria (green) and ER (magenta) after mitosis. Experiments were repeated five times independently with similar results; scale bar, 5 µm (**b**, **i**), 1 µm (**c**, **d**, **h**), 10 µm (**e**), and 2 µm (**f**).

Additionally, according to the simulations, we found our approach is not sensitive to pixel size (**Extended Data Fig. 2a**). Thus, our success in SD-SIM equipped with an sCMOS camera (∼38 nm pixel) made us wonder whether our SN2N and RL-SN2N can be applied to diffraction-limited SD-confocal microscopy (∼65 nm pixel) or SD-SIM with an EMCCD camera (∼94 nm pixel) (**Methods**). On the Argo-SIM slice, our SN2N removed the readout noise of SD-confocal images (**Extended Data Fig. 4d-4e**). Likewise, by applying RL-SN2N (with further 2× upsampling) on two-color whole-cell volumes from the EMCCD SD-SIM, we successfully recorded two intermediate phases of cell mitosis, in which the 3D mitochondrion and nuclear protein distributions came into presence from background noise (**Extended Data Fig. 6**).

### Empowering long-term live-cell STED imaging

Similar to confocal SR microscopy, the increase in spatial resolution of STED microscopy from depletion brings dramatically decreased SNR^46^. Because the depletion is driven by de-excitation through stimulated emission, enlarging the depletion laser power will inherently decrease the emission brightness of fluorescence labels and cause adverse effects such as photobleaching and phototoxicity, preventing long-term monitoring of samples^46, 47^. Thus, the SN2N extraction of structures from few photons gives the possibility to tackle the contradiction for imaging live cells with both high spatial resolution and SNR. First, we systematically evaluated SN2N denoising performances under different depletion powers (**Fig. 5a**-**5b**, see also **Supplementary Fig. 7**), in which the data^47^ was collected from a custom STED setup incorporating a SPAD array detector with 5 × 5 elements for adaptive pixel-reassignment (APR) (**Methods**). By assembly of SPAD center pixel collected signals, the spatial resolution of STED is further improved but most photons would be dropped. From ref.^47^, the 5 × 5 APR STED results, or equivalently the STED image scanning microscopy (STED-ISM), can fully exploit the collectible photons to preserve SNR (**Fig. 5c**). Differently, without these collected photons from circumjacent 24 pixels, our SN2N effectively recovered the live microtubule structures from only center pixel collected photons, especially under a high depletion power (**Fig. 5b**). The two-peak draws of microtubule intersections, FRC curves, and FWHM (full width at half maximum) measurements also highlight the increment process of spatial resolution without sacrificing SNR after SN2N denoising (**Fig. 5d**-**5f**).

**Fig. 5.**
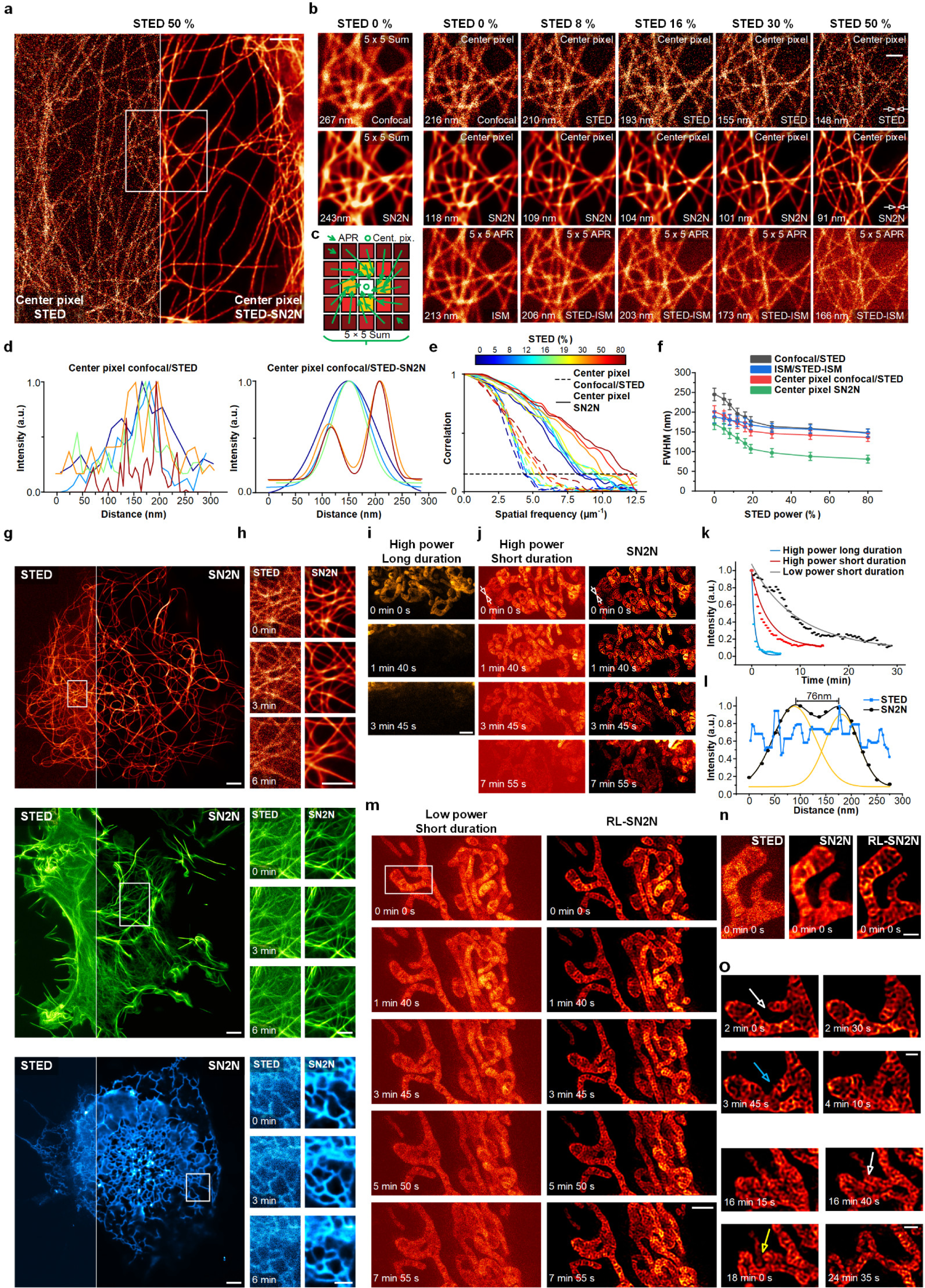
SN2N and RL-SN2N permit long-term live-cell STED imaging. **a**, A representative example of live HeLa cells labeled with SiR-tubulin under STED (center pixel, STED depletion laser power as 50%, left) and its SN2N result (STED-SN2N, right). Data from ref.^47^ (**Methods**). **b**, Magnified views of the white boxed regions in **a**. First column: Confocal image (5 × 5 sum, top) and its SN2N result (bottom); the other columns: STED images (center pixel, top) under different depletion laser powers (0%, 8%, 16%, 30%, and 50% from left to right), its SN2N results (middle), and its ISM results (5 × 5 adaptive pixel reassignment, APR, STED-ISM, bottom). FRC-measured resolution values are marked at the left bottom. Notably, STED with a depletion laser power as 0% is equivalent to the confocal configuration. **c**, A sketch of different strategies for pixel assembly, including center pixel (‘Cent. pix.’), 5 × 5 sum, and APR. **d**, Fluorescence intensity profiles along the white arrow in **b** of confocal/STED (center pixel, left) and their SN2N results (right) under depletion laser powers. Different colors indicate different depletion laser powers. **e**, FRC analysis of the confocal/STED images (center pixel) before and after SN2N denoising under depletion laser powers. The black dashed line denotes the 1/7 FRC threshold. **f**, Average FWHM values of 5 × 5 sum confocal/STED, ISM/STED-ISM, center pixel confocal/STED, and center pixel SN2N results (*n* = 6). **g**, STED snapshots of live COS-7 cells labeled with SiR-Tubulin (top), Lifeact-EGFP (middle), and Sec61β–EGFP (bottom) under a commercial STED microscope (Leica, **Methods**). Left: the original STED images. Right: denoised images using SN2N. **h**, Raw STED frames (left) and SN2N (right) denoised counterparts of magnified views of the white-boxed regions in **g**. Three representative time points are provided from top to bottom. **i**, Three representative frames of PKMO labeled live COS-7 cells under another commercial STED microscopy (Abberior) at high depletion power (86%) and long duration time (100 μs per pixel). **j**, Examples STED raw images (left) and their SN2N results (right) at high depletion power (86%) and short duration time (10 μs per pixel). **k**, Photobleaching analysis of STED images used in **i**, **j**, and **m**, quantifying the normalized signal in each case. **l**, Fluorescence profiles along the white arrow in **j** imaged by STED and SN2N. **j**, Examples STED raw images (left) and their SN2N results (right) at high depletion power (86%) and short duration time (10 μs per pixel). **m**-**p**, Long-term STED imaging of PKMO labeled mitochondrial cristae in live COS-7 cells at high depletion power (41%) and short duration time (10 μs per pixel). **m**, Representative frames of STED (left) and RL-SN2N (right) results. **n**, Magnified views of the white boxed regions in **m** by raw STED, SN2N, and RL-SN2N. **o**, Representative montages of the mitochondrial fusion and fission events. The yellow and blue arrows highlight the regions of mitochondrial fusion and fission, respectively, while the white arrows indicate the moments before events. Centerline, medians; limits, 75% and 25%; whiskers, maximum and minimum; error bars, s.e.m. Experiments were repeated five times independently with similar results; scale bar, 2 µm (**a**), 1 µm (**b**, **g**, **h**), 500 nm (**i**, **m**-**o**).

It is straightforward to apply our SN2N on a commercial STED system (Leica, TCS SP8 STED 3X, **Methods**) for time-lapse imaging of microtubules-, actin-, and ER-labeled COS-7 cells (**Fig. 5g**-**5h**, **Supplementary Video 7**), routinely enabling high-quality live-cell SR imaging. Beyond that, we also recorded the long-term dynamics of mitochondrial cristae (PK Mito Orange, PKMO^48^ labeled) using another commercial STED system (Abberior Instruments, STEDYCON, **Methods**) under various imaging conditions (**Supplementary Video 7**). To acquire high-quality STED images *in situ*, the high depletion power and long pixel duration time were applied and hereby instantly extinguished the fluorescence signal before 5 frames of recording (**Fig. 5i**). The high depletion power with short pixel duration time could delay the bleaching effects without loss of spatial resolution but create noisy captures, in which our SN2N effectively restored this SNR degradation (**Fig. 5i**-**5k**). The measured cristae-to-cristae distance reflects the achievable resolution of ∼76 nm (**Fig. 5l**). Further turning down the depletion laser power will offer significantly less susceptible to photobleaching but short of spatial resolution maximization (**Fig. 5k**, **5m**). Fortunately, our RL-SN2N can lift the dropped resolution in the absence of amplified photobleaching (**Fig. 5m-5n**). Short exposure and low illumination energy facilitated us to record the perplexing locomotion of cristae during mitochondrial fusion (**Fig. 5o**) and fission (**Fig. 5p**) over half an hour. In contrast, conventional STED produced uninterpretable SR images under the same conditions (**Fig. 5m**).

### Improving reconstruction efficiency of SOFI

Super-resolution optical fluctuation imaging (SOFI)^30^ can routinely break the diffraction limit by exploiting the natural temporal fluctuations of fluorescence emissions under optical systems in their native states. However, the statistical uncertainty of reconstructions from short sequences may dramatically affect image continuity and homogeneity, which leads it generally requiring hundreds of raw images (∼1,000 frames for 2^nd^ order) to preserve structural integrity^31^. To increase its reconstruction efficiency, we integrated our SN2N solution into SOFI reconstruction pipeline (**Fig. 6a**). Specifically, the raw image sequence is calculated by *n*^th^ order cumulant (core SOFI), and the resulting image is followed by the RL-SN2N procedure (**Methods**). Using SIM as reference, the 2^nd^ order SOFI-SN2N is first validated on a wide-field microscope (**Methods**). We found that the strong snowflake artifacts in conventional 20-frame SOFI were effectively eliminated, and the original microtubule structures were highlighted with high axial contrast. (**Fig. 6b**, **Extended Data Fig. 7a, Supplementary Video 8**) The two-peak analyses, FRC metrics, and FWHM measurements all demonstrate the massively improved temporal resolvability and effectively increased spatial resolution of SOFI-SN2N (**Fig. 6b**-**6d**). Next, we extended our SOFI-SN2N to different orders of cumulants executed on a commercial SD-confocal microscope (**Fig. 6b**-**6d**, **Methods**). Consistently, both visual examination and FRC analysis exhibit that the 2^nd^ order SOFI-SN2N can produce artifacts-free results from only 20 frames (**Fig. 6e**-**6g**). On the other hand, the increase of resolution from higher order cumulant brings a cost of more frames needed, and our SOFI-SN2N enables efficient 3^rd^ order and 4^th^ order SOFI reconstructions from 50 frames and 100 frames, respectively (**Fig. 6f**, **Extended Data Fig. 7b-7e**). The FWHM and FRC measured resolution values (**Fig. 6h**) also evince the spatial resolution enhancement. Finally, the recording of OMM dynamics during mitochondrial fusion during ten minutes gives us a glance at live-cell SN2N-SOFI SR imaging (**Fig. 6i-6j**, **Supplementary Video 8**).

**Fig. 6.**
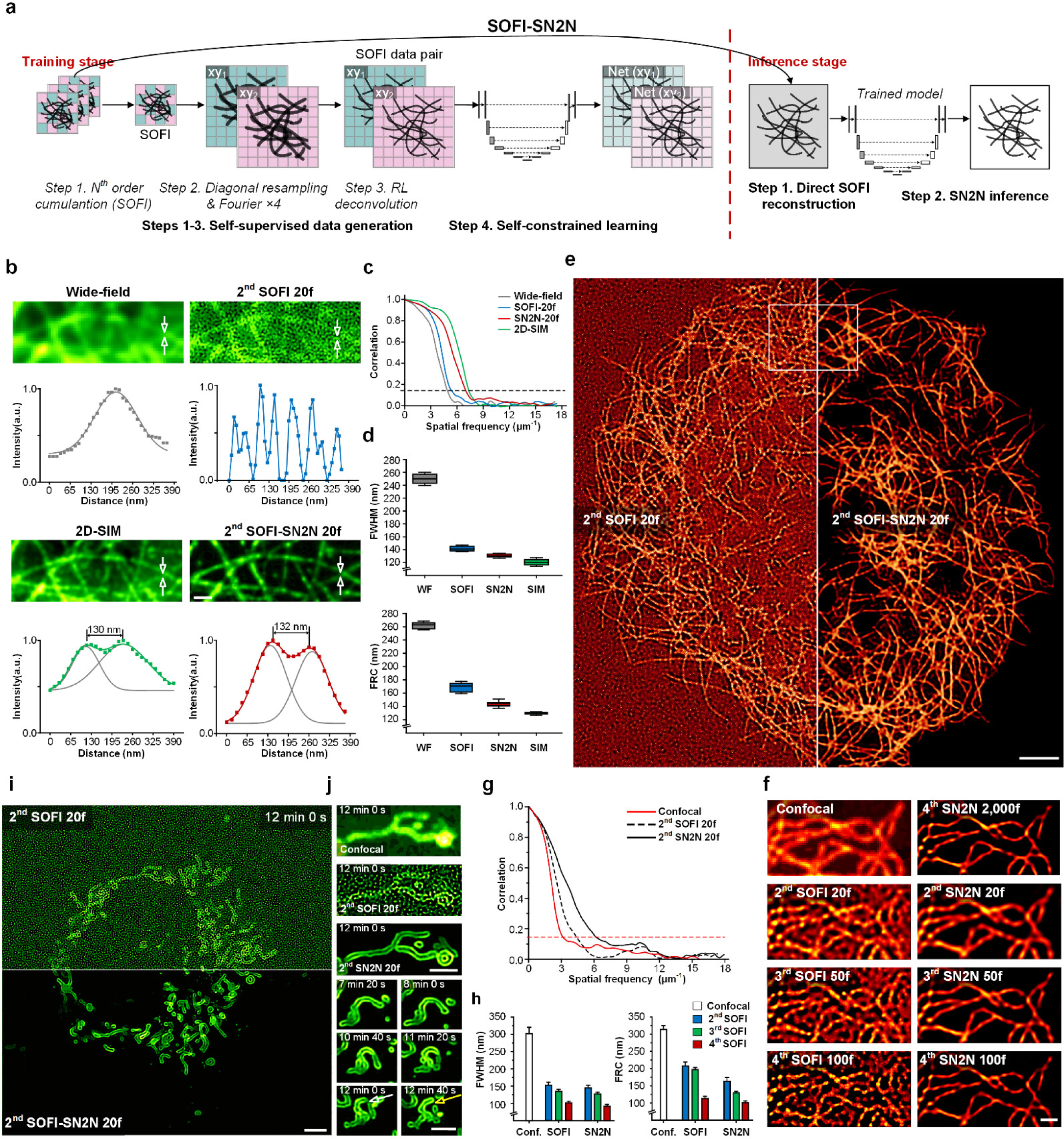
Integration of SN2N and SOFI massively improves the SR reconstruction efficiency. **a**, Workflow of SOFI-SN2N. Training stage (red dashed box): Step 1, the acquired image sequence is calculated by *n*^th^ order SOFI; Step 2, the SOFI data pair is created by diagonal resampling followed by Fourier upsampling 4× on the raw SOFI image; Step 3, the SOFI data pair is then processed by a RL deconvolution followed by a linearization operation; Step 4, the self-constrained learning is executed using the generated SOFI data pairs. Inference stage: Step 1, generating the SOFI image. The acquired image sequence is calculated by *n*^th^ order SOFI, and the resulting image is processed by Fourier upsampling 2× followed by an RL deconvolution and a linearization operation; Step 2, This resulting SR image is then fed into the trained SN2N model for high-quality reconstruction. **b**, Cross-validation of SOFI-SN2N. Snapshots of microtubules in a COS-7 cell labeled with QD_525_ under wide-field microscopy (top left), 2D-SIM (bottom left), 2^nd^ order SOFI using 20 frames (2^nd^ SOFI 20f, top right) and its SN2N result (2^nd^ SOFI-SN2N 20f, bottom right). The Intensity profiles and multiple Gaussian fitting indicated by the white arrows of the corresponding modalities are provided at their bottom. The numbers indicate the distance between peaks. **c**, FRC analysis of the images in **b**. The black dashed line denotes the 1/7 FRC threshold. **d**, Average FWHM (top) and FRC (bottom) values (*n* = 5). WF: wide-field. **e**, Examples of microtubules in a COS-7 cell labeled with QD_525_ under a commercial SD-confocal microscopy reconstructed by 2^nd^ order SOFI using 20 frames and its SN2N result (right). **f**, Zoomed views from the white box in **e**. First row: confocal image (left) and 4^th^ order SOFI using 2,000 frames denoised by SN2N (4^th^ SOFI-SN2N 2,000f, right); the other rows, from top to bottom: 2^nd^, 3^rd^, and 4^th^ orders SOFI reconstructions (left) using 20, 50, 100 frames, respectively, and their SN2N results (right). **g**, FRC analysis of raw SD-confocal image, 2^nd^ SOFI 20f reconstruction, and its SN2N result. The red dashed line denotes the 1/7 FRC threshold. **h**, Average FWHM (left) and FRC (right) values of SD-confocal images, SOFI images, and their SN2N results (*n* = 5). **i**, A representative live COS-7 cell labeled with Skylan-S-TOM20 imaged by 2^nd^ SOFI-20f of SD-confocal microscopy and its SN2N result (bottom). **j**, Zoomed views of OMM structures. Top three rows, from top to bottom: magnified views of the white boxed region in **i** under SD-confocal microscopy, 2^nd^ SOFI 20f reconstruction, and 2^nd^ SOFI-SN2N 20f result; bottom three rows: montages of a representative mitochondrial fission event. The yellow and white arrows highlight the mitochondrial fission and before fission, respectively. Centerline, medians; limits, 75% and 25%; whiskers, maximum and minimum; error bars, s.e.m. Experiments were repeated three times independently with similar results; scale bars, 2 µm (**b**, **i**, **j**); 5 µm (**e**) and 1 µm (**f**).

In fixed-cell experiments, the temporal sampling^22, 24^ (first and second 20 frames for SOFI reconstruction) can be applied directly (**Extended Data Fig. 7f**). Interestingly, due to the inconsonant temporal fluctuation behaviors, the temporal sampling leads to statistical differences between the adjacent frames and produces strong artifacts. In contrast, our spatial sampling is impressed with sturdily extracting the microtubule structures for requiring no temporal consistency (**Extended Data Fig. 7f**).

### SN2N on expanded samples

In addition to use of fluorescence fluctuations, the expansion microscopy (ExM)^29^ is another system-agnostic SR modality by artificially enlarging the size of samples to break the diffraction-limit. However, considering the determined number of fluorophores, this space extension of samples will result in the geometrically decreased SNR according to the expansion times (**Extended Data Fig. 7g-7k**). Under a wide-field microscope (**Methods**), we offered the ExM-SN2N enabling high-quality SR imaging of ∼110 nm and ∼67 nm resolutions for 2× and 4× expansions (**Extended Data Fig. 7g-7i**), respectively. Under different noise levels, the ExM results after SN2N denoising exhibited significantly improved signal-to-background ratios (SBRs). Similarly, the SN2N outcomes of the 4.5 times-expanded cells yielded (**Extended Data Fig. 7-7k**) noise-eliminated results, revealing the complex ER tubule structures.

### Artifacts-removal for live-cell SIM

Although structured illumination microscopy (SIM) is recognized to have a higher photon efficiency than other SR modalities, it still requires an adequate SNR for each raw image to prevent random reconstruction artifacts^10, 12, 24^. Therefore, to minimize the artifacts, we included the SN2N engine into SIM reconstruction (’raw resampling’, **Extended Data Fig. 8a**). The generated twin 9-raw-image were individually reconstructed by the HiFi-SIM^49^ procedure (high-fidelity SIM, **Methods**) to create the training sets, and at inference stage, the input is SIM image by the original 9 raw images. First, we systematically evaluated the performance of our SIM-SN2N using the BioSR open-sourced dataset^50^ under high (H), medium (M), and low (L) SNR levels. Comparing to the ground-truth ultrahigh SNR reconstructions, our SIM-SN2N effectively disentangled the real features from artifacts and produced stable SSIM values for various conditions and samples (**Extended Data Fig. 8b-8g Video 9**). Then, the 2D-SIM imaging of mitochondrial cristae exhibited strong background artifacts, which was further amplified as the emission fluorescence progressively decreasing (**Extended Data Fig. 8h-8i**). The suppression of artifacts in SIM-SN2N supports its superior performance in obtaining high-fidelity SR images from raw images of low SNR (**Extended Data Fig. 8j, Supplementary Video 9**).

In particular, when meeting ultrafast SIM imaging under ultralow SNR conditions, the resampled dataset might encounter reconstruction failures, in which the parameter estimation is highly unstable. On the other hand, the frequency-component reassembly step in SIM reconstruction is usually accompanied by an artificial 2× upsampling, which breaks the pixel independency, and this prevents the direct application of SN2N on SIM images. To meet this challenge, we conducted a new strategy of spatial sampling by defining 3 × 3 pixels as one unit and a new average calculation of 4 directions followed by the Fourier 3× operation (’SIM resampling’, **Extended Data Fig. 8k**). This one-pixel interval enabled the direct artifacts-removal on the SIM reconstructed images only with a slight drop of SSIM metrics (**Extended Data Fig. 8l-8q**). Tested on the ultrafast (188 Hz, ∼0.6 ms exposure per raw frame) TIRF-SIM experiments, the ‘raw sampling’ failed to offer eligible SIM reconstructions, and in contrast, the ‘SIM resampling’ provided high-quality images along 6,800 consecutive SR frames (**Extended Data Fig. 8r-8t**).

## DISCUSSION

In the N2N ecosystem, theoretically, the infinite data pairs are essential to approach the supervised learning methods’ performance, because of the need for averaging training sets to remove the zero-mean noise. Using twin images from our self-supervised data generation, the developed self-constrained learning process further generalizes this N2N concept to remove noise with randomness, and also relaxes the need for infinite data amount^17^. As a result, our SN2N is competitive with supervised learning methods but overcoming the need for large dataset and clean ground-truth, in which a single noisy frame is feasible for training. We applied SN2N to the photoelectric detector directly captured data, including two commercial SD-SIM systems with two different types of cameras, one custom-built and two commercial STED microscopes, one ExM under a wide-field microscope, and one commercial SD-confocal microscope, demonstrating its extensive application value and superior performance. Besides, we also integrated SN2N into the prevailing SR reconstructions, including RL deconvolution, SOFI, and SIM for artifacts removal, enabling efficient reconstructions from limited photons by one-to-two orders of magnitude.

A concern of learning-based recovery is that the spatially denoised features might distort the temporal signals in a nonlinear manner. Recording cells labeled with cytosolic Ca^2+^ indicators^51^, we found SN2N acted without nonlinearly perturbing the amplitudes of different Ca^2+^ transients, indicating that the SN2N denoising is quantitatively accurate (**Extended Data Fig. 9**). Although SN2N was shown to work well overall, it would be appropriate to discuss limitations observed in the current version. Notably, the internal mechanism of learning-based denoising methods is substantially different from the classical ones. In an abstract sense, SN2N is learning to extract structures from noisy input according to the training set, while the numerical algorithms usually intend to remove the noise. Thus, the extraction of structures from models trained by higher SNR data would exhibit strong background artifacts caused by the misleading information learned between noise and structures (**Extended Data Fig. 10a-10b**), and the outcomes from models trained by lower SNR data are free from this failure (**Extended Data Fig. 10d-10f**). We could moderate this issue by the iterative execution of the SN2N model (SN2N^2^) to shrink the gap between training and test sets (**Extended Data Fig. 10f**). In addition, this network extraction is also influenced by the structural scale of input images, and the up/down sampling operation should be applied to match the pixel size before SN2N inference (**Extended Data Fig. 10g-10k**), otherwise the network would produce erroneous results (middle panel, **Extended Data Fig. 10h, 10i**).

Our SN2N is a model-agnostic solution. The 2D/3D U-Nets used in this work are simple end-to-end baselines, and the extensions to other advanced networks are straightforward, such as the routinely used residual/dense blocks^52^, adversarial training strategy^53^, and transformer-based solutions^54^. Furthermore, the incorporations of SN2N with network-based deconvolution and SIM are expected to enable stable and efficient SR reconstructions for the real-time potential. Finally, the loss realizations of our SN2N have many variants, and the distance calculation by SSIM or in the Fourier domain could lead to additional improvements.

Random noise is an ineluctable obstacle in fluorescence microscopy, especially for the live-cell SR recording. Our SN2N and its extensions provide powerful solutions for routine 2D∼5D imaging of suborganelle dynamics at ultra-high spatiotemporal resolution and high-fidelity for long durations. Overall, we anticipate that the elimination of noise could benefit precise structure segmentation and facilitate automatic multi-parameter analysis, establishing a panoramic view of the organelle interaction systems^44, 45^.

## METHODS

### SN2N framework

#### SN2N core

In principle, N2N does not requires clean target to train a denoising model approaching supervised learning performance. However, this is achieved by assuming that the noise is zero-mean and the training set contains infinite noisy data pairs sharing identical contents with independent noise components. First, to remove the need for data pairs, we designed a diagonal resampling strategy to generate such twin images from one single frame using the inherent spatial redundancy of SR images. In specific, we assembled every 4 pixels (as follows) in an SR image as one unit:

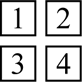

and then we applied a special diagonal binning operation on these four pixels to create two new pixels for N2N data pair. We averaged the two diagonal pixels, 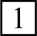 and 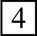, as one new pixel; and the other two diagonal pixels, 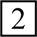 and 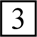, as the other new pixel. Throughout the input image, two 2× subsampled images were created. Second, to rescale back to their original image size, we adapted a Fourier interpolation method to upsample the resulting image pair by two times. Thus, we transformed the two subsampled images into the Fourier domain and applied zero-padding to extend the borders in the Fourier space by half of the original sizes in each direction. Then, the inverse Fourier transformed image was twice the size of the input image, identical to the original SR image. Additionally, to prevent boundary artifacts, we extended the image (in spatial domain) with mirror-symmetric half-copies before the Fourier padding, and the resulting image was Fourier upsampled and then cropped back to the original size.

Third, originally in N2N, the infinite training set can execute average of noise to its true zero-mean result. However, in case of limited data for training, there would be inevitable differences between these means across the different realizations of noise. Thus, the insufficient training set will introduce large predictive uncertainty, inherently resulting in limited denoising performance. To minimize this uncertainty, we designed a self-constrained learning process with considering the identical underlying content from the noisy data pair. Specifically, the noisy data pair were successively and individually fed into the network, and the two resulting predictions were calculated by the following loss function to execute the training stage.

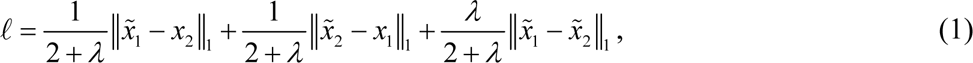

where 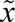 represents the network outcome of the input *x*, and ‖ ‖ norm. The first two terms are the conventional N2N loss. Because the corresponding two denoised results should have no differences, we included the last term as our self-constrained loss with constraint weight *λ* to enforce the consistency of the predictions.

#### Patch2Patch data augmentation

We developed a data augmentation pipeline, namely Patch2Patch, employing multidimensional random patch transformations to increase the data amount, which is adaptable to both 2D (*xy*) and 3D (*xyz*) datasets (**Extended Data Fig. 1b**). Within the Patch2Patch framework, there are three modes for augmentation along the temporal axis, in a single image, and between different experiments. These designs will effectively increase variations of noise realizations and structural features without altering the intrinsic characteristics of data. Mode 1: For image/volume dataset with additional temporal dimension (*xy*-*t*/*xyz*-*t*), Patch2Patch defaults to create augmentation along the temporal dimension. The randomly chosen patches at time point A (Stack A, Image A) are exchanged with the patches from randomly selected time point B (Stack A, Image B) at the same locations. Mode 2: For the augmentation of a single image/volume, two distinct patches at different positions are randomly swapped. Mode 3: For augmentation between different experiments, the randomly chosen patches at experiment A (Image A) are exchanged with the patches from randomly selected experiment B (Image B) at randomly picked locations. To further increase the data amounts, we also randomly applied one of six geometric transformations for each pair of patches: no transformation, horizontal flip, vertical flip, 90° rotation to the left, 90° rotation to the right, and 180° rotation.

##### Network architecture

Because our SN2N can be applied on any DNNs, we simply adopted the widely recognized U-Net^35^ architecture to showcase the strengths of SN2N (**Extended Data Fig. 1a**). The 2D U-Net architecture consists of a 2D encoder module (contracting path), a 2D decoder module (expanding path), and four skip connections bridging the encoder and decoder. Both the 2D encoder and decoder modules are structured into four distinct blocks. Each encoder block is equipped with two 3 × 3 convolutional layers, followed by a leaky rectified linear unit (LeakyReLU) and a 2 × 2 max pooling operation with a stride of two in both dimensions. Conversely, each decoder block contains two 3 × 3 convolutional layers, followed by a LeakyReLU and a 2D bilinear interpolation. Batch normalization (BN)^55^ is integrated after each convolutional layer. The skip connections serve to concatenate low-level and high-level feature maps, enhancing the preservation of spatial information.

For denoising tasks involving three-dimension datasets (*xyz*), we directly shifted the U-Net from the 2D version to its 3D extension to better leverage the axial spatial information. With all its internal operations tailored to a three-dimensional framework, the 2D U-Net was transformed into the 3D U-Net. In this work, we changed the convolution operations from 3 × 3 to 3 × 3 × 3, the max pooling from 2 × 2 to 2 × 2 × 2, and the interpolation operations from 2D bilinear to 3D trilinear interpolation.

#### Learning and inference processes

Regarding the training data generation, the percentile image normalization is first applied before training to remove the baseline background and moderate the large intensity gap between the bright and dim fluorescence signals, which is defined for an image *x* as:

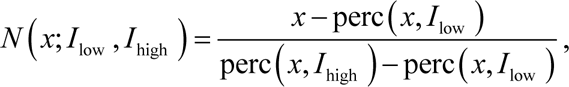

where *perc*(*x*, *I*) is the *I*-th percentile of all pixel values of *x*, and *I*_low_ and *I*_high_ represent the lowest and highest values, respectively. Notably, for some data under ultra-low SNR conditions, a wavelet-based background subtraction^12^ was executed before this percentile normalization.

For small dataset, we first executed the Patch2Patch pre-augmentation pipeline on the raw dataset to enlarge the training set (Step. 1, **Extended Data Fig. 1**). Then, a sliding window approach was employed to generate small patches (128 × 128 or 128 × 128 × 16 tiles by default) suitable as network input for training (Step. 1, **Extended Data Fig. 1**). The interval of the sliding window was customizable to adjust different image processing requirements (64 pixels by default). During this step, a background patch rejection was performed on the fly, in which the patches with averaged intensity twice lower than that of the entire image/volume were filtered. For these small patches, the spatial diagonal resampling strategy was applied to produce pairs of twice smaller sub-images, each sharing identical content but different noise realizations (Step. 2, **Extended Data Fig. 1**). After that, the Fourier upsampling was used to create paired SN2N data to match the original structural scale. Additionally, the conventional data augmentation strategies such as rotation and flipping could be included to further increase the network generalization (Step. 3, **Extended Data Fig. 1**). Finally, the self-constrained learning process utilized the basic U-Net network and selected either the 2D U-Net or 3D U-Net based on the input data dimensions (Step. 4, **Extended Data Fig. 1**).

We set the constraint weight *λ* as 1 routinely in most of our experiments. The models were optimized utilizing the Adam optimizer^56^ with a learning rate of 2 × 10^−4^, and the first-moment and second-moment estimates were regulated with exponential decay rates of 0.5 and 0.999, respectively. For every training iteration, the batch size is set to the maximum number reaching the GPU (graphics processing unit) memory limit, in which the training stage was conducted using NVIDIA GeForce RTX 4090 GPUs. Finally, in the inference process, the raw data after the percentile normalization is directly fed into the trained SN2N network for predictive analytics. For input data sizes that exceed the memory limit (most of the volumetric data), the inputs are automatically cropped into dozens of subvolumes and fed into the SN2N network, and the predictions are stitched back together to generate the final denoised results.

### Integrations of the SN2N framework with SR reconstructions

#### SN2N-enhanced RL deconvolution

We have integrated RL deconvolution^40, 41^ with our SN2N framework, namely RL-SN2N (**Fig. 3a**). This approach leverages the contrast and resolution enhancement capabilities of RL deconvolution while mitigating the artifacts that arise from deconvolution under low SNR conditions. To preserve the stochastic independence of noise at the pixel level, we perform RL deconvolution after the initial self-supervised data generation. We used the accelerated RL deconvolution:

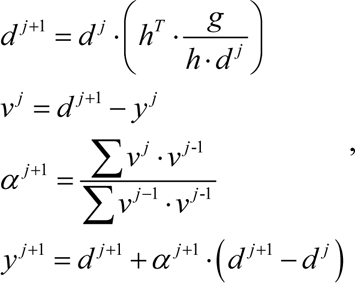

where *y ^n^*^+1^ is the image after *n*+1 iterations; *g* is the input image; and *h* is the PSF. The adaptive acceleration factor *α* was introduced by Andrews et al.^57^, representing the length of an iteration step, which can be estimated directly from experimental results. The RL deconvolution was executed by the corresponding theoretically calculated 2D/3D PSFs by the Gaussian kernel approximation, and the iterations were usually selected by 10∼15 times. During the inference phase, the trained RL-SN2N model is fed with the directly deconvolved image/volume.

#### SN2N-enhanced SOFI

We integrated our SN2N solution into the SOFI reconstruction pipeline (SOFI-SN2N, **Fig. 6a**) to increase its imaging efficiency^30^. After the acquisition of the image sequence, we calculated the *n*^th^ order of auto-correlation cumulant, i.e., the 2^nd^, 3^rd^, and 4^th^ order SOFI, on each pixel along time. Then, we performed the self-supervised data generation and RL deconvolution successively. To match the improved spatial resolution, in SOFI-SN2N, we applied 4× Fourier upsampling on the diagonal resampled data pair rather than the routinely used 2×. Finally, after RL deconvolution, we used an intensity linearization step by taking the *n*-th root directly to minimize the nonlinear effects of SOFI. In the inference stage, the trained SOFI-SN2N model is fed with the SR image reconstructed by the *n*^th^ order SOFI followed by 2× Fourier interpolation before RL deconvolution and intensity linearization.

#### SN2N-enhanced SIM

We designed two strategies for artifacts removal of SR-SIM images. Mostly, we first performed the self-supervised data generation to every modulated raw frame (9 frames) for creating the twin SIM sequences. After that, we individually reconstructed the paired SR-SIM images with the HiFi-SIM^49^ pipeline. Finally, these two resulting SIM images were considered as the training set of our SIM-SN2N. In the test stage, the ordinary SIM reconstruction was directly inputted into the SIM-SN2N network.

Beyond that, the self-supervised data generation might influence the parameter estimation, especially under ultra-low SNR conditions for ultra-fast imaging, leading to failed reconstruction. On the other hand, the SIM reconstruction brings the inherent artificial upsampling (2-fold), and hence the adjacent pixels are highly correlated. To create independent data pairs from SIM reconstruction, we performed the resampling step within 3 × 3 pixels (**Extended Data Fig. 8**). Specifically, we assembled every 9 pixels as one unit:

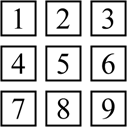

and then we applied a special binning operation on these 9 pixels to create two new pixels for the N2N data pair. We averaged the four corner pixels, 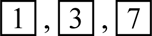, and 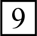, as one new pixel; and the other four intermediate pixels, 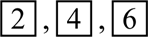, and 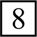, as the other new pixel. Throughout the input image, two 3× subsampled images were created. Followed by a 3× Fourier upsampling, we finalized the required data pair directly from an SR-SIM image.

### SN2N-empowered automated subcellular segmentation and tracking

#### Subcellular segmentation

We employed the Otsu^58^ method to automatically determine the hard thresholds for identifying the corresponding organelle features in images. Additionally, to achieve more precise segmentation, we also utilized pre-trained models from several learning-based approaches, e.g., ERnet^44^ for ER structures and Mitonet^45^ for Mito shapes. Following the segmentation of ER, we eliminated isolated pixels in the binarized masks and then extracted the skeleton structures for the network topology construction, in which the nodes represent intersections in the skeleton graph, and edges represent connections between these nodes. After Mito segmentation, we computed the connected domains within the binary masks and identified the skeletons and key points. These key points were subsequently categorized into junctions or end points based on their respective topological positions (**Extended Data Fig. 5d**).

#### Tracking of Lys

Tracking of Lys was performed using TrackMate 7.11.1^59^. To characterize their dynamic behaviors, we computed the mean square displacement (MSD) for the trajectories across all timepoints^60^. The calculated MSD curves are approximated with a power-law function^60^:

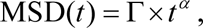

where *t* represents the time interval, Γ is the proportionality factor that relates to both particle motion dynamics and the physical properties of the system, and *α* characterizes the different modes of particle movements^61^. Then, a logarithmic transformation is applied to the MSD formula, followed by a linear regression to estimate *α*:

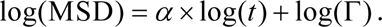

In our analysis, the Lys movements were classified into confined motion (*α* < 0.85), free diffusion (0.85 ≤ *α* ≤ 1.2), and directed movement (*α* > 1.2)^61^.

#### Mito diameter

To estimate the Mito diameter at a selected point, we drew the tangent line of the nearest Mito contour. Using the perpendicular line of this tangent line, we obtained the other intersected point to the opposite Mito contour. Finally, we approximated the diameter by measuring the distance between the initially selected point and this intersection point^62^.

#### ER-Lys distance

We identified the edges of ER tubules from the ERnet-generated masks, and the distances were calculated based on the centroid coordinates of Lys and the most adjacent ER edges. Then, the final ER-Lys distances were quantified by averaging the distances across the entire trajectory.

#### ER-Mito distance

The ER-Mito contact level was quantified by the minimum distance from each Mito to the edges of ER tubules, in which the distances as 0 were distributed as contacted and larger than 0 as not-contacted. The Mito diameter was estimated at the point with the closest ER-Mito distance.

#### Lys-Mito interactions

We calculated the Mander’s overlap coefficient (MOC) to examine the Lys-Mito interactions. The masks of Mito and Lys were convolved with the PSF of the SD-SIM system, and the MOC values were calculated from the resulting masks at each timepoint. Subsequently, we calculated the ratio of the mean value and standard deviation of the MOC curve and obtained an empirical threshold of 0.26, in which the associated MOC values larger than this threshold were identified as functional sites in specific instances of Lys-Mito contact^24^ (**Extended Data Fig. 5j**). With MOC values below the threshold, intersection points on the Mito edge were identified from the connecting line of Lys and Mito centroids. We then used it as the selected point for the Mito diameter estimation. For events of MOC values higher than the threshold, we drew the perpendicular line of the connecting line from the two intersection points between the edges of Lys and Mito, and the intersection of this perpendicular line and the Mito edge was picked as the selected point for the Mito diameter estimation.

### Performance metrics

#### Simulations of microtubule filaments

To perform benchmarks with simulated ground-truth, we created the microtubule-like structures. In specific, we utilized the *insertShape* (MATLAB function) to sketch multiple lines with random orientations on a blank canvas of 4096 × 4096 pixels (**Supplementary Fig.** 1). To simulate the effect of incomplete labeling observed in practical experiments, we applied a small Gaussian mask (σ as 2 pixels), introducing random notches along the lines (indicated by red circles in Supplementary Fig. 1). Then, to mimic the curviness of cytoskeleton, an elastic deformation was applied to bend the straight lines to curved lines in both *x* and *y* dimensions, in which the resulting image is considered its pixel size as 16.25 nm. The synthetic structures were convolved with a 150 nm size PSF and downsampled by 2-fold, for 2048 × 2048 pixels with a 32.5 nm pixel size (simulated blurred ground-truth). Finally, the images were contaminated by three different Poisson-Gaussian noise levels (10%, 20%, and 40%).

#### Pixel-wise metrics

In this work, the SSIM^38^, PSNR, and RMSE were used as metrics to evaluate the pixel-level consistency between reconstructed images and ground-truth images. For the fixed samples, we directly enlarged the exposure time to acquire the high SNR data. To remove potential small baseline background and noise, the ground-truth images were created from these high SNR images by subtracting a constant background value and subsequently filtering a small Gaussian kernel. To qualify data and model uncertainties, we adopted the standard deviation (STD) using the ten predictions from ten repetitively collected inputs or ten repetitively trained models:

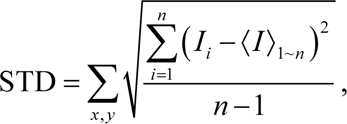

where *n* is the sequence length (default as 10); *I_i_* (*x*, *y*) represents the intensity of the *i*^th^ image in the sequence, and 〈*I* (*x*, *y*)〉_1∼*n*_ is the averaged intensity of the sequence.

#### FRC resolution

The calculation of FRC resolution requires two independent frames of identical contents under the same imaging conditions^37^. In case of confocal, SD-SIM, and STED, we repetitively acquired the same content twice. For SOFI, these two frames were generated by splitting the raw image sequence into two image subsets, e.g., the first 20 frames and the last 20 frames, and reconstructing them independently.

#### Line restoration ratio (LQR)

To quantitatively evaluate the reconstruction quality of parallel lines, we employed the line quality ratio (LQR)^12^ metric:

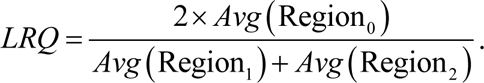

The *Region1*, *Region2*, and *Region0* represent the parallel lines and the region in between, and *Avg* calculates the mean intensity of pixels within the corresponding area. To avoid overconfident determination, we used a threshold of 0.2 to ascertain the successful separation of parallel lines.

#### Prediction uncertainty estimation

An optimized denoising algorithm can be characterized by minimal data and model uncertainties. Data uncertainty indicates the algorithm’s ability to efficiently reduce noise (minimize errors), whereas model uncertainty reflects the model’s efficiency in utilizing data (sustain performance under limited or varying data conditions)^63^. To estimate data uncertainty, we collected ten independent frames of identical contents under the same imaging conditions and fed them to the trained SN2N network. The STD of the ten resulting predictions was calculated as the data uncertainty. Regarding the model uncertainty, we repetitively trained the DNNs with ten times and inputted the same data into these ten models. The STD of the resulting predictions served as a measure of model uncertainty.

#### FWHM measurements

The FWHM values were estimated from the Gaussian fittings of the manually picked intensity profiles. Particularly, the profiles and values plotted in **Extended Data Fig. 4a** were automatically created by using the LuckyProfiler ImageJ plugin^64^, which can autonomously identify and quantify the optimal FWHM locations within images. It enabled us to select the necessary regions for FWHM calculations and apply Gaussian fitting algorithms.

### SD-SIM setup

We used two commercial SD-SIM systems to capture SR confocal images. We validated the denoising performance using a commercial fluorescent sample (the Argo-SIM slide, Argolight, France) with ground-truth patterns consisting of fluorescing double line pairs (spacing from 0 nm to 390 nm, ii_ex_= 300–550 nm, http://argolight.com/products/argo-sim) under the SpinSR10 system and conducted live-cell imaging tests using the Live-SR system.

#### SpinSR10 system

The SpinSR10 system is a commercial SD-SIM (SpinSR10, Olympus, Japan) equipped with a wide-field objective (×100/1.49 oil, APON, Olympus) and an sCMOS camera (ORCA Fusion, Hamamatsu, Japan). Four laser beams of 405 nm, 488 nm, 561 nm, and 640 nm were combined with the SD-SIM. The detection optical path adopted a further 3.2× magnification, and the total magnification was 320 times. We captured SD-SIM images in its SoRa (super-resolution) mode and collected confocal images by switching to its conventional SD-confocal mode.

#### Live-SR system

The Live-SR SD-SIM is based on an inverted fluorescence microscope (IX81, Olympus) equipped with a wide-field objective (100×/1.3 oil, Olympus), a scanning confocal system (CSU-X1, Yokogawa), and a Live-SR module (GATACA systems, France). Four laser beams of 405 nm, 488 nm, 561 nm, and 647 nm were combined with the SD-SIM. The images were captured either by an sCMOS camera (C14440-20UP, Hamamatsu) or an EMCCD camera (iXon3 897, Andor, United Kingdom).

### STED setup

We acquired STED images from two commercial STED systems.

#### Abberior STED

Long-term activities of mitochondrial cristae (PKMO^48^ labeled) were recorded by a commercial STED microscope (STEDYCON, Abberior Instruments, Germany) equipped with a wide-field objective (100×/1.45, CFI Plan Apochromat Lambda D, Nikon, Japan). PKMO was excited at 561 nm wavelength, and STED was performed using a pulsed depletion laser at 775 nm wavelength with gating of 1 to 7 ns and dwell times of 10 μs. A pixel size of 25 nm was used for STED recording and each line was scanned 1 or 10 times (line accumulations). The pinhole was set to 0.7 to 1.0 AU.

#### Leica STED

Other live-cell STED images were obtained using a gated STED (gSTED) microscope (TCS SP8 STED 3X, Leica, Germany) equipped with a wide-field objective (100×/1.40 oil, HCX PL APO, Leica). The excitation and depletion wavelengths were 488 nm and 592 nm for the Sec61β-GFP and LifeAct-GFP, 594 nm and 775 nm for the Alexa Fluor 594, 635 nm and 775 nm for the Alexa Fluor 647, 651 nm and 775 nm for SiR-tubulin. The detection wavelength range was set to 495-571 nm for GFP, 605-660 nm for Alexa Fluor 594, 657-750 nm for SiR, and 649-701 nm for Alexa Fluor 647. For comparison, confocal images were acquired in the same field before the STED imaging. All images were obtained using the LAS AF software (Leica).

### SOFI setup

#### Wide-field microscopy

The three phases of structured illumination under the same orientation can be averaged to a uniform wide-field illumination. Taking advantage of that, we use the SIM setup described in the above section to generate the wide-field images by integrating three frames (corresponding to three phases of structured illumination) on the camera plane, which enables more flexible cross-validation of SIM and SOFI-SN2N results. In other words, we employ the identical commercial inverted fluorescence microscope (IX83, Olympus) equipped with an objective (100×/1.7 HI oil, APON, Olympus) and a sCMOS (Flash 4.0 V3, Hamamatsu, Japan) camera to capture the wide-field images for our SOFI-SN2N reconstruction.

#### SD-confocal microscopy

A commercial SD-confocal microscope system (Dragonfly SD system, Andor) based on an inverted fluorescence microscope (DMi8, Leica) with a wide-field objective (100×/1.3 oil, Plan Apo, Leica), is used in this work. Four laser beams of 405 nm, 488 nm, 561 nm, and 647 nm were combined with the SD-confocal microscope. The images were captured by a sCMOS camera (Zyla 4.2 Plus, Andor).

### SIM setup

The SIM system is based on a commercial inverted fluorescence microscope (IX83, Olympus) equipped with an objective (×100/1.49 oil, UAPON, Olympus, for 2D-SIM; 100×/1.7 HI oil, APON, Olympus, for TIRF-SIM) and a multiband dichroic mirror (DM, ZT405/488/561/640-phase R; Chroma) as described previously^34^. In short, laser light with wavelengths of 488 nm (Sapphire 488LP-200) and 561 nm (Sapphire 561LP-200, Coherent) and acoustic optical tunable filters (AOTF, AA Opto-Electronic, France) were used to combine, switch, and adjust the illumination power of the lasers. A collimating lens (focal length: 10 mm, Lightpath) was used to couple the lasers to a polarization-maintaining single-mode fiber (QPMJ-3AF3S, Oz Optics). The output lasers were then collimated by an objective lens (CFI Plan Apochromat Lambda 2× NA 0.10, Nikon) and diffracted by the pure phase grating that consisted of a polarizing beam splitter, a half-wave plate, and the SLM (3DM-SXGA, ForthDD). The diffraction beams were then focused by another achromatic lens (AC508-250, Thorlabs) onto the intermediate pupil plane, where a carefully designed stop mask was placed to block the zero-order beam and other stray light and to permit passage of ±1 ordered beam pairs only. To maximally modulate the illumination pattern while eliminating the switching time between different excitation polarizations, a home-made polarization rotator was placed after the stop mask^34^. Next, the light passed through another lens (AC254-125, Thorlabs) and a tube lens (ITL200, Thorlabs) to focus on the back focal plane of the objective lens, which interfered with the image plane after passing through the objective lens. Emitted fluorescence collected by the same objective passed through a dichroic mirror, an emission filter, and another tube lens. Finally, the emitted fluorescence was split by an image splitter (W-VIEW GEMINI, Hamamatsu, Japan) before being captured by a sCMOS (Flash 4.0 V3, Hamamatsu) camera.

### Expansion microscopy setup

We employed the commercial inverted fluorescence microscope (IX83, Olympus) equipped with a wide-field objective (100×/1.49 oil, UAPON, Olympus) and a sCMOS (Flash 4.0 V3, Hamamatsu) camera to capture the wide-field images of the expanded samples.

### Imaging sample preparation

#### Cell maintenance and preparation

COS-7 cells (ATCC, CRL-1651) and HeLa cells (ATCC, CCL-2) were cultured in high-glucose DMEM (Gibco, 21063029) supplemented with 10% fetal bovine serum (FBS) (Gibco) and 1% 100 mM sodium pyruvate solution (Sigma-Aldrich, S8636) in an incubator at 37°C with 5% CO_2_ until ∼75% confluency was reached. MCF7 cells were cultured in MEM (thermo fisher, 11095072) supplemented with 10% fetal bovine serum (FBS), 0.01 mg/ml human recombinant insulin (Sigma, I9278), and 1% 100 mM sodium pyruvate solution. For the SD-SIM/SD-confocal/STED imaging experiments, 35-mm glass-bottomed dishes (Cellvis, D35-14-1-N) were used. For the wide-field and 2D-SIM imaging experiments, cells were seeded onto coverslips (H-LAF 10L glass, reflection index: 1.788, diameter: 26 mm, thickness: 0.15 mm, customized) coated with 0.01% Poly-L-lysine solution (Sigma, P4707) for 10 minutes and washed twice with Sterile Water before seeding transfected cells.

#### Live-cell samples for SD-SIM and SIM

To label late endosomes or lysosomes (Lys), we incubated COS-7 cells in 50 nM LysoTracker Deep Red (Thermo Fisher Scientific, L12492) for 45 min and washed them 3 times in PBS before imaging. To label mitochondria, COS-7 cells were incubated with 250 nM MitoTracker™ Green FM (Thermo Fisher Scientific, M7514) and 250 nM MitoTracker® Deep Red FM (Thermo Fisher Scientific, M22426) in HBSS containing Ca^2+^ and Mg^2+^ or no phenol red medium (Thermo Fisher Scientific, 14025076) at 37°C for 15 min before being washed three times before imaging. To perform nuclear staining on COS-7 cells, SPY650-DNA (Cytoskeleton, CY-SC501) was diluted to 1:1000 in PBS for ∼1 h and washed 3 times in PBS.

To label cells with genetic indicators, COS-7 cells were transfected with LifeAct-EGFP/LAMP1-EGFP/LAMP1-mChery/Tom20-mCherry/Sec61β-EGFP/Golgi-BFP/mGold^65^-Mito-N-7/DsRed-ER. The transfections were executed using Lipofectamine^TM^ 2000 (Thermo Fisher Scientific, 11668019) according to the manufacturer’s instructions. After transfection, cells were plated in pre-coated coverslips. Live cells were imaged in a complete cell culture medium containing no phenol red in a 37°C live cell imaging system. For the calcium lantern imaging in SD-SIM, the calcium signal was stimulated with a micropipette containing 10 μmol/L 5’-ATP-Na_2_ solutions (Sigma-Aldrich, A1852)^51^.

#### Samples for STED imaging

To label the ER-tubule/actin/microtubule in live cells, COS-7 cells were either transfected with Sec61β-EGFP/Lifeact-EGFP, or incubated with SiR-Tubulin (Cytoskeleton, CY-SC002) or PKMO^48^ for ∼20 mins without wash before imaging.

#### Immunofluorescence for SOFI

The COS-7 cells were grown in 35-mm glass-bottomed dishes overnight and rinsed with PBS, then immediately fixed with prewarmed 4% PFA (Santa Cruz Biotechnology, sc-281692) for 10 mins. After three washes with PBS, cells were permeabilized with 0.1% Triton® X-100 (Sigma-Aldrich, X-100) in PBS for 10 mins. Cells were blocked in 5% BSA/PBS for 1h in room temperature. Mouse anti-Tubulin DM1a (Sigma, T6199) was diluted to 1:100 and stained cells in 2.5% BSA/PBS blocking solution for 2 h at room temperature. The cells were then washed with PBS five times for 10 mins per wash and stained with biotin-XX goat anti-mouse IgG antibody (Invitrogen, B2763). The cells were then washed with PBS five times for 10 mins per wash and stained with Qdot™ 525 (QD_525_) Streptavidin Conjugate (Invitrogen, Q10143MP) for 60 mins. Finally, cells were washed five times with PBS and imaged.

#### Live-cell samples for SOFI

To label the mitochondria in live cells, COS-7 cells were transfected with Skylan-S-TOM20^66^. The transfections were executed using LipofectamineTM 2000 (Thermo Fisher Scientific, 11668019) according to the manufacturer’s instructions. After transfection, cells were plated in glass-bottomed dishes. The Skylan-S was under sequential illumination with a 405 nm laser (low power) when imaging. In addition, live cells were imaged in complete cell culture medium containing no phenol red in a 37°C live cell imaging system.

### Samples preparation for expansion microscopy

#### Sample expansion

The sample expansion was performed as previously described^29, 67^. The labeled cells were incubated with 0.1 mg mL^−1^ of Acryloyl-X (AcX, Thermo, A20770) diluted in PBS overnight at r.t. and washed three times with PBS. To prepare the gelation solution, freshly prepared 10% (w/w) N,N,N′,N′-tetramethylethylenediamine (TEMED, Sigma, T7024) and 10% (w/w) ammonium persulfate (APS, Sigma, A3678) were added to the monomer solution (1×PBS, 2 M NaCl, 2.5% (w/v) acrylamide (Sigma, A9099), 0.15% (w/v) N,N′-methylenebisacrylamide (Sigma, M7279) and 8.625% (w/v) sodium acrylate (Sigma, 408220)) to a final concentration of 0.2% (w/w) each. Next, the cells were embedded with the gelation solution first for 5 min at 4 °C, and then for 1 h at 37 °C in a humidified incubator. The gels were immersed into the digestion buffer (50 mM Tris, 1 mM EDTA, 0.1% (v/v) Triton X-100, and 0.8 M guanidine HCl, pH 8.0) containing 8 units mL^−1^ proteinase K (NEB, P8107S) at 37 °C for 4 h, and then placed into ddH_2_O to expand. Water was changed 4-5 times until the expansion process reached a plateau. By determining the gel sizes of before and after the expansion, we quantified the expansion factor to be 4.5 times. The gels were immobilized on poly-D-lysine-coated glass No. 1.5 cover-glass for further imaging.

#### α-tubulin immunostaining

COS-7 cells were seeded in a Lab-Tek II chamber slide (Nunc, 154534). Cells were firstly extracted in the cytoskeleton extraction buffer (0.2% (v/v) Triton X-100, 0.1 M PIPES, 1 mM EGTA, and 1 mM MgCl_2_, pH 7.0) for 1 min at room temperature (r.t.). Next, the extracted cells were fixed with 3% (w/v) formaldehyde and 0.1% (v/v) glutaraldehyde for 15 min, reduced with 0.1% (w/v) NaBH_4_ in PBS for 7 min, and washed three times with 100 mM glycine. Then the cells were permeabilized with 0.1% (v/v) Triton X-100 for 15 min, and blocked with 5% (w/v) BSA in 0.1% (v/v) Tween 20 for 30 min. for antibody staining, the cells were incubated with monoclonal rabbit anti-α-tubulin antibody (EP1332Y, 1:250 dilution, Abcam, ab52866) in antibody dilution buffer (2.5% (w/v) BSA in 0.1% (v/v) Tween 20) overnight at 4 °C, washed three times with 0.1% (v/v) Tween 20, incubated with Alexa Fluor 488-conjugated F(ab’)2-goat anti-rabbit secondary antibody (1:1,000 dilution, Thermo, A11070) in antibody dilution buffer for 2 h at r.t. and washed three times with 0.1% (v/v) Tween 20.

#### Sec61β-GFP transfection

COS-7 cells were seeded in a Lab-Tek II chamber slide (Nunc, 154534) and cultured to reach around 50% confluence. For transient transfection of Sec61β-GFP in a single well, 500 ng plasmid and 1 μL of X-tremeGENE HP (Roche) were diluted in 20 μL Opti-MEM sequentially. The mixture was vortexed, incubated for 15 min at r.t., and applied to cells. 24 h after transfection, the cells were washed three times with PBS and fixed as described in the α-tubulin immunostaining experiment.

### Open-sourced datasets

#### STED images with different excitation/depletion laser powers

We used the published dataset from ref.^47^ for testing our SN2N’s denoising performance on STED images. A custom STED system incorporating a SPAD array detector with 5 × 5 elements was used for data collection. A series of images (**Fig. 5a**) were collected from living HeLa cells labeled with SIR tubulin under gradually increased STED power (0%, 5%, 8%, 12%, 16%, 19%, 30%, 50%, and 80% depletion laser powers). Another series of images (**Supplementary Fig. 7**) were acquired from α-tubulin labeled fixed HeLa with gradually increased STED power (0%, 10%, 20%, 25%, 30%, 40%, and 90% depletion laser powers). With 0% depletion laser power, the system was switched to a conventional point-scanning confocal mode. We used the direct sum and the reassigned sum of signals from 25 elements as confocal/STED and ISM/STED-ISM images, respectively.

#### BioSR SIM dataset

We used the open-sourced dataset, the BioSR dataset from ref.^50^, for evaluating the denoising performance on SIM images. The CCPs (clathrin-coated pits), microtubules, and ER data under different noise levels were used as SIM images of high, medium, and low SNR conditions in this work.

### Image rendering and processing

We used the ‘biop-12colors’ color map to color code the 3D volumes in **Fig. 4b**, **4e**, **4f**, **Extended Data Fig. 9e, 9f, and Extended Data Fig. 10a-10f**. The 3D volumes in **Fig. 4g**-**4i**, and **Extended Data Fig. 5a, 5c, 5e** were rendered using the Microscape software (https://www.microscape.xyz). All data processing was achieved using Python scripts, MATLAB, and ImageJ. All figures were prepared with MATLAB, ImageJ, Microsoft Visio, and OriginPro, and videos were all produced with and Microsoft PowerPoint and our light-weight MATLAB framework, which is available at https://github.com/WeisongZhao/img2vid.

## Data availability

All the data that support the findings of this study are available from the corresponding author upon request.

## Code availability

The tutorials and the updating version of our SN2N can be found at https://github.com/WeisongZhao/SN2N.

## Acknowledgments

We thank the assistance of Tianyan Liu from Zhixing Chen’s lab at the Peking university for STED imaging of PKMO-labeled mitochondrial cristae. This work was supported by the National Key Research and Development Program of China (grant no. 2022YFC3400600 to L. C.), the National Natural Science Foundation of China (grant no. 62305083 to W. Z., grant no. T2222009 to H. L., grant no. 32227802 to L. C., grant no. 81925022 to L. C., grant no. 92054301 to L. C., grant no. 32301257 to S. Z., grant no. 32071458 to H. M.), the Young Elite Scientists Sponsorship Program by China Association for Science and Technology (grant no. 2023QNRC001 to W. Z.), and the Heilongjiang Provincial Postdoctoral Science Foundation (grant no. LBH-Z22027 to W. Z.), the Natural Science Foundation of Heilongjiang Province (grant no. YQ2021F013 to H. L.), the Beijing Natural Science Foundation (grant no. Z20J00059 to L. C.), Guangdong Basic and Applied Basic Research Foundation (grant no. 2022A1515011683 to J. H.). L. C. acknowledges support by the High-performance Computing Platform of Peking University.

## Author contributions

W. Z. conceived the research and supervised the project; L. Q. implemented the corresponding software; S. Z., X. Y., and K. W. performed the experiments and collected the data; Q. L. analyzed the data and prepared the figures; L. Q., and Y. H. prepared the videos; X. L., H. M., G. H., W. C., C. G., J. H., J. T., H. L., and L. C. participated in discussions during the development of the manuscript; W. Z. and L. Q. wrote the manuscript with input from all authors; All authors participated in the discussions and data interpretation.

## EXTENDED DATA FIGURES

**Extended Data Fig. 1.**
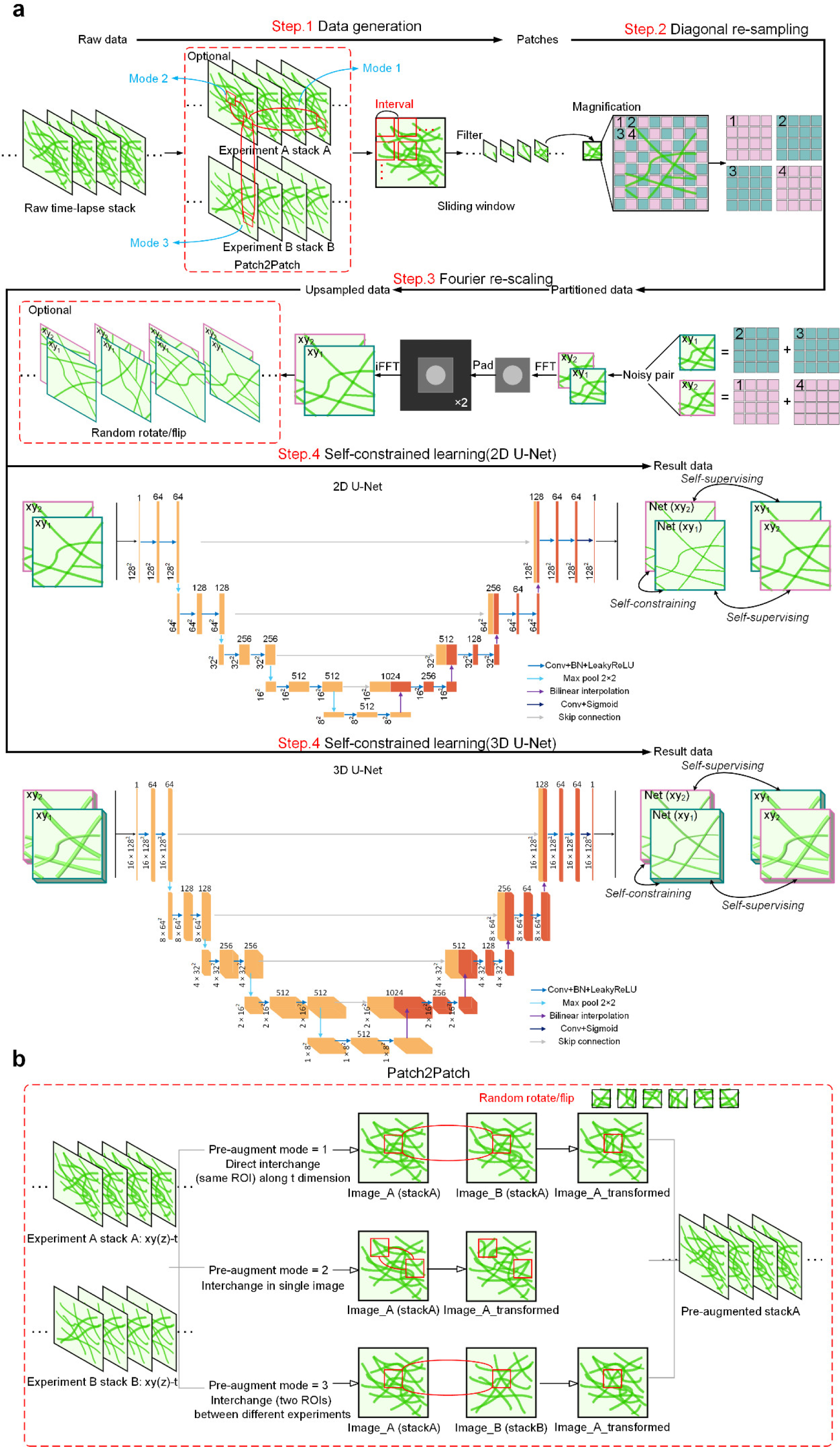
Workflow of SN2N and network architectures. **a**, Detailed flow diagram of SN2N (**Methods**). Steps 1-3, self-supervised data generation. First, the data pre-augmentation (optional) is performed using the Patch2Patch (random patch transformations in multiple dimensions) strategy. After that, a sliding window approach is employed to generate small patches suitable for input into the network for training. Subsequently, the spatial diagonal resampling strategy followed by Fourier upsampling is used to create paired SN2N data. Additionally, basic augmentations such as rotation and flipping (optional) are applied to the generated data pairs. Step 4: self-constrained learning process. SN2N utilizes the classical U-Net network and selects either the 2D U-Net or 3D U-Net based on the input data dimensions. The generated paired images are considered as one training example, and the resulting two predictions are used to calculate the loss for back propagation. **b**, Patch2Patch (P2P) pipeline (**Methods**). It includes three available modes for augmentation along the temporal axis, in a single image, and between different experiments.

**Extended Data Fig. 2.**
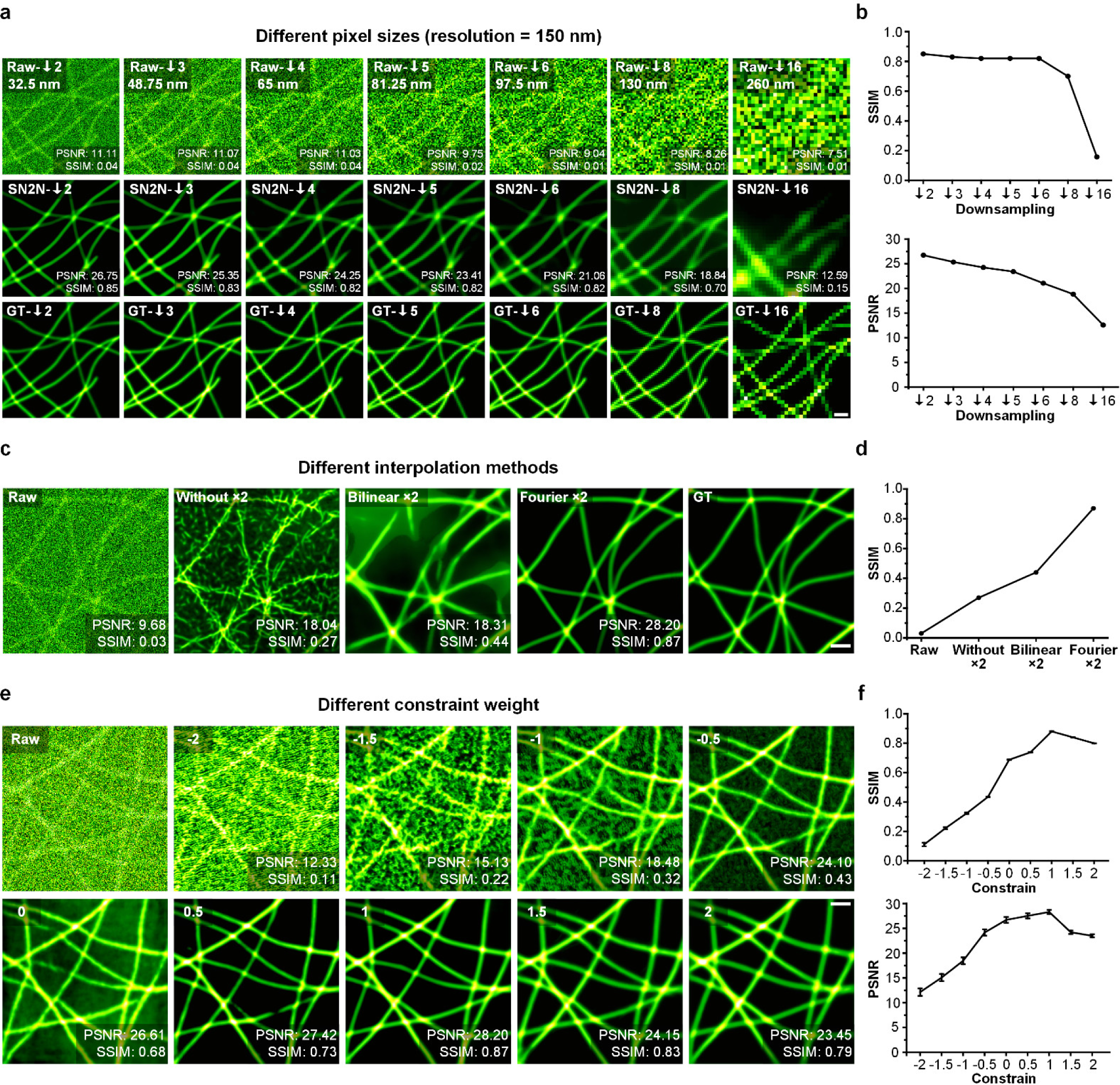
Testing results of different pixel sizes, interpolation methods, and constraint weights. **a**, SN2N denoising results under different pixel sizes with same resolution. From top to bottom: Raw images, SN2N results, and clean ground truth images. The synthetic structures (16.25 nm pixel size) were convolved with a 150 nm size PSF and downsampled by 2, 3, 4, 5, 6, 8, and 16 times (from left to right). Then the resulting images with different pixel sizes (labeled on the top left corner) were injected with Poisson-Gaussian noise on the same level (40%, Level 1). SSIM and PSNR values of SN2N results are marked on the bottom right corner. **b**, Average SSIM (top) and PSNR (bottom) values from data under different downsampling rates. **c**, SN2N denoising results under different interpolation methods. From left to right: Raw input, SN2N results using data without interpolation, with bilinear interpolation, and our Fourier interpolation as training sets, and ground truth image. **d**, Average SSIM value of SN2N under different interpolation strategies. **e**, SN2N results under different self-constrained regularization weights (values labeled on the top left corner). SSIM and PSNR values of SN2N results are marked in the bottom right corner. **f**, Average SSIM and PSNR values (*n* = 10). Error bars: s.e.m. Experiments were repeated ten times independently with similar results; scale bars, 1 µm.

**Extended Data Fig. 3.**
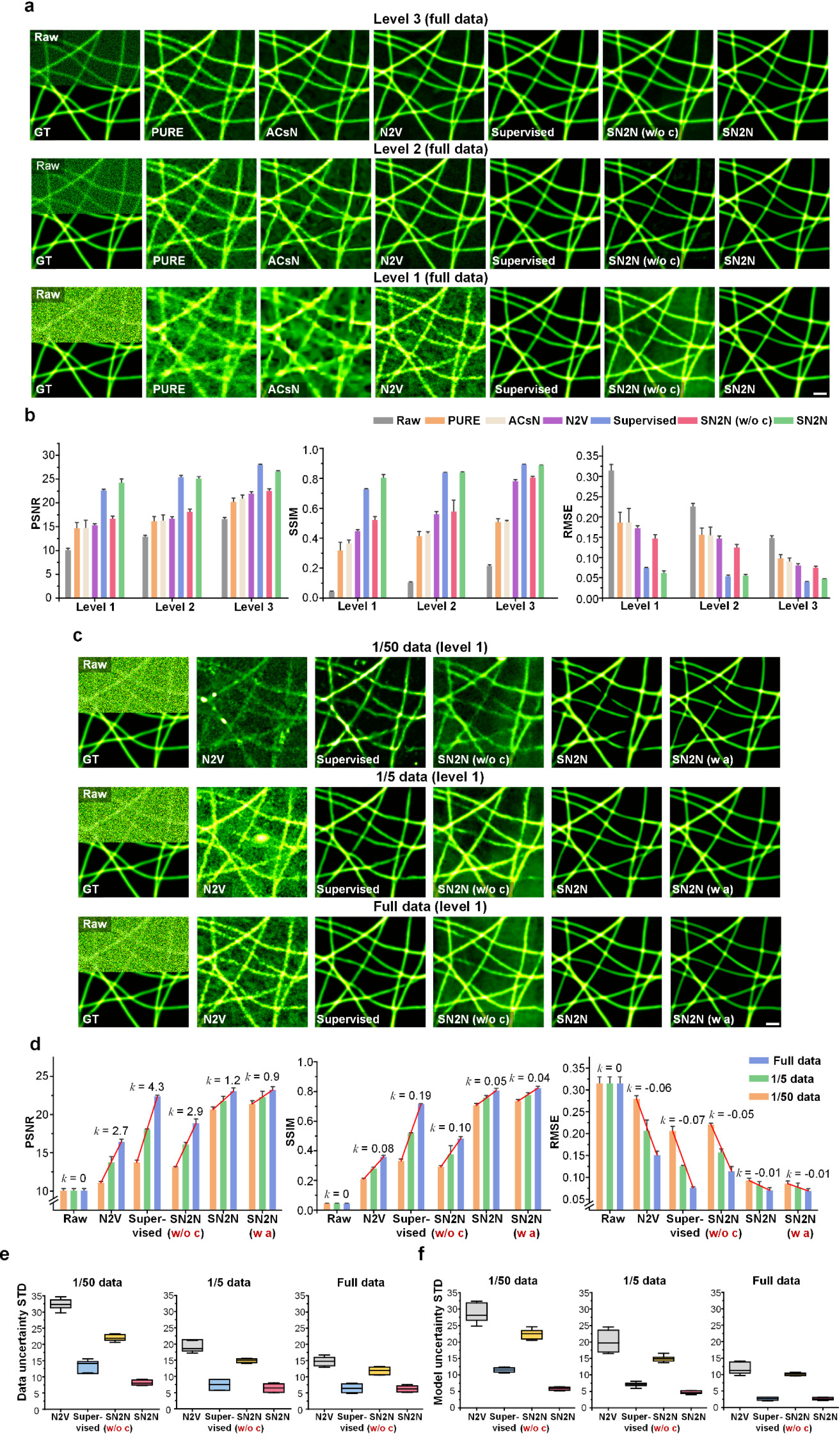
Testing results of different noise levels and data amounts. **a**, Denoising results of various methods under three different noise levels (Level 3, Level 2, and Level 1, from top to bottom) using the full training set. From left to right: Raw input, denoising results of PURE, ACsN, N2V, supervised learning (’Supervised’), SN2N without constraint (’SN2N w/o c’), and full SN2N. **b**, Quantitative comparisons of the results shown in **a** using PSNR (left), SSIM (middle), and RMSE (left) metrics (*n* = 10). **c**, Denoising results of learning-based methods using three different amounts (1/50, 1/5, and full data, from top to bottom) of training data under Level 1 of noise. **d**, Quantitative comparison of the results shown in **c** using PSNR (left), SSIM (middle), and RMSE (left) metrics (*n* = 10). *k* denotes the slope (red lines) of the corresponding metric values along the data increment. **e**-**f**, Data uncertainty (**e**) and model uncertainty (**f**) of neural network models trained by different data amounts. Average standard derivation (STD) values calculated from ten predictions of ten repetitively acquired data or ten repetitively trained models. Experiments were repeated three times independently with similar results; scale bars, 1 µm.

**Extended Data Fig. 4.**
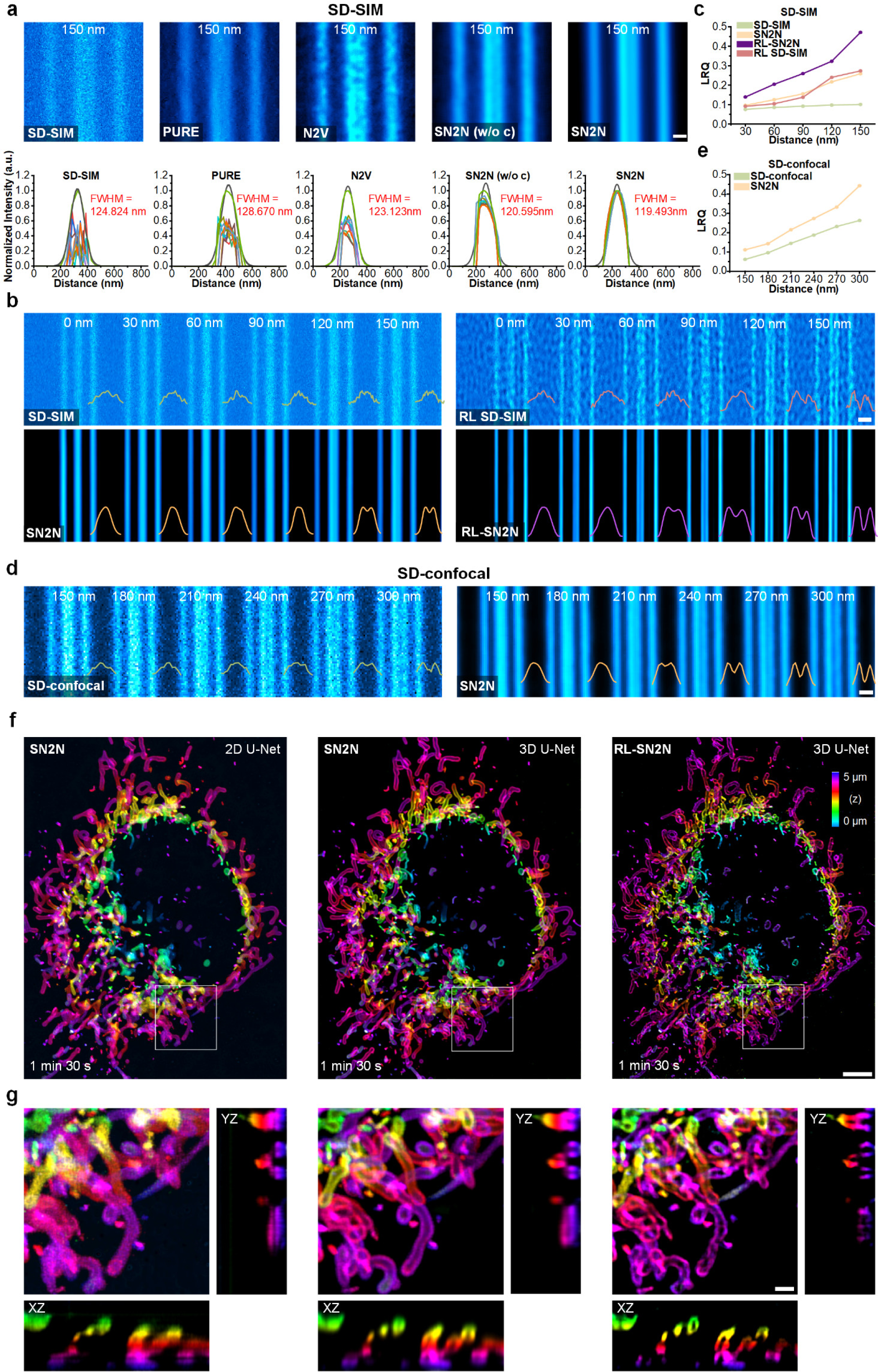
Comparisons of SN2N versus RL-SN2N using SD-SIM and applying SN2N on SD-confocal microscopy. **a**, Zoomed-in views (top) and FWHM distribution plots (bottom, calculated by LuckyProfiler) of denoising results by different methods (*c.f.*, Fig. 2a). **b**, Comparison of SN2N and RL-SN2N. Top: SD-SIM (left) and its RL result (right); Bottom SN2N result (left) and RL-SN2N result (right). **c**, LRQ values of results in **b**. **d**, SN2N denoising results (right) of SD confocal image (left) recording the Argo-SIM slide. **e**, LRQ values of results in **d**. **f**, Comparisons of SN2N with 2D U-Net (left), SN2N with 3D U-Net (middle), and RL-SN2N with 3D U-Net (right) of volumetric data (*c.f.*, Fig. 3b). **g**, Magnified views and their *xz* and *yz* cross-sections from white boxed regions in **f**. Scale bars, 500 nm (**a**, **b**, **d**); 1 µm (**f**, **g**);

**Extended Data Fig. 5.**
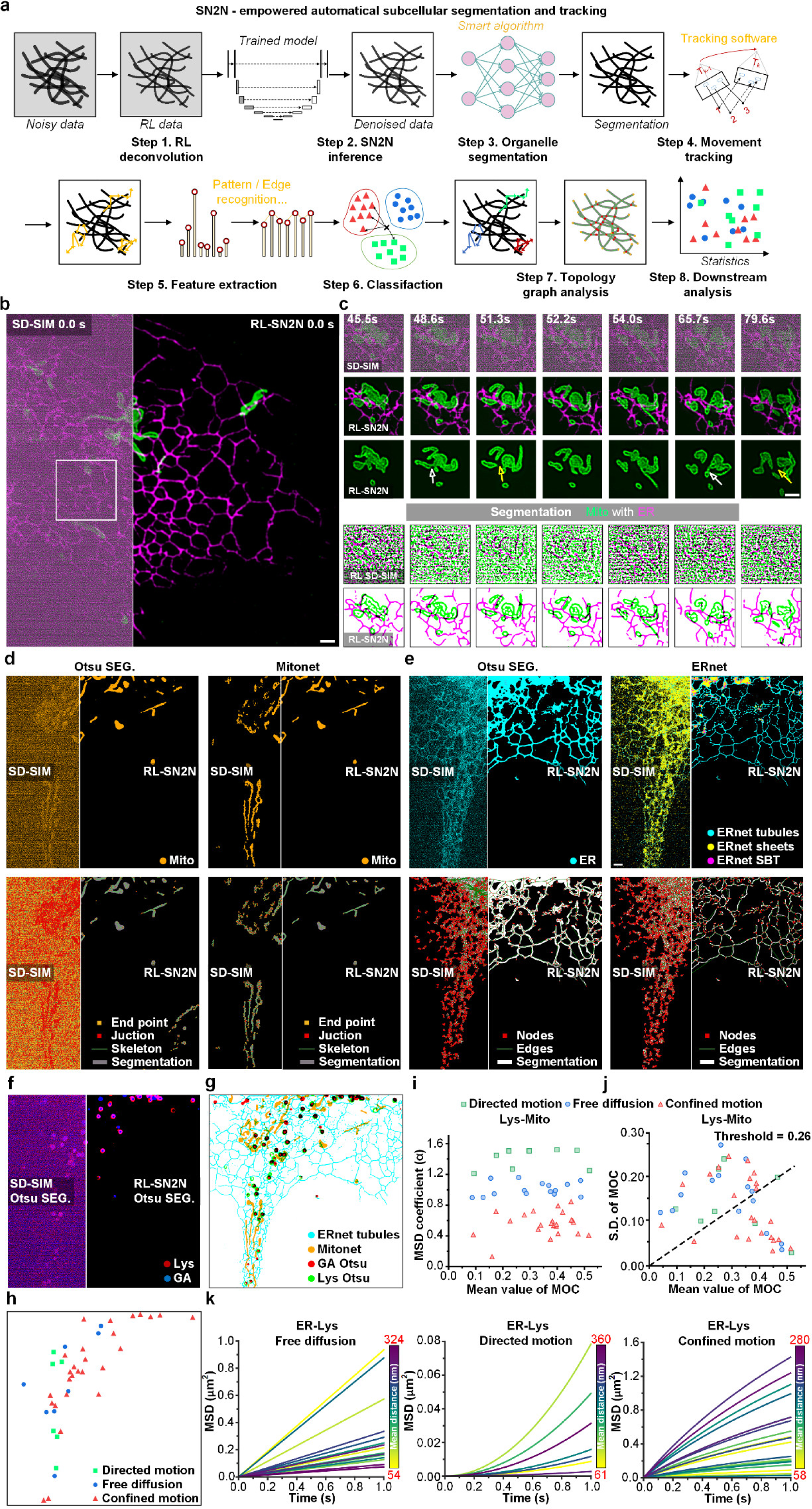
SN2N-empowered automated subcellular segmentation and tracking. **a**, Workflow. Step 1, RL deconvolution; Step 2, RL-SN2N inference; Step 3, segmentation; Step 4, tracking. Step 5, extraction of motion features. Step 6, classification. Step 7, topology graph construction. Step 8, specific downstream analysis. **b**, A representative example for dual-color SR imaging of mitochondria (Mito, green) and ER (magenta) labeled with Tom20-mCherry and Sec61β-EGFP in live COS-7 cells under raw SD-SIM (left) and RL-SN2N (right). **c**, The white box in **c** is enlarged and shown at seven time points under different configurations. From top to bottom: Raw SD-SIM, dual-color RL-SN2N, single-channel (Mito) RL-SN2N, RL SD-SIM segmentation, and RL-SN2N segmentation results. The yellow and white arrows indicate the mitochondrial fission and before fission, respectively. **d**, **e**, Results of Mito (**d**) and ER (**e**) segmentations (first row) using the Otsu hard threshold (first column) and Mitonet/ERnet (second column) and their skeletonizations (second row) under SD-SIM (left) and RL-SN2N (right). **f**, Otsu segmentation results for Lys (red) and GA (blue) under SDSIM (left) and RL-SN2N (right). **g**, A representative 4-color segmentation result under RL-SN2N. **h**, Spatial distribution of Lys assigned with different motion behaviors. **i**, Distribution of estimated *α* values of Lys versus their temporal average of minimum distances to Mito (*n* = 46). **j**, Distribution of the Lys-Mito MOCs’ standard deviation (S.D.) versus their mean values. **k**, Illustrations of the MSD curves for different motion behaviors of Lys. Curves are color-coded by the corresponding ER-Lys distances. Scale bars, 2 μm (**b**, **c**, **e**).

**Extended Data Fig. 6.**
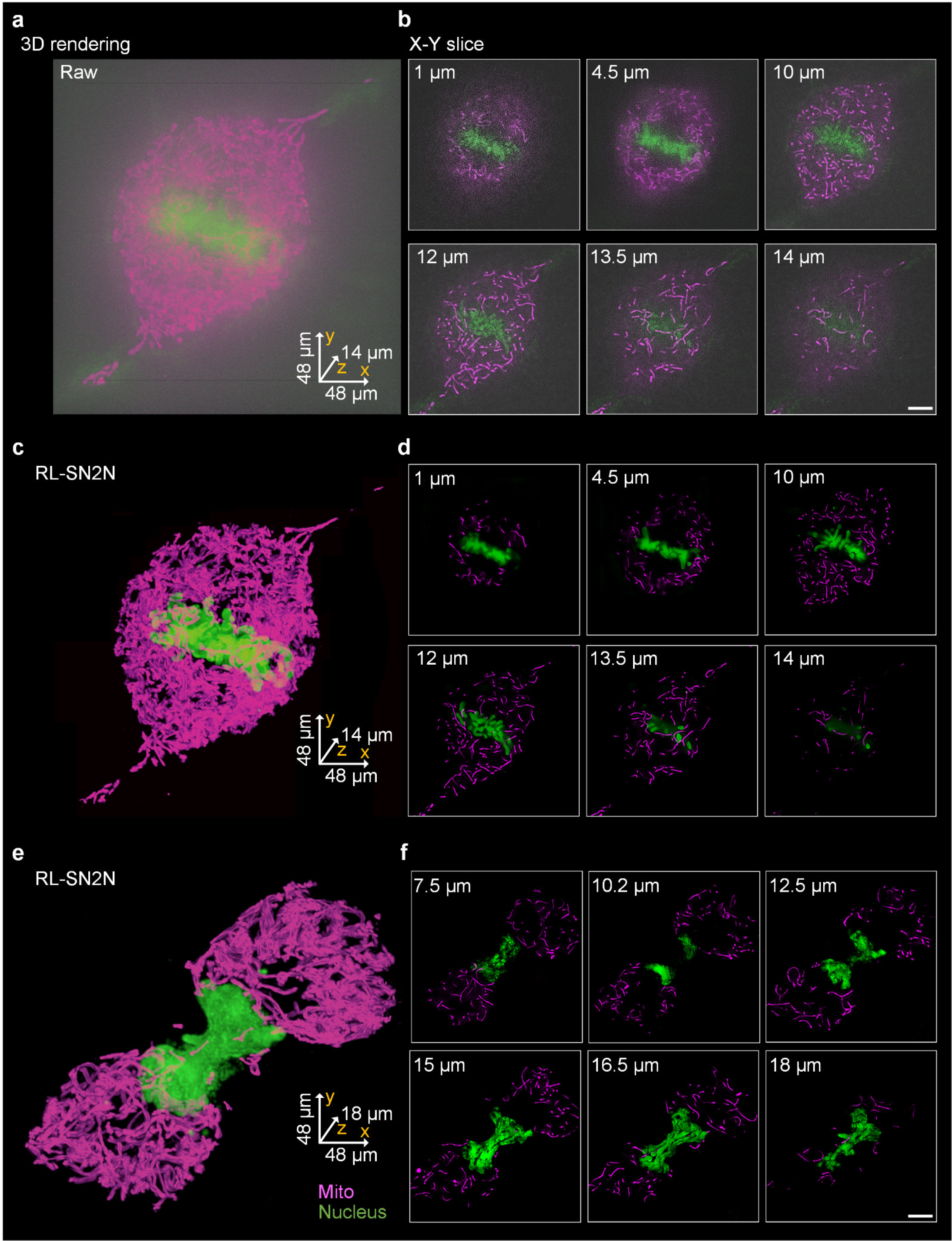
RL-SN2N can suppress noise in undersampled data from EMCCD SD-SIM. **a**, 3D renderings of live COS-7 cells labeled with Hoechst (green) and MitoTracker Deep Red (magenta) under raw SD-SIM equipped with an EMCCD camera (94 nm pixel size versus <150 nm resolution). **b**, Representative lateral slices from volume in **a**. **c**, RL-SN2N results of **a** with additional 2× upsampling before RL deconvolution (47 nm pixel size). **d**, Representative lateral slices from volume in **c**. **e**, Another RL-SN2N time point. **f**, Representative lateral slices from volume in **e**. Scale bars, 5 μm.

**Extended Data Fig. 7.**
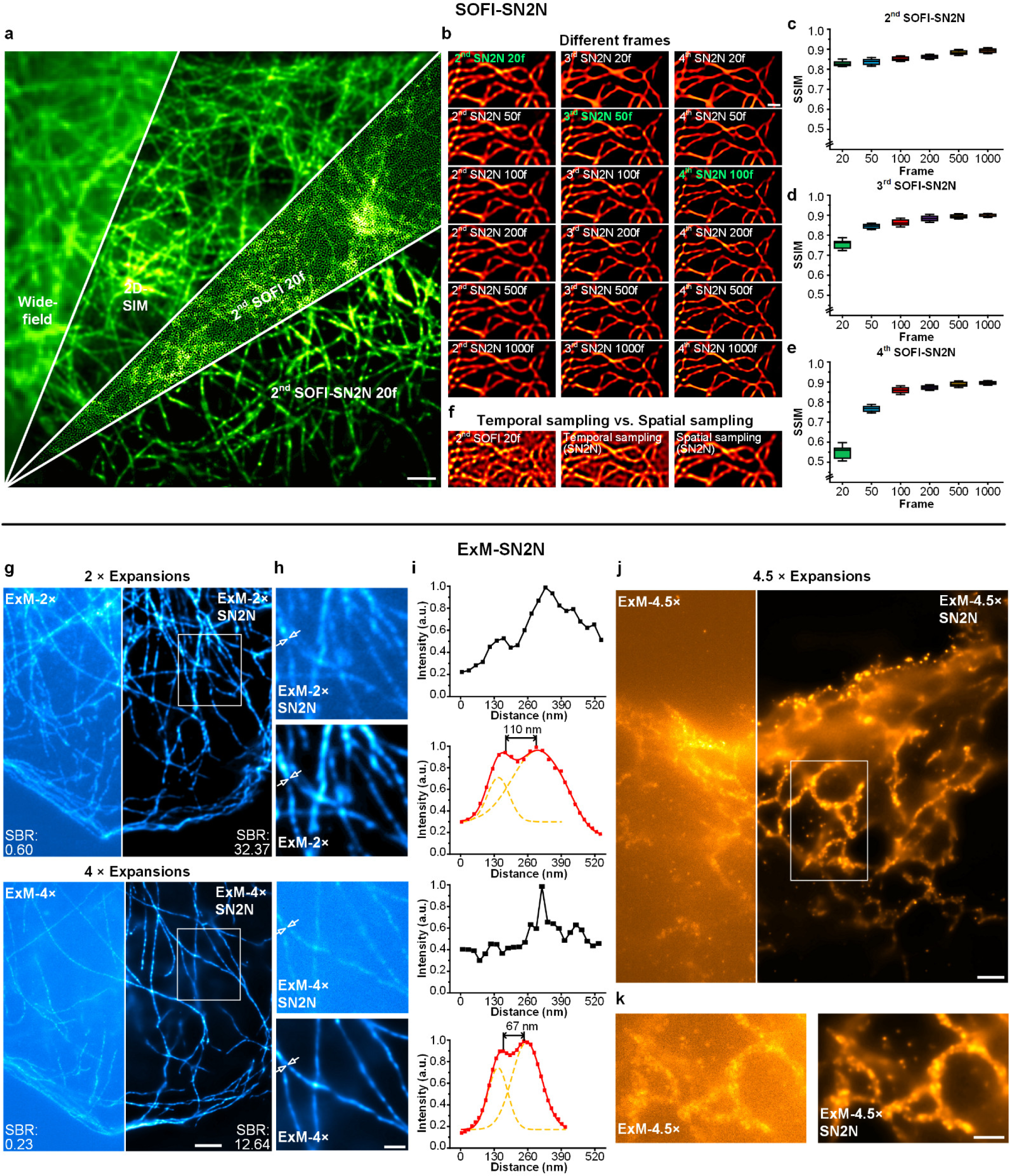
Full data of SOFI-SN2N results and SN2N-assisted expansion microscopy (ExM-SN2N). **a**, The entire view of the wide-field, 2D-SIM, 2^nd^ order SOFI with 20 frames (2^nd^ SOFI 20f), 2^nd^ order SOFI-SN2N with 20 frames images (clockwise arranged) (*c.f.*, Fig. 6b). **b**, SN2N results of 2^nd^, 3^rd^, and 4^th^ orders SOFI (from left to right) using 20, 50, 100, 200, 500, 1000 frames (from top to bottom) (*c.f.*, Fig. 6e). **c**-**e**, Average SSIM values of 2^nd^ (**c**), 3^rd^ (**d**), and 4^th^ (**e**) SOFI-SN2N results (*n* = 5). (**f**) Comparison of temporal and spatial sampling methods. From left to right: SOFI reconstruction, SN2N result using temporal sampling (the first 20 frames vs. the second 20 frames), and SN2N result using spatial sampling. **g**, A 2 times-expanded (2×, top) and 4-times expanded (4×, bottom) COS-7 cell was immunostained with a primary antibody against α-tubulin and a second antibody conjugated with Alexa Fluor 488 under wide-field microscopy (left) and its SN2N denoised result (right). Signal-to-background ratios (SBR) are labeled. **h**, Magnified views of the white boxed regions in **a** under ExM (top) and SN2N denoised results (bottom). **i**, Intensity profiles and multiple Gaussian fitting of the filaments indicated by the white arrows in **h**. Numbers represent the distances between peaks; a.u., arbitrary units. **j**, A 4.5-times expanded (4.5×) COS-7 cell labeled with Sec61β–GFP under wide-field microscopy (left) and its SN2N denoised result (right). **k**, Enlarged regions enclosed by the white box in **j** seen under ExM-4.5× (left) and its SN2N result (right). Centerline, medians; limits, 75% and 25%; whiskers, maximum and minimum; error bars, s.e.m., scale bars, 2 µm (**a**), 1 µm (**b**, **h**, **j**, **k**), and 5 μm (**g**).

**Extended Data Fig. 8.**
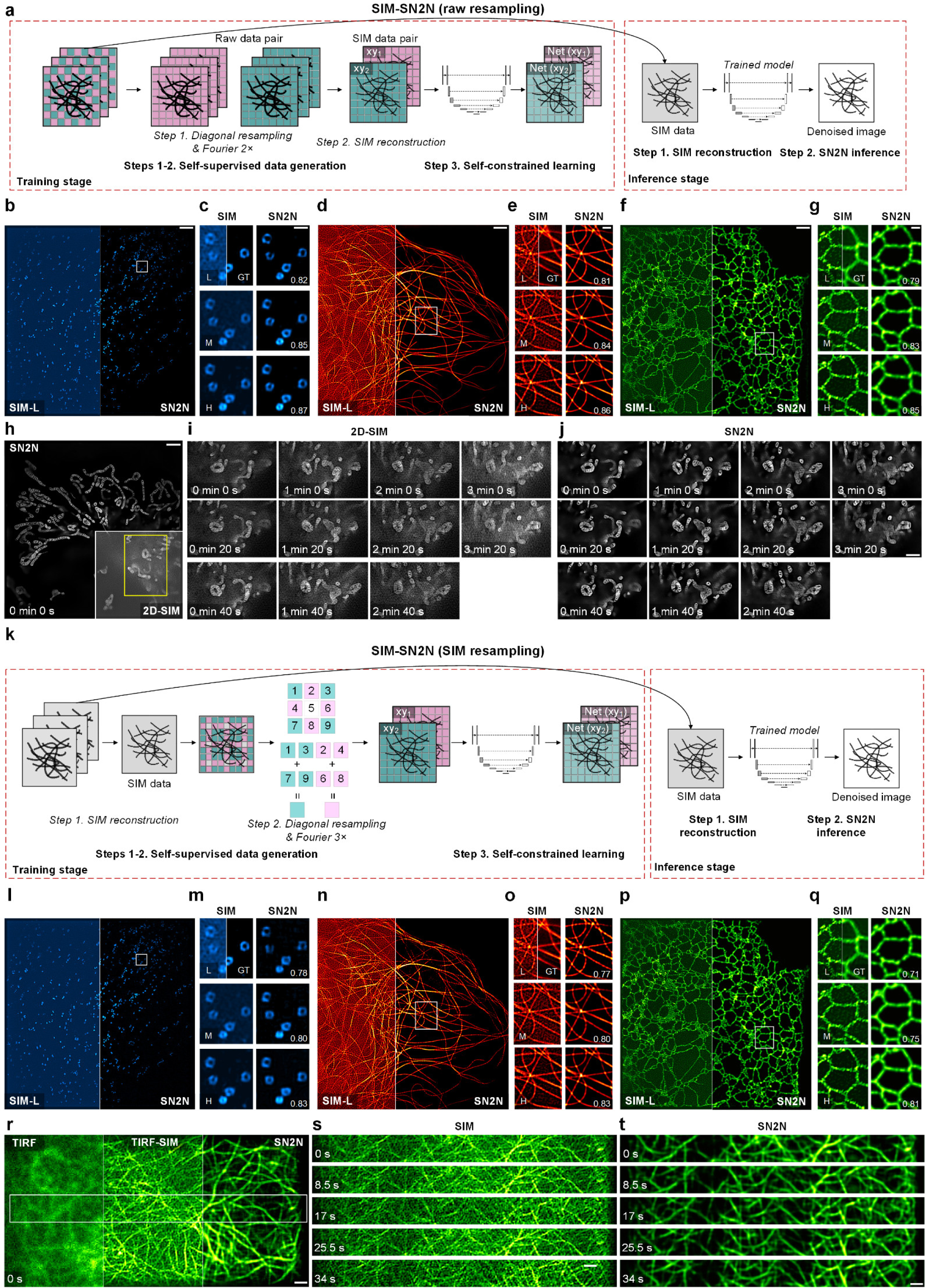
SN2N removes random, non-continuous artifacts in low-SNR SIM with two strategies. **a**, Pipeline of SIM-SN2N using raw image resampling strategy. The self-supervised data generation is applied on the 9 raw images followed by the SIM reconstruction. The resulting paired SR SIM images are the training set. At the inference stage, the SR SIM image is directly fed into the trained SIM-SN2N model. **b**, **d**, **f**, clathrin-coated pits (CCPs, **b**), microtubules (**d**), and ER (**f**), recorded by SIM under low-SNR condition (SIM-L, left) and their SN2N results (right). **c**, **e**, **g**, SIM reconstructions (left) of CCPs (**c**), microtubules (**e**), and ER (**g**) under low (L, top), medium (M, middle), and high (H, bottom) SNR conditions and their SN2N results (right). The ground truth (GT) images are displayed on the right part of the corresponding low-SNR SIM images. SSIM values are labeled on the bottom right corners. **h**, The mitochondrial cristae structures in live COS-7 cells labeled with MitoTracker Green under 2D-SIM (bottom left boxed region) and SN2N-SIM imaging at the first time point. **i**, **j**, Representative montages of 11 time points from yellow boxed region in **h** under 2D-SIM (**i**) and SIM-SN2N (**j**). **k**, Workflow of SIM-SN2N using SIM image resampling strategy. After SIM reconstruction, we apply a self-supervised data generation method specially designed for SIM, in which the resampling operation acts in 3 × 3 pixels (1 + 3 + 7 + 9 versus 2 + 4 + 6 + 8) followed by a 3× Fourier interpolation. **l**, **n**, **p**, CCPs (**l**), microtubules (**n**), and ER (**p**), recorded by SIM under low-SNR condition (left) and their SN2N results (right). **m**, **o**, **q**, SIM reconstructions (left) of CCPs (**m**), microtubules (**o**), and ER (**q**) under low (top), medium (middle), and high (bottom) SNR conditions and their SN2N results (right). The ground truth images are displayed on the right part of the corresponding low-SNR SIM images. SSIM values are labeled on the bottom right corners. **r**, A representative living COS-7 cell labeled with LifeAct–EGFP under ultrafast TIRF (left), TIRF-SIM (middle), and SIM-SN2N (right). **s**, **t**, Enlarged regions enclosed by the white box in **r**, under TIRF-SIM (**s**) and SIM-SN2N (**t**). Experiments were repeated three times independently with similar results. Scale bars, 2 μm (**b**, **d**, **f**, **h**) and 1 μm (**c**, **e**, **g**, **j**, **r**, **t**)

**Extended Data Fig. 9.**
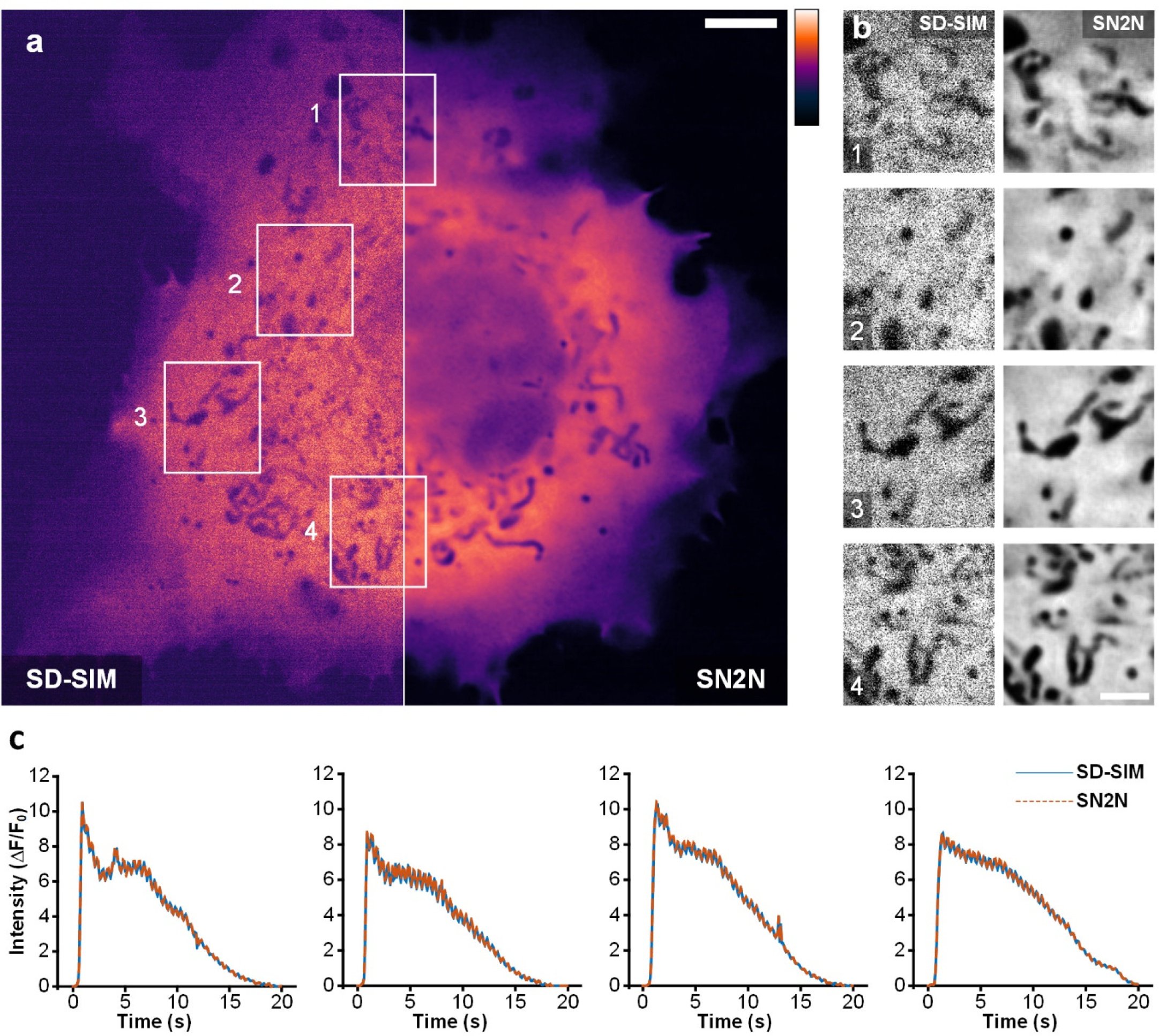
SN2N maintains the linear response of Ca^2+^ transients obtained by the SD-SIM. **a**, A representative live COS-7 cell was transfected with GCaMP6s, stimulated with ATP (10 μM). One snapshot under the SD-SIM (left) and after the SN2N (right) were shown. **b**, Magnified views of regions enclosed by white boxes 1-4 in **a**. **c**, ATP stimulated calcium traces from corresponding macrodomains in **b**. Experiments were repeated three times independently with similar results. Scale bar, 5 µm (**a**), 2 µm (**b**).

**Extended Data Fig. 10.**
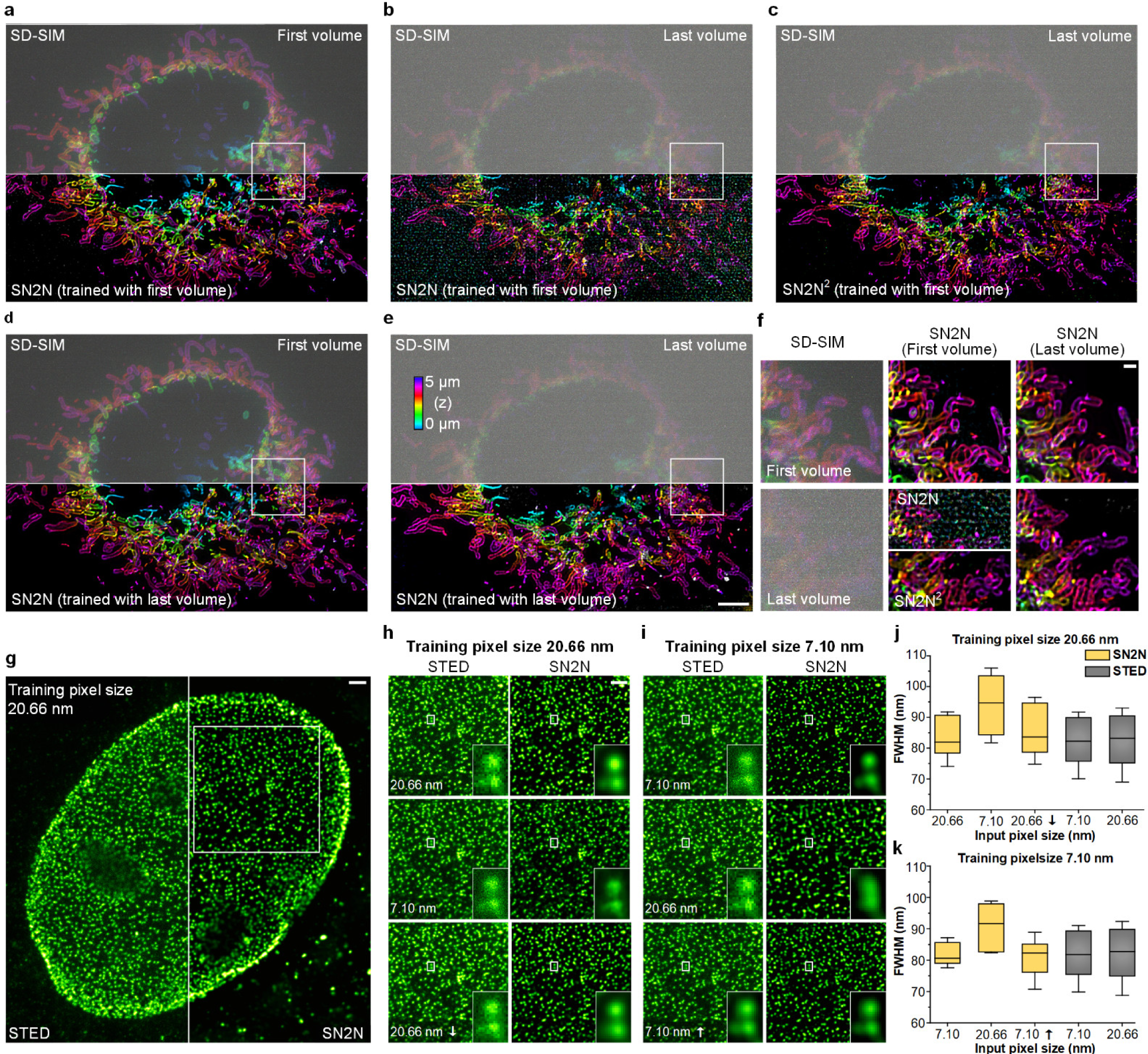
Generalization of unsupervised learning methods across different SNR conditions and pixel sizes (structural scales). **a**-**f**, Testing generalization of SN2N across different SNR conditions. **a**-**c**, Color-coded 3D distributions of all mitochondria (labeled with Tom20–mCherry) of a live COS-7 cell (*c.f.*, Fig. 3b) at the first volume (0 min) (**a**) and the last volume (2.5 min) (**b**) under SD-SIM (top) and SN2N trained with the first volume (bottom), and the SN2N prediction of SN2N perdition (SN2N^2^, bottom) from the last SD-SIM volume (top). **d**, **e**, SN2N predictions (bottom, trained with the last SD-SIM volume) from the first (0 min) (**d**) and the last (2.5 min) (**e**) SD-SIM volumes (top). **f**, Zoom-in views from white-boxed regions in **a**-**e**. First column: 0 min (top) and 2.5 min (bottom) SD-SIM; second column: SN2N and SN2N^2^ (bottom half of bottom) results (trained with 0 min SD-SIM volume) of 0 min (top) and 2.5 min (bottom) SD-SIM; SN2N results (trained with 2.5 min SD-SIM volume) of 0 min (top) and 2.5 min (bottom) SD-SIM. **g**-**k**, Testing generalization of SN2N across different pixel sizes. **g**, Nuclear pores in HeLa cells were labeled with an anti-Mab414 primary antibody and the Alexa594 secondary antibody and observed under STED and STED-SN2N configurations. **h**, STED images (left) of 20.66 nm pixel size (top), 7.10 nm pixel size (middle), and 20.66 nm pixel size subsampled from 7.10 nm (bottom), and their SN2N results (right) from model trained by data of 20.66 nm pixel size. **i**, STED images (left) of 7.10 nm pixel size (top), 20.66 nm pixel size (middle), and 7.10 nm pixel size Fourier upsampled from 20.66 nm (bottom), and their SN2N results (right) from the model trained by data of 7.10 nm pixel size. **j**, **k**, Average FWHM values of STED (gray) and SN2N results (yellow) from the model trained by data of 20.66 nm pixel size (**j**) and 7.10 nm pixel size (**k**) (*n* = 5). Centerline, medians; limits, 75% and 25%; whiskers, maximum and minimum; error bars, s.e.m. Experiments were repeated three times independently with similar results. Scale bar, 5 µm (**e**), 1 µm (**f**-**h**).

