## Supplementary Information for "Self-inspired learning to denoise for live-cell super-resolution microscopy"

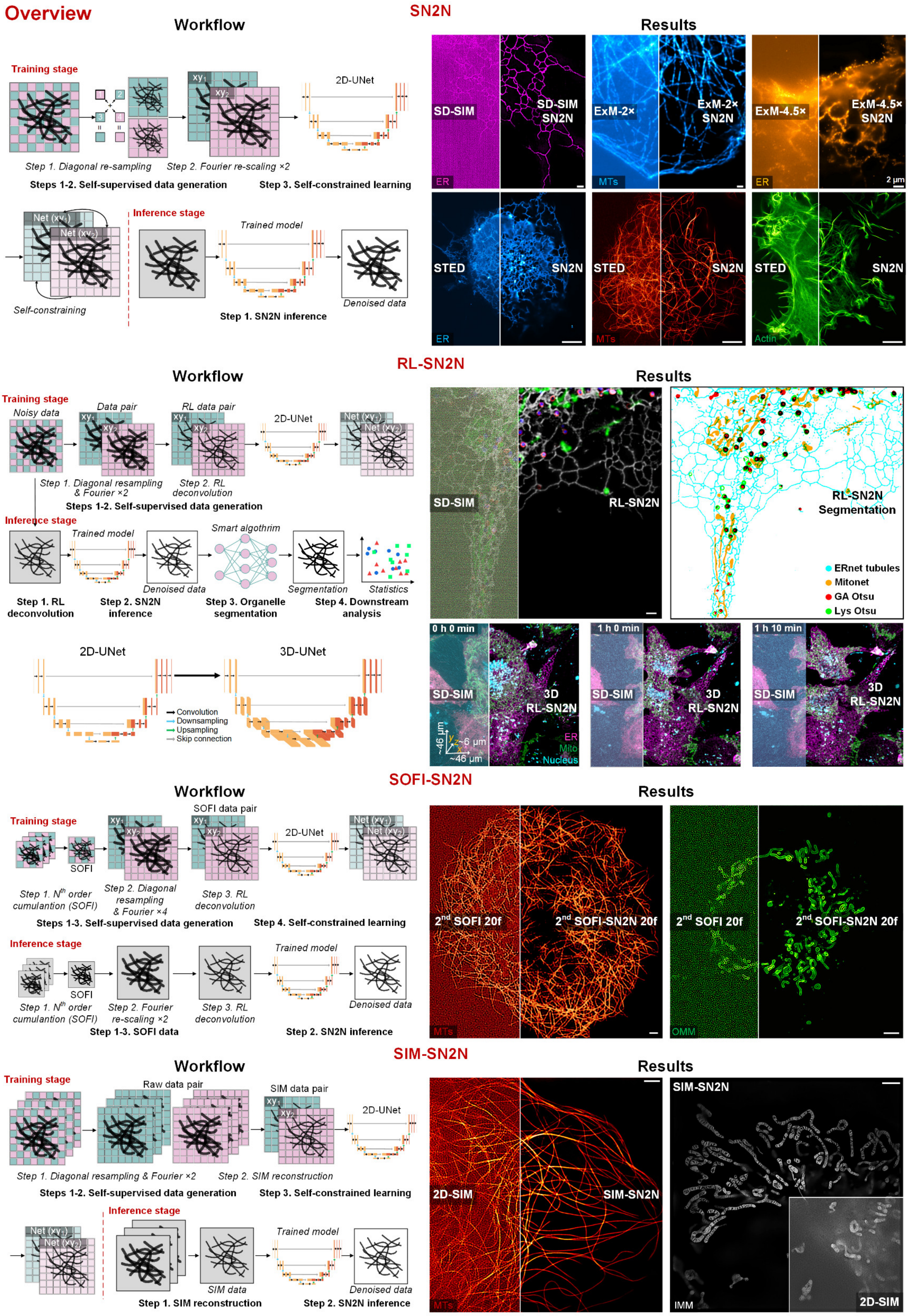
Overview
SN2N
Workflow
Results
Training stage
Step 1. Diagonal re-sampling
Step 2. Fourier re-scaling ×2
2D-UNet
Steps 1-2. Self-supervised data generation
Step 3. Self-constrained learning
Inference stage
Trained model
Self-constraining
Denoised data
Step 1. SN2N inference
SD-SIM
SD-SIM SN2N
ExM-2×
ExM-2× SN2N
ExM-4.5×
ExM-4.5× SN2N
ER
MTs
2 μm
STED
SN2N
Actin
RL-SN2N
Noisy data
Data pair
RL data pair
Step 1. Diagonal resampling & Fourier ×2
Step 2. RL deconvolution
Steps 1-2. Self-supervised data generation
Smart algothrim
Segmentation
Statistics
Step 1. RL deconvolution
Step 2. SN2N inference
Step 3. Organelle segmentation
Step 4. Downstream analysis
3D-UNet
Convolution
Downsampling
Upsampling
Skip connection
RL-SN2N Segmentation
ERnet tubules
Mitonet
GA Otsu
Lys Otsu
0 h 0 min
1 h 0 min
1 h 10 min
3D RL-SN2N
Mito
Nucleus
SOFI-SN2N
SOFI data pair
SOFI
Step 1. N^th order cumulantion (SOFI)
Step 2. Diagonal resampling & Fourier ×4
Step 3. RL deconvolution
Steps 1-3. Self-supervised data generation
Step 4. Self-constrained learning
Step 2. Fourier re-scaling ×2
Step 1-3. SOFI data
Step 2. SN2N inference
2^nd SOFI 20f
2^nd SOFI-SN2N 20f
OMM
SIM-SN2N
Raw data pair
SIM data pair
Step 1. Diagonal resampling & Fourier ×2
Step 2. SIM reconstruction
Step 1. SIM reconstruction
SIM data
2D-SIM
IMM

### Supplementary Figures.

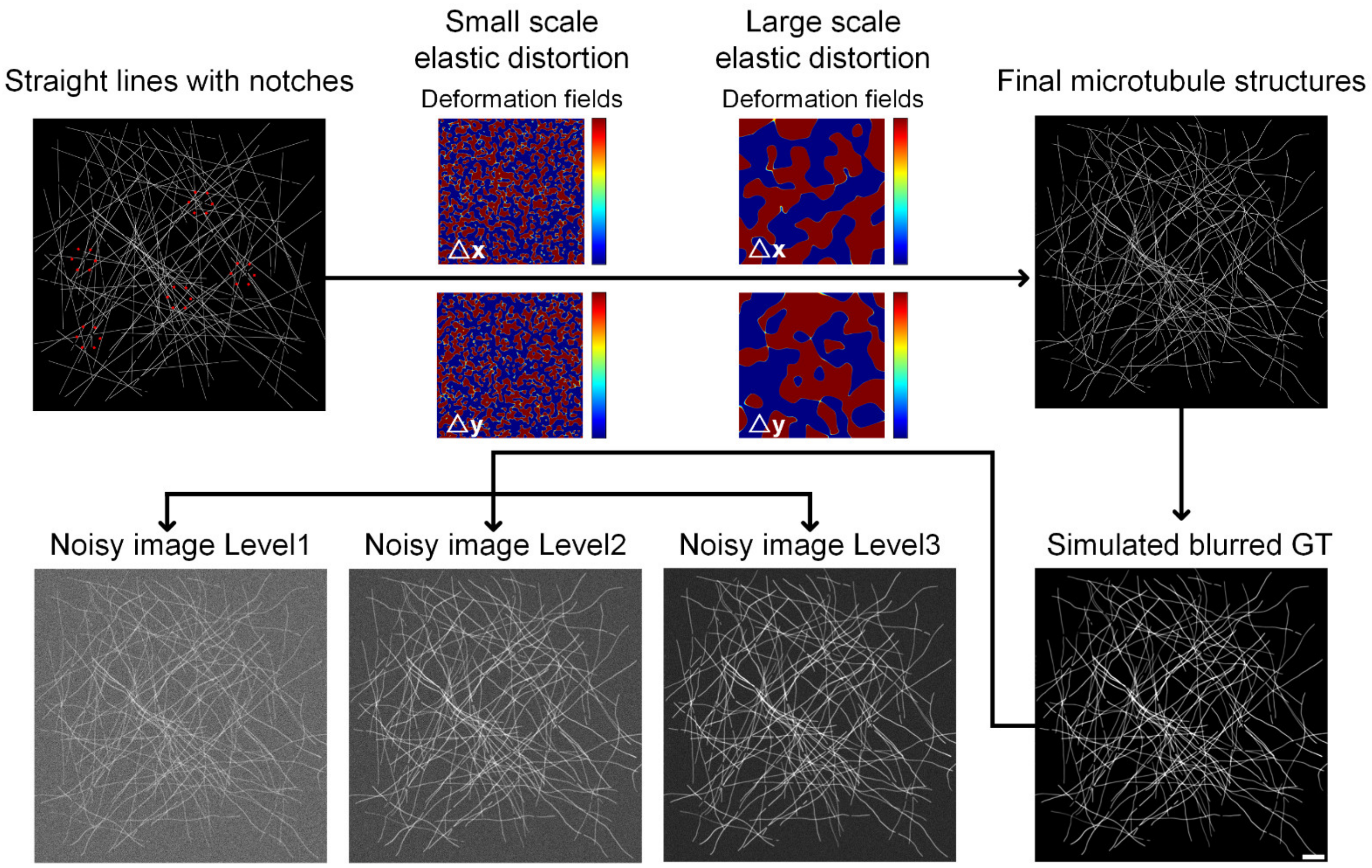


**Supplementary Fig. 1 | Simulation workflow of microtubule filaments, ground-truth (GT) image, and super-resolution (SR) images under different noise levels.** Scale bar, 5 μm.

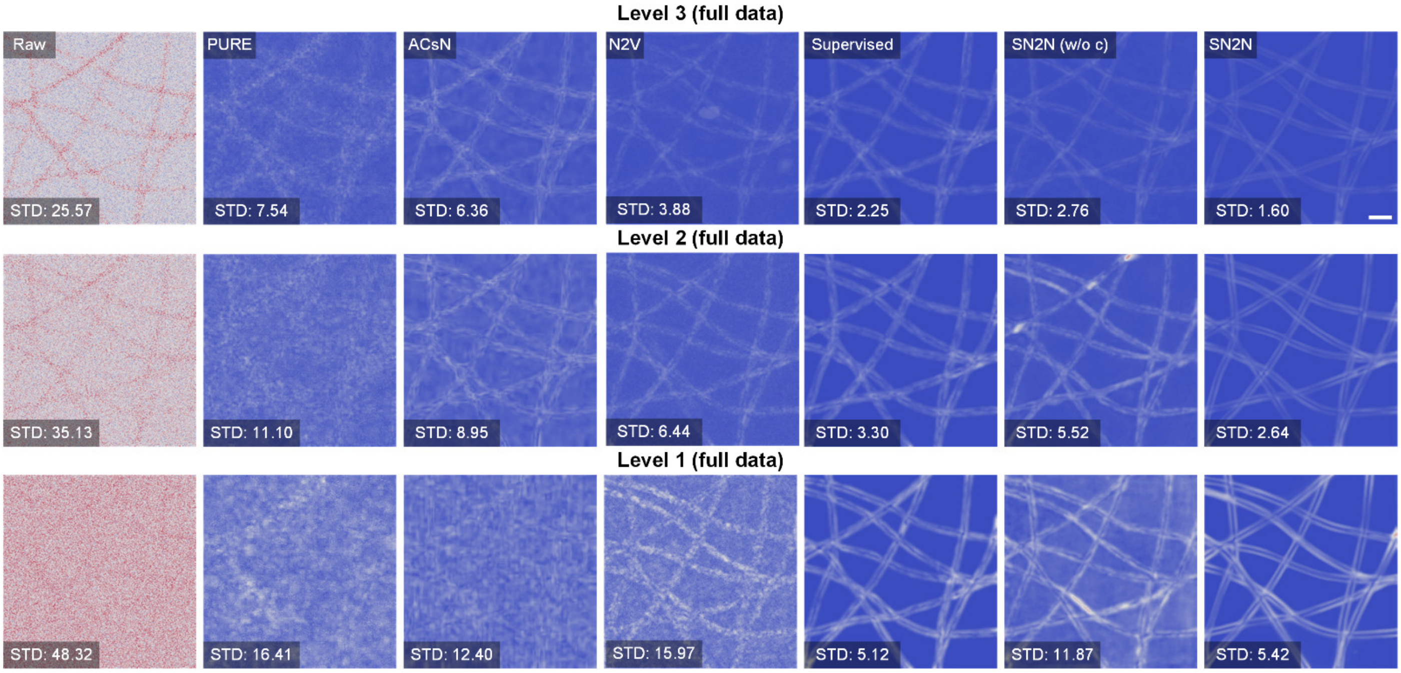


**Supplementary Fig. 2 | Full data uncertainty results of simulation data under different noise levels (*c.f.*, Extended Data Fig. 3c).** Scale bar, 1 μm.

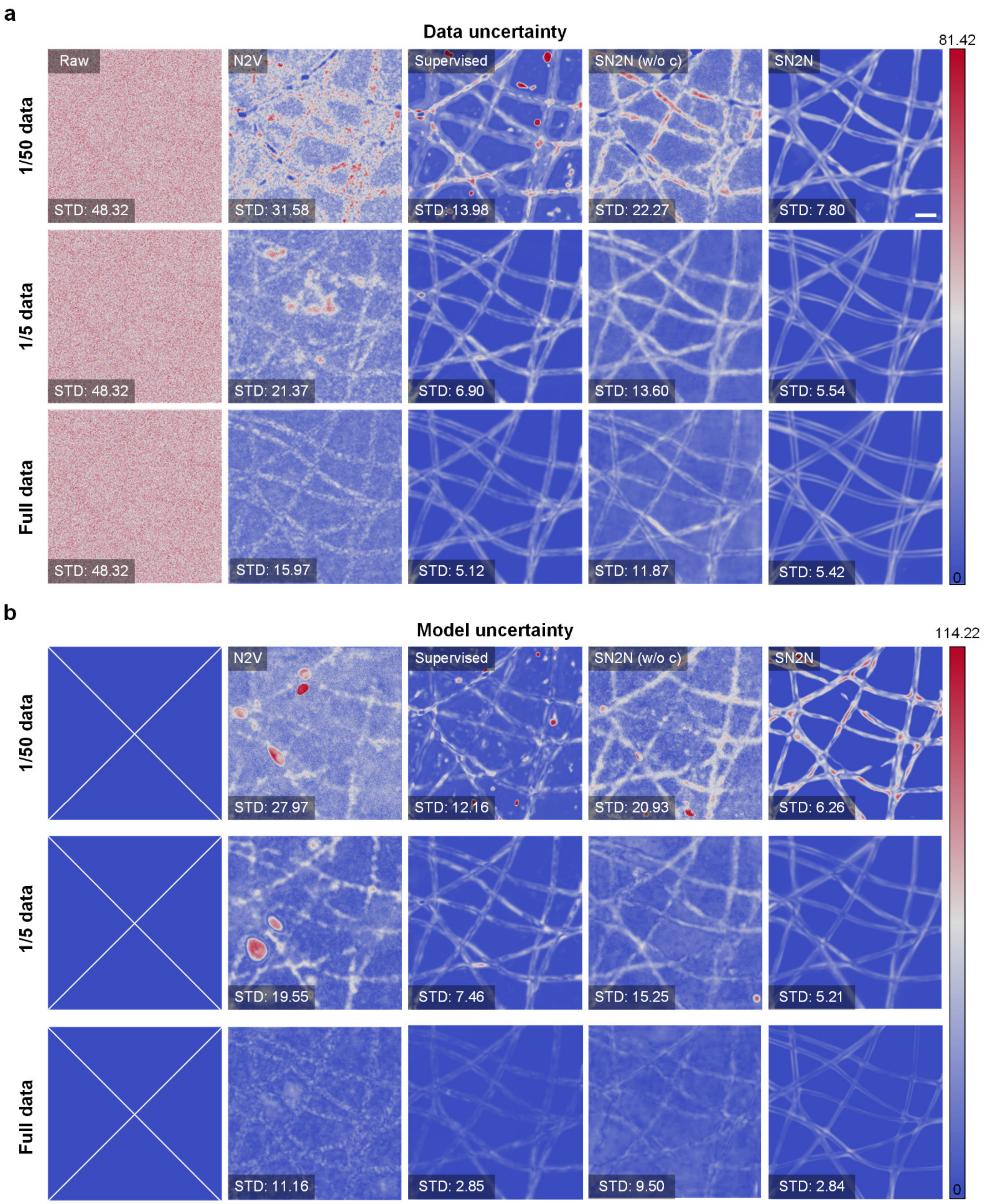


**Supplementary Fig. 3 | Full data uncertainty and model uncertainty results of simulation data with different training data amounts (*c.f.*, Extended Data Fig. 3a).** Scale bar, 1 μm.

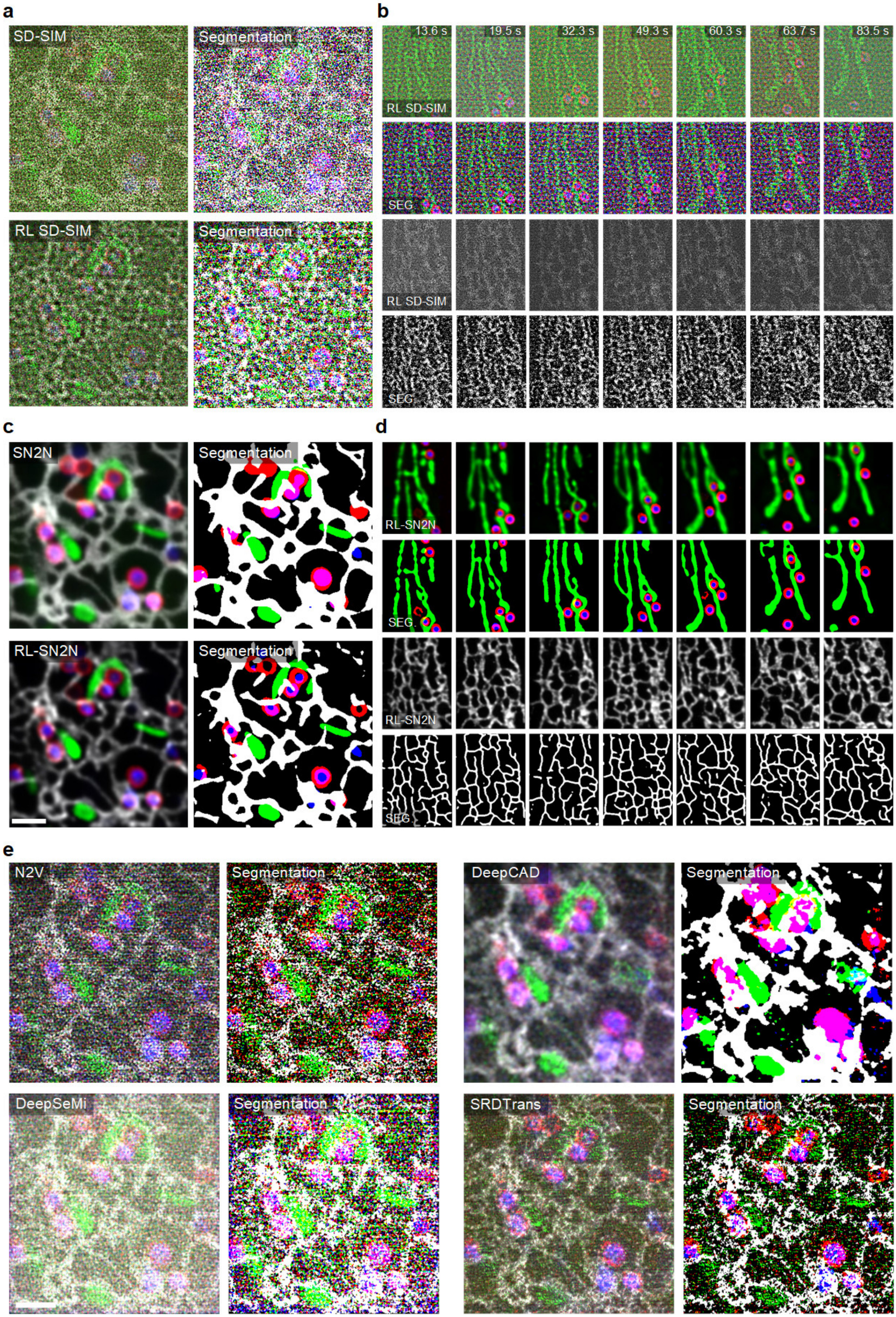

a
SD-SIM
Segmentation
RL SD-SIM
Segmentation
b
13.6 s
19.5 s
32.3 s
49.3 s
60.3 s
63.7 s
83.5 s
RL SD-SIM
SEG
RL SD-SIM
SEG
c
SN2N
Segmentation
RL-SN2N
Segmentation
d
RL-SN2N
SEG.
RL-SN2N
SEG.
e
N2V
Segmentation
DeepCAD
Segmentation
DeepSeMi
Segmentation
SRDTrans
Segmentation

**Supplementary Fig. 4 | Benchmarking of SN2N against N2V[1], DeepCAD[2], DeepSeMi[3], and SRDTrans[4] using live-cell imaging data and their segmentation results (*c.f.*, Fig. 3c).** SEG.: segmentation. The DeepCAD model was trained by temporally interleaved data (odd and even frames) as input and target. Scale bar, 1 µm.

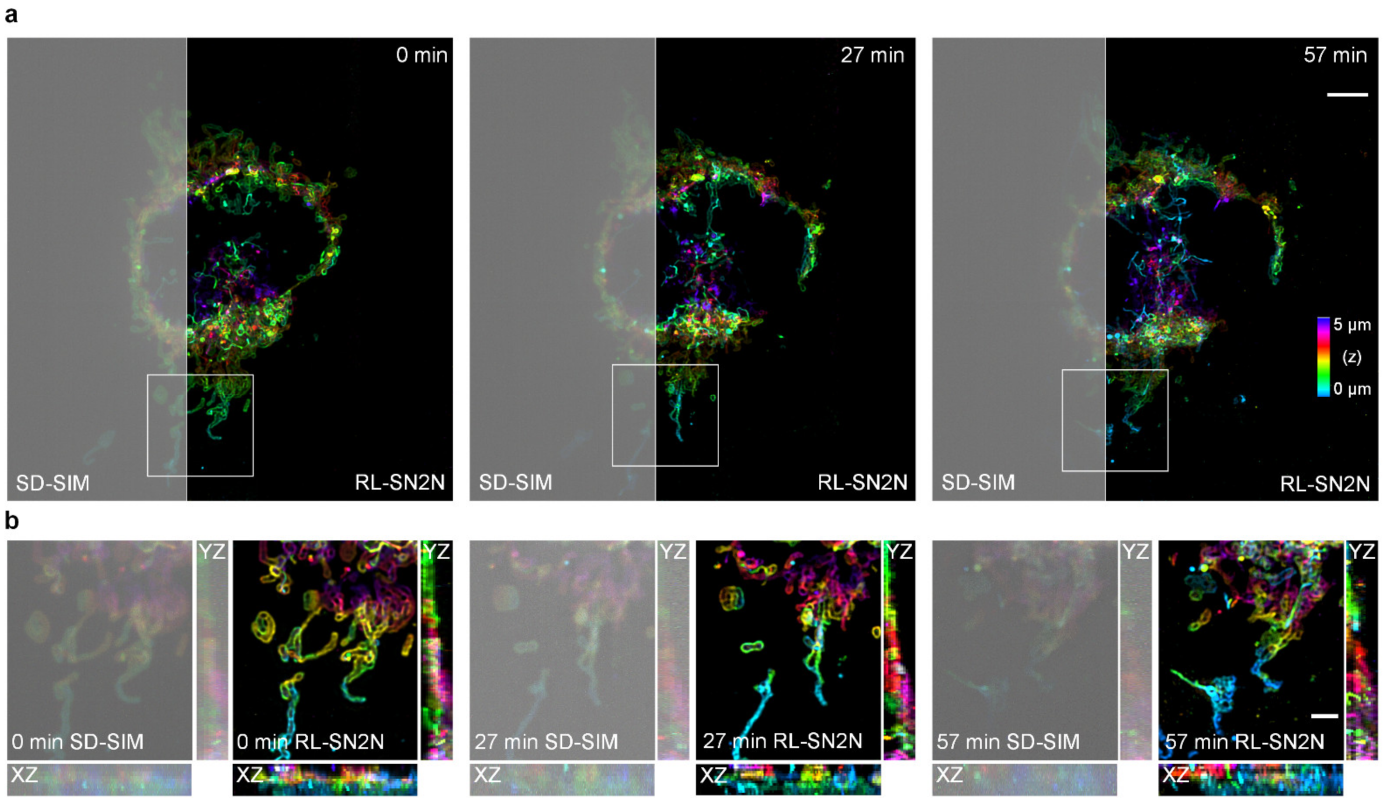


**Supplementary Fig. 5 | RL-SN2N reveals the 3D distribution of mitochondria after long-term recording (*c.f.*, Fig. 4e)**. **a**, Three time points of mitochondrial network in a live COS-7 cell labeled with Tom20-mCherry under SD-SIM (left) and RL-SN2N (right). **b**, Magnified views and their *xz* and *yz* cross-sections from white boxed regions in **a** under SD-SIM (left) and RL-SN2N (right). Scale bars, 5 µm (**a**); 2 µm (**b**).

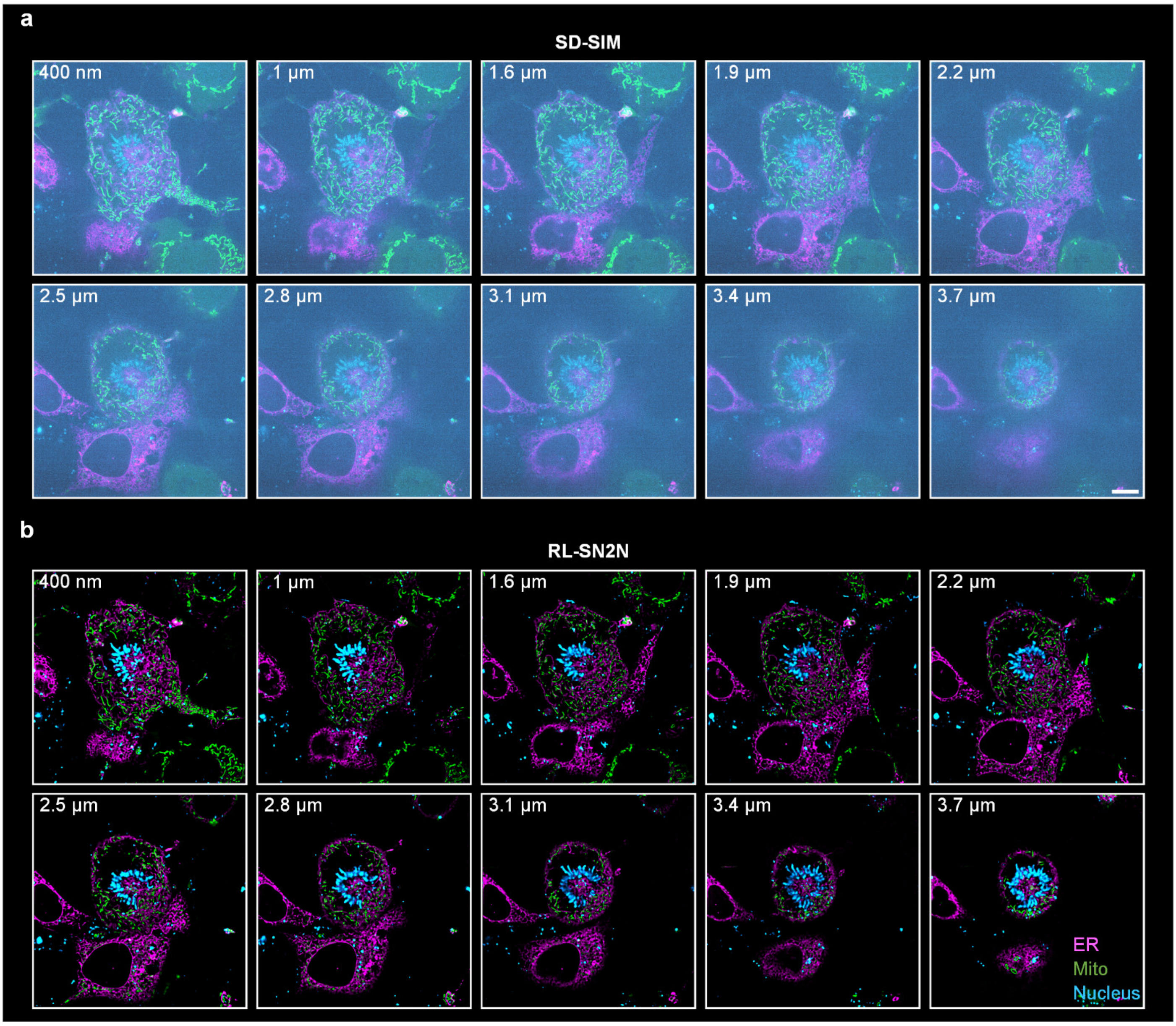


**Supplementary Fig. 6 | Representative lateral slices at the first time point of 5D data (*c.f.*, Fig. 4g). a**, **b**, SD-SIM (**a**) lateral slices at different axial positions (labeled on the top left corner) and their SN2N denoised results (**b**). Scale bar, 5µm.

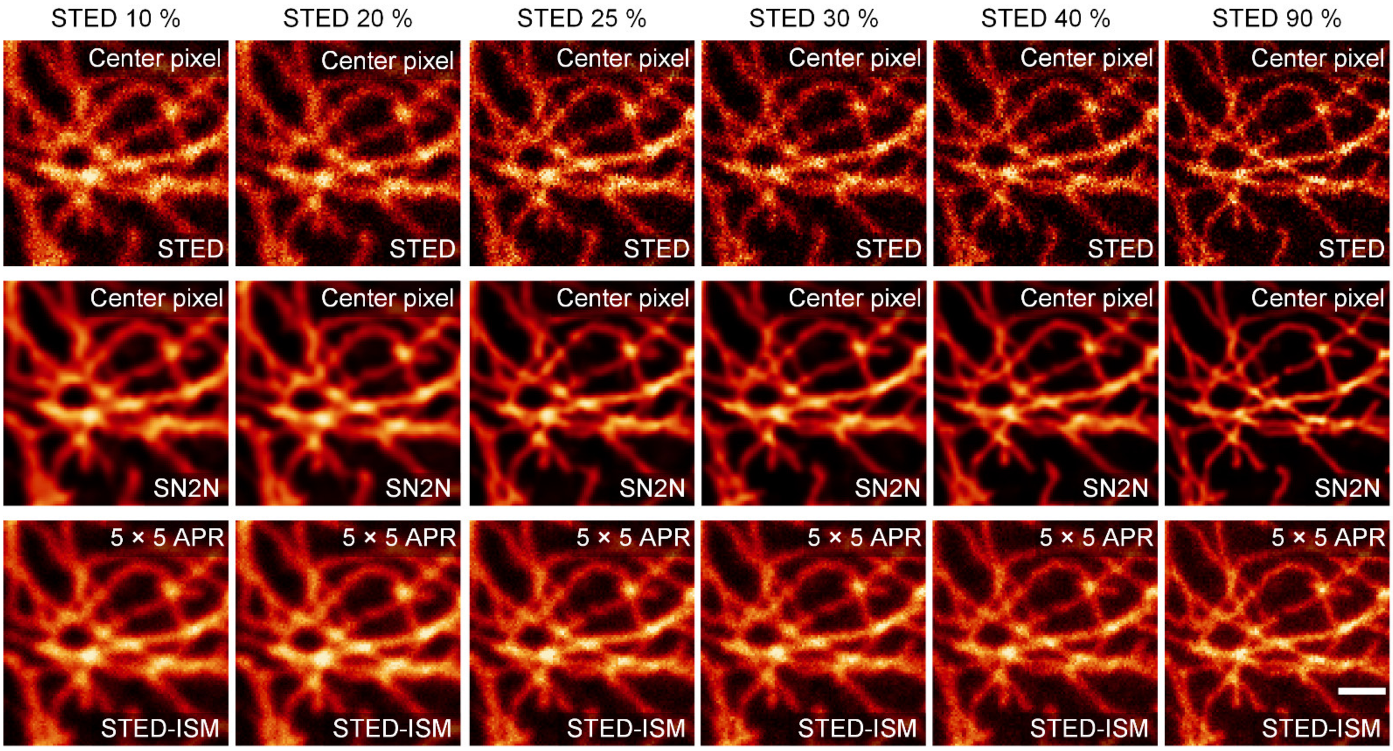


**Supplementary Fig. 7 | STED (top), STED-SN2N (middle), and STED-ISM (bottom) results of microtubules in a fixed cell under different depletion laser powers.** Data from ref.[5]. Scale bar, 1 μm.

### Supplementary Tables.

**Supplementary Table 1 | SSIM[6], PSNR, and RMSE values for denoising simulated microtubules under different noise levels (*c.f.*, Extended Data Fig. 3b).**

| Full data | | PURE | ACsN | N2V | Supervised | SN2N (w/o c) | SN2N |
|---|---|---|---|---|---|---|---|
| **SSIM** | **Level 1** | 0.3182 ± 0.0028 | 0.3654 ± 0.0004 | 0.4473 ± 0.0001 | 0.7278 ± 0.0001 | 0.5222 ± 0.0005 | **0.8032 ± 0.0004** |
| | **Level 2** | 0.4129 ± 0.0009 | 0.4306 ± 0.0002 | 0.5603 ± 0.0003 | 0.8370 ± 0.0001 | 0.5790 ± 0.0030 | **0.8400 ± 0.0000** |
| | **Level 3** | 0.5072 ± 0.0005 | 0.5101 ± 0.0001 | 0.7808 ± 0.0001 | 0.8926 ± 0.0000 | 0.8062 ± 0.0001 | **0.8882 ± 0.0000** |
| **PSNR** | **Level 1** | 14.6763 ± 1.2347 | 14.7244 ± 2.4205 | 15.2940 ± 0.0867 | 22.6393 ± 0.0422 | 16.6833 ± 0.2847 | **24.2803 ± 0.3170** |
| | **Level 2** | 16.1902 ± 0.8094 | 16.2847 ± 1.2442 | 16.7018 ± 0.1482 | 25.2417 ± 0.1346 | 18.1278 ± 0.2647 | **25.1182 ± 0.1410** |
| | **Level 3** | 20.2255 ± 0.5811 | 20.9137 ± 0.5247 | 21.9142 ± 0.1992 | 27.9624 ± 0.0173 | 22.5314 ± 0.1575 | **26.6088 ± 0.0221** |
| **RMSE** | **Level 1** | 0.1861 ± 0.0005 | 0.1865 ± 0.0011 | 0.1720 ± 0.0001 | 0.0738 ± 0.0000 | 0.1468 ± 0.0001 | **0.0613 ± 0.0000** |
| | **Level 2** | 0.1559 ± 0.0003 | 0.1546 ± 0.0004 | 0.1463 ± 0.0000 | 0.0539 ± 0.0000 | 0.1243 ± 0.0000 | **0.0555 ± 0.0000** |
| | **Level 3** | 0.0978 ± 0.0001 | 0.0903 ± 0.0001 | 0.0803 ± 0.0000 | 0.0401 ± 0.0000 | 0.0748 ± 0.0000 | **0.0467 ± 0.0000** |

**Supplementary Table 2 | SSIM, PSNR, and RMSE values for denoising simulated microtubules under different training data amounts (*c.f.*, Extended Data Fig. 3d).**

| Level 1 | | N2V | Supervised | SN2N (w/o c) | SN2N | SN2N (w a) |
|---|---|---|---|---|---|---|
| **SSIM** | **1/50 data** | 0.2091 ± 0.0002 | 0.3322 ± 0.0002 | 0.2916 ± 0.0001 | 0.7062 ± 0.0002 | **0.7360 ± 0.0001** |
| | **1/10 data** | 0.2783 ± 0.0001 | 0.5173 ± 0.0001 | 0.3764 ± 0.0023 | 0.7586 ± 0.0002 | **0.7760 ± 0.0002** |
| | **Full data** | 0.3591 ± 0.0001 | 0.7145 ± 0.0000 | 0.4826 ± 0.0001 | 0.8069 ± 0.0002 | **0.8232 ± 0.0001** |
| **PSNR** | **1/50 data** | 11.0758 ± 0.0441 | 13.7362 ± 0.1555 | 13.1180 ± 0.0828 | 20.6622 ± 0.1965 | **21.3920 ± 0.2242** |
| | **1/10 data** | 13.7447 ± 0.5549 | 18.0178 ± 0.0176 | 16.0825 ± 0.1418 | 21.7903 ± 0.6402 | **22.3012 ± 0.6223** |
| | **Full data** | 16.4204 ± 0.2085 | 22.3782 ± 0.0395 | 18.8657 ± 0.5431 | 23.0682 ± 0.4132 | **23.2253 ± 0.3057** |
| **RMSE** | **1/50 data** | 0.2795 ± 0.0001 | 0.2059 ± 0.0001 | 0.2209 ± 0.0001 | 0.0928 ± 0.0001 | **0.0854 ± 0.0001** |
| | **1/10 data** | 0.2067 ± 0.0005 | 0.1257 ± 0.0001 | 0.1571 ± 0.0001 | 0.8182 ± 0.0001 | **0.0772 ± 0.0001** |
| | **Full data** | 0.1512 ± 0.0001 | 0.0761 ± 0.0000 | 0.1144 ± 0.0000 | 0.0704 ± 0.0000 | **0.0691 ± 0.0000** |

**Supplementary Table 3 | LRQ[7], SSIM, and STD values for different denoising algorithms on the commercial Argo-SIM slide under SpinSR10 SD-SIM system (*c.f.*, Fig. 2b-2e).**

| | | SD-SIM | PURE | N2V | SN2N (w/o c) | SN2N |
|---|---|---|---|---|---|---|
| LRQ | 120 nm | 0.1673 | 0.1707 | 0.1808 | 0.1954 | **0.2119** |
| | 150 nm | 0.1747 | 0.1834 | 0.2035 | 0.2203 | **0.2690** |
| | 180 nm | 0.1843 | 0.1941 | 0.2188 | 0.2501 | **0.2951** |
| | 210 nm | 0.1969 | 0.2119 | 0.2388 | 0.2720 | **0.3431** |
| | 240 nm | 0.2047 | 0.2292 | 0.2613 | 0.2973 | **0.3846** |
| SSIM | | 0.2715 ± 0.0001 | 0.5306 ± 0.0014 | 0.6200 ± 0.0012 | 0.7018 ± 0.0231 | **0.8974 ± 0.0001** |
| STD | | 19.6506 ± 0.0038 | 18.3152 ± 0.0203 | 8.0821 ± 0.0231 | 4.2259 ± 0.0017 | **1.2241 ± 0.0501** |

**Supplementary Table 4 | SSIM values for denoising microtubules in fixed COS-7 cells under different training data amounts (*c.f.*, Fig. 2h).**

| | | Supervised (w/o aug) | SN2N (w/o aug) |
|---|---|---|---|
| SSIM | 1 | 0.5127 ± 0.0008 | **0.7643 ± 0.0041** |
| | 10 | 0.6358 ± 0.0034 | **0.7965 ± 0.0017** |
| | 100 | 0.6947 ± 0.0007 | **0.8451 ± 0.0028** |
| | 500 | 0.7398 ± 0.0010 | **0.8652 ± 0.0002** |

**Supplementary Table 5 | SSIM values for denoising microtubules in fixed COS-7 cells under different data augmentation strategies (*c.f.*, Fig. 2j).**

| | SSIM |
|---|---|
| SN2N-1f with basic augmentation | 0.7928 ± 0.0042 |
| SN2N-1f with Patch2Patch augmentation | 0.8488 ± 0.0035 |
| SN2N-1f with full augmentation | **0.8631 ± 0.0009** |
| SN2N-500f without augmentation | 0.8588 ± 0.0005 |

**Supplementary Table 6 | SSIM values for denoising microtubules in fixed COS-7 cells under different exposure time (*c.f.*, Fig. 2i).**

| | | Supervised-1f (w/o aug) | SN2N-1f (w/o aug) |
|---|---|---|---|
| SSIM | 1 × | 0.5062 ± 0.0014 | **0.7659 ± 0.0032** |
| | 2 × | 0.5742 ± 0.0019 | **0.8135 ± 0.0036** |
| | 5 × | 0.6931 ± 0.0018 | **0.8287 ± 0.0042** |
| | 10 × | 0.8146 ± 0.0041 | **0.8677 ± 0.0040** |

### Captions for Supplementary Videos.

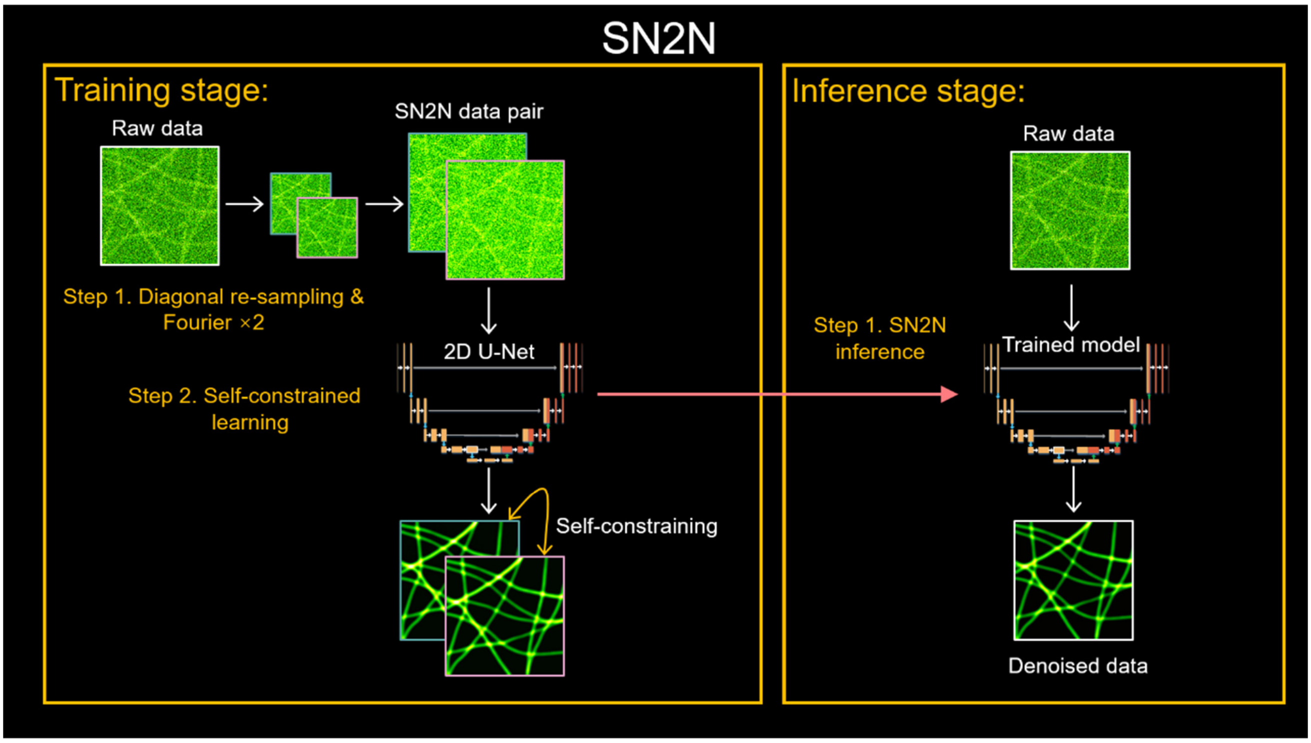


**Supplementary Video 1 | SN2N workflows.** Part I provides the detailed training workflow of SN2N (*c.f.*, **Extended Data Fig. 1**). Part II showcases the workflows of SN2N, RL-SN2N, 3D RL-SN2N, SOFI-SN2N, SIM-SN2N (raw re-sampling) and SIM-SN2N (SIM re-sampling).

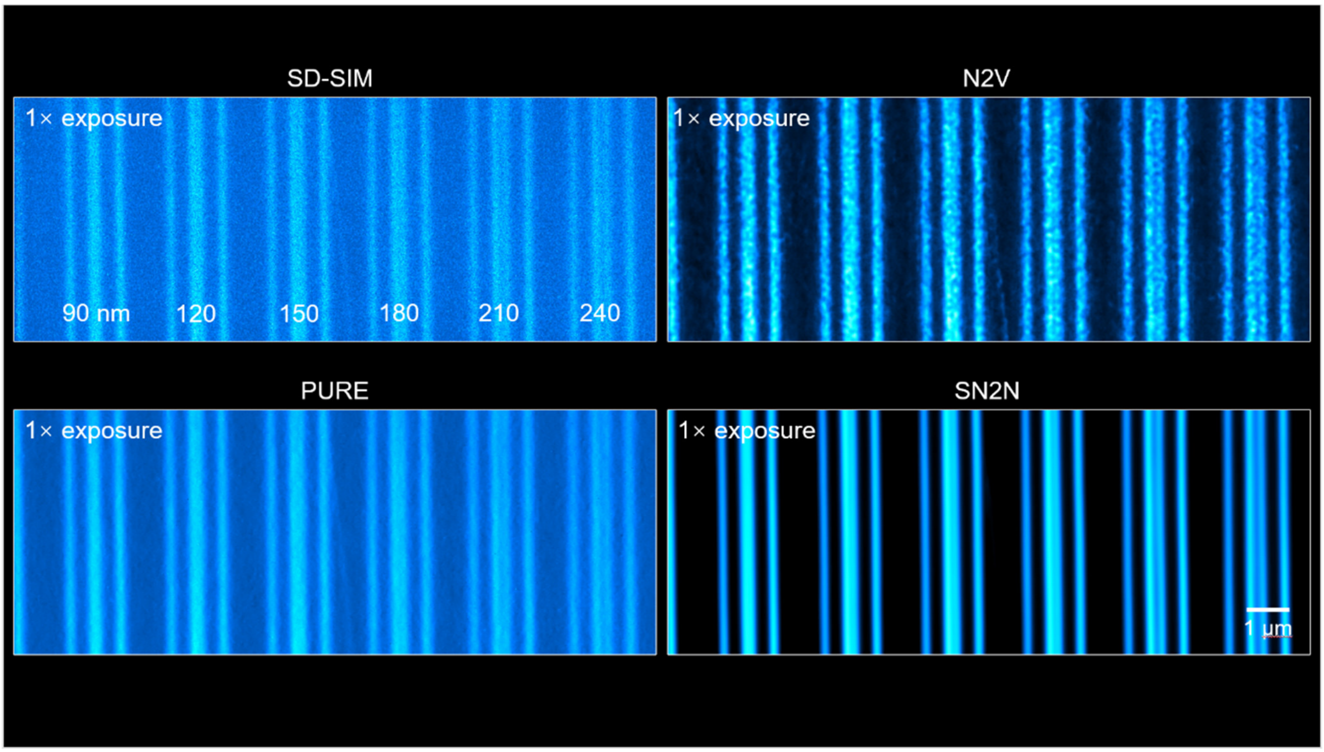


**Supplementary Video 2 | SN2N eliminates noise and noise fluctuations in known structures.** Part I shows the comparisons of SD-SIM, PURE, N2V, and SN2N on the commercial Argo-SIM slide under the SpinSR10 SD-SIM system across various exposure conditions (*c.f.*, **Fig. 2a**). Part II shows the noise fluctuations captured by SD-SIM and SN2N under 1× exposure condition.

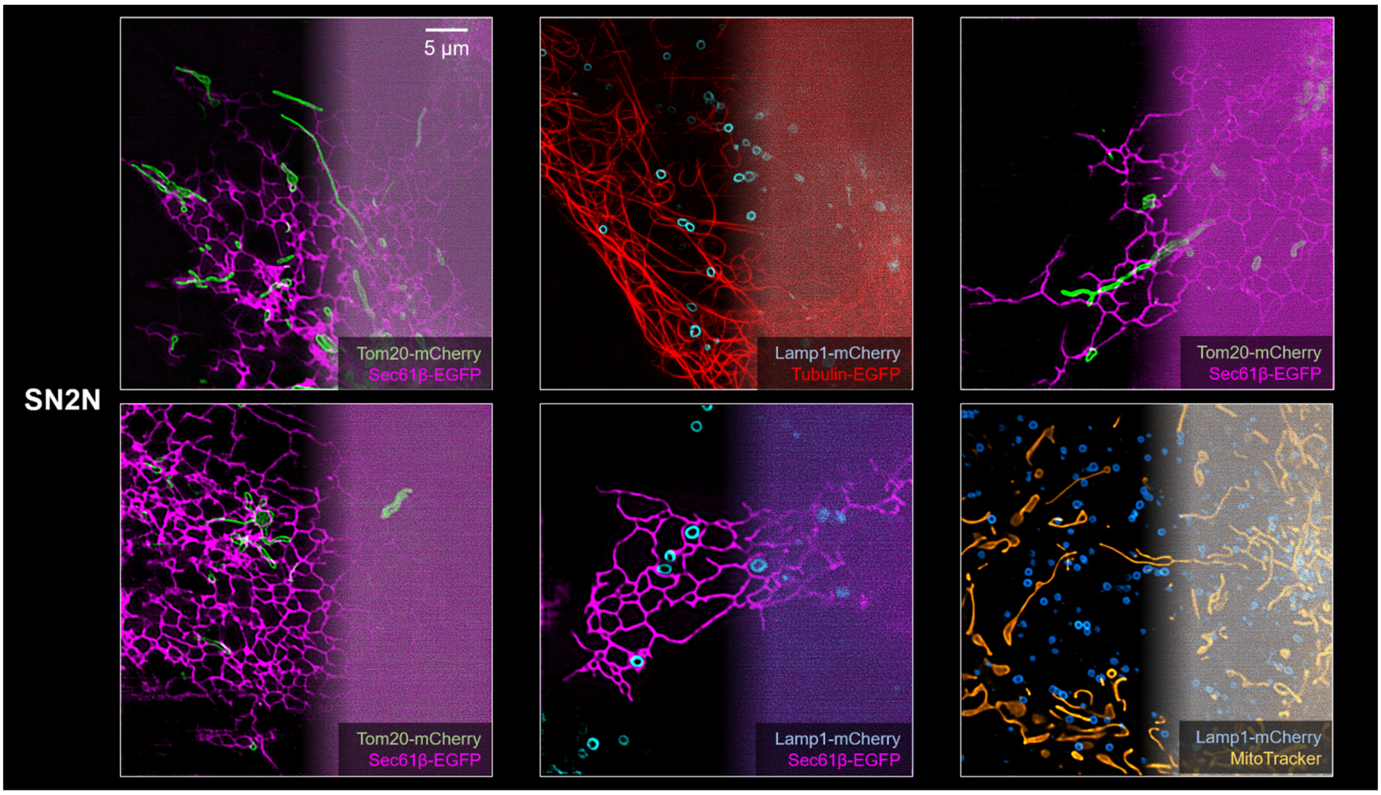


**Supplementary Video 3 | RL-SN2N enables dual-color live-cell SR SD-SIM imaging.** Part I shows the fission events of mitochondria (Mito, green) and ER (magenta) labeled with Tom20-mCherry and Sec61β-EGFP in live COS-7 cells captured by SD-SIM and RL-SN2N (*c.f.*, **Extended Data Fig. 5d**). The yellow arrows indicate the mitochondrial fissions. Part II demonstrates a live-cell SR imaging gallery of live COS-7 cells labeled with Tom20-mCherry (Mito, green), Sec61β-EGFP (ER, magenta), Lamp1-mCherry (Lys, cyan), Tubulin-EGFP (Tubulin, red), or MitoTracker (Mito, orange) captured by SD-SIM and RL-SN2N.

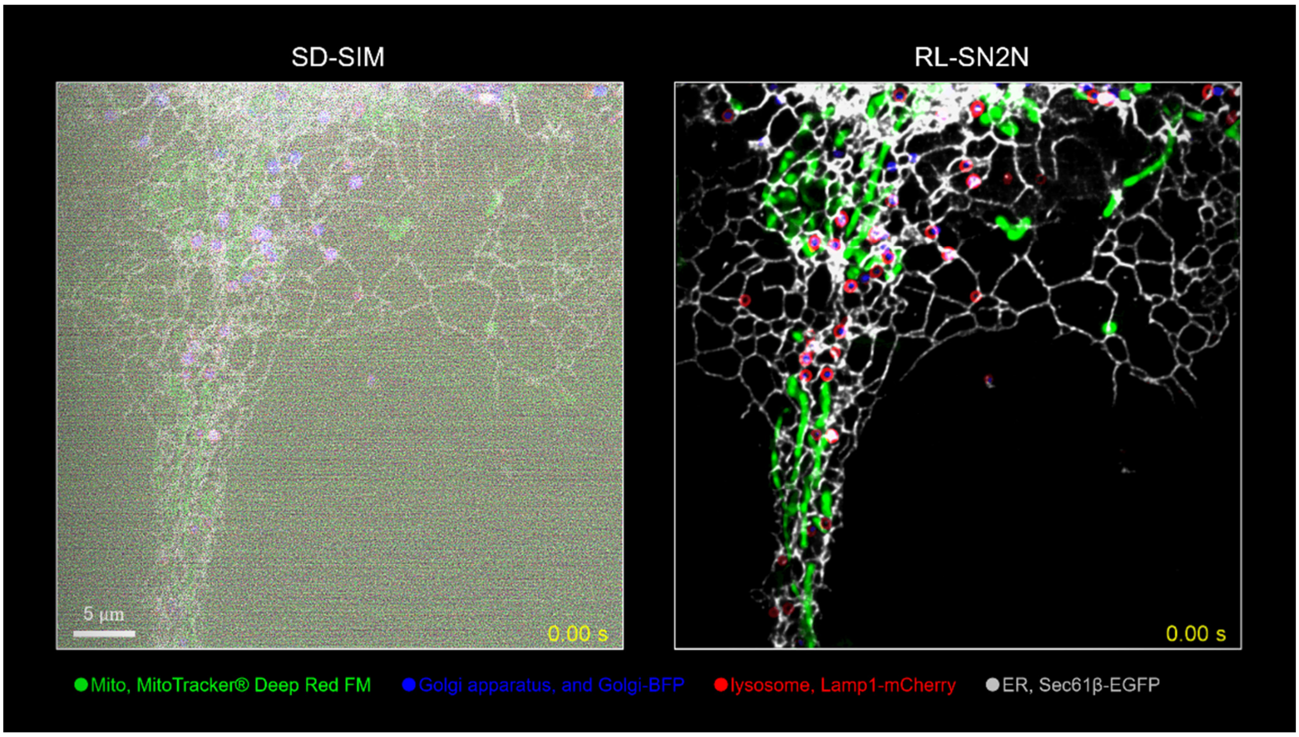


**Supplementary Video 4 | RL-SN2N unlocks four-color live-cell SR imaging and facilitates the downstream analysis.** Part I shows the four-color imaging of Mito (green), ER (gray), lysosome (Lys, red), and Golgi apparatus (GA, blue) labeled with MitoTracker® Deep Red FM, Sec61β-EGFP, Lamp1-mCherry, and Golgi-BFP in live COS-7 cells under raw SD-SIM and RL-SN2N (*c.f.*, **Fig. 3c**). Part II highlights the four-color segmentation results under RL-SN2N. Part III provides the trajectories and spatial distributions of lysosomes assigned with different motion behaviors. Part IV tracks multiple organelle interaction events captured by RL-SN2N.

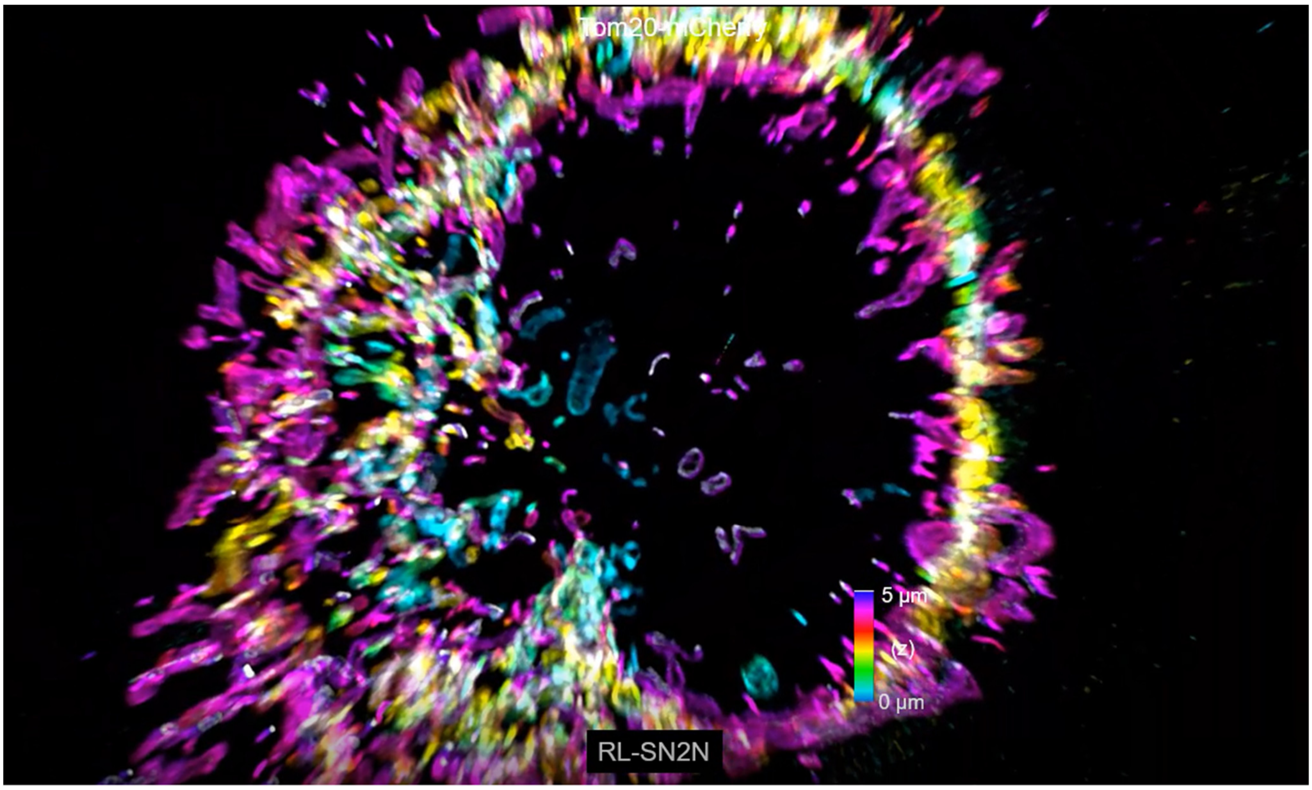


**Supplementary Video 5 | 3D RL-SN2N offers volumetric SR imaging of the outer mitochondrial membrane (OMM) network in live cells.** Part I shows the 3D color-coded volumes of the OMM network in live COS-7 cells labeled with Tom20–mCherry, captured by raw SD-SIM and RL-SN2N (*c.f.*, **Fig. 4b**). Part II, OMM atlas. High-quality 3D SR data enables us to easily map and manipulate each mitochondrion. Part II demonstrates the 4D imaging of OMM network in live COS-7 cells captured by SD-SIM and RL-SN2N (*c.f.*, **Fig. 4e**).

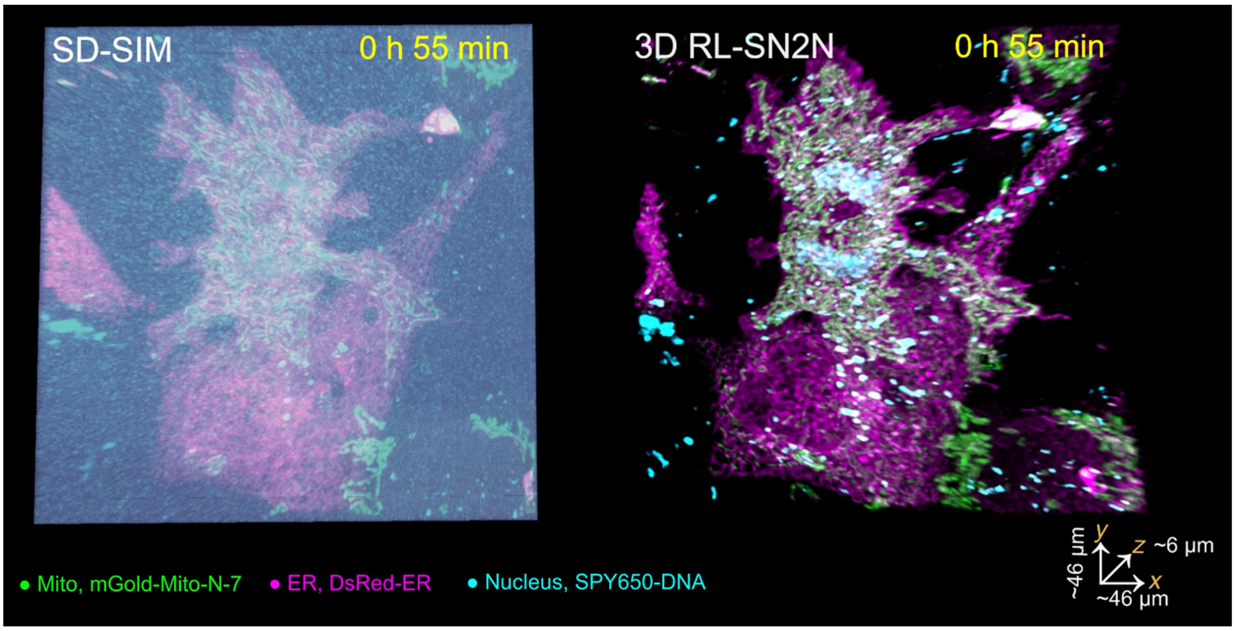


**Supplementary Video 6 | 3D RL-SN2N on SD-SIM enables fast long-term live-cell SR imaging across 5D, recording the entire cell mitosis process.** Part I shows the 5D imaging of mitochondria (green, mGold-Mito-N-7), ER (magenta, DsRed-ER), and nucleus (cyan, SPY650-DNA) in live COS-7 cells under raw SD-SIM and RL-SN2N (*c.f.*, **Fig. 4g**). Part II displays the dynamics of mitochondrial (green) and ER (magenta) networks post mitosis.

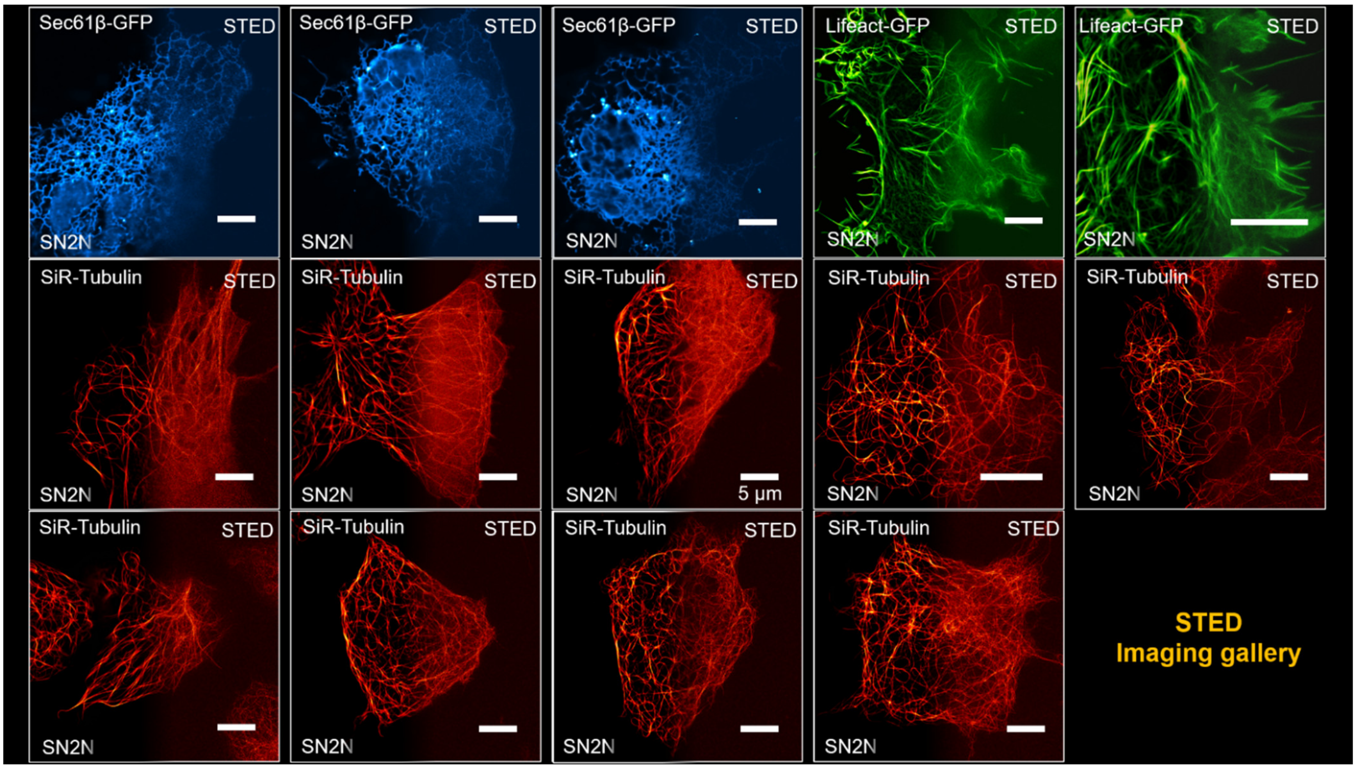


**Supplementary Video 7 | SN2N and RL-SN2N permit long-term live-cell STED imaging.** Part I shows a live-cell SR imaging gallery of live COS-7 cells labeled with SiR-Tubulin (left), Lifeact-EGFP (middle), and Sec61β–EGFP (right) captured using a commercial STED microscope (Leica) and enhanced by SN2N (*c.f.*, **Fig. 5g**). Part II features long-term live-cell imaging of mitochondrial cristae in PKMO-labeled COS-7 cells, captured with a commercial STED system (Abberior) under various conditions, including high depletion power (86%) with both long (100 μs per pixel) and short (10 μs per pixel) durations, as well as low depletion power (41%) with a short duration (10 μs per pixel) (*c.f.*, **Fig. 5i-m**). Part III presents a comparison of long-term live-cell super-resolution imaging of mitochondrial cristae using STED and RL-SN2N.

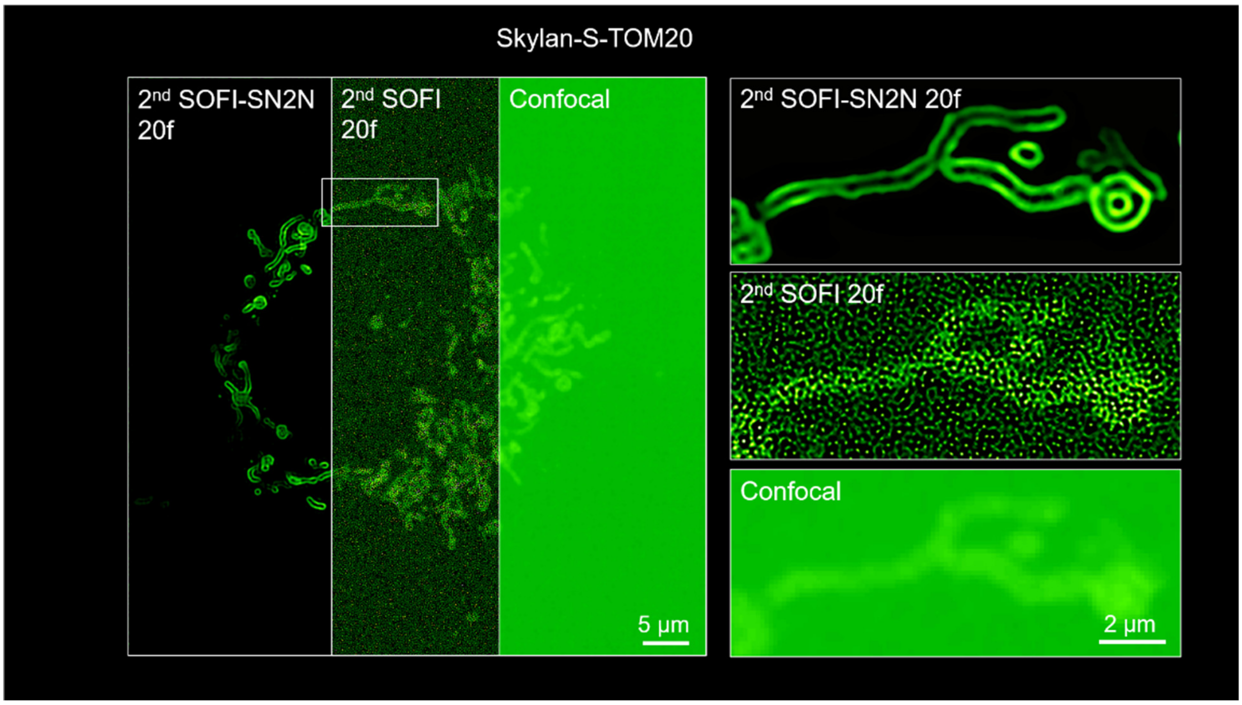


**Supplementary Video 8 | SOFI-SN2N supports efficient SOFI reconstruction in both fixed and live cells.** Part I demonstrates a representative COS-7 cell labeled with $QD_{525}$, imaged with a commercial SD-confocal microscope and reconstructed using $2^{nd}$, $3^{rd}$, and $4^{th}$ orders SOFI, along with its SOFI-SN2N results across 20, 50, 100, 200, 500, 1000 frames (*c.f.*, **Fig. 6e-6f**). Part II compares the OMM structures in a live COS-7 cell labeled with Skylan-S-TOM20, captured by SD-confocal, $2^{nd}$ order SOFI with 20 frames, and its SOFI-SN2N results.

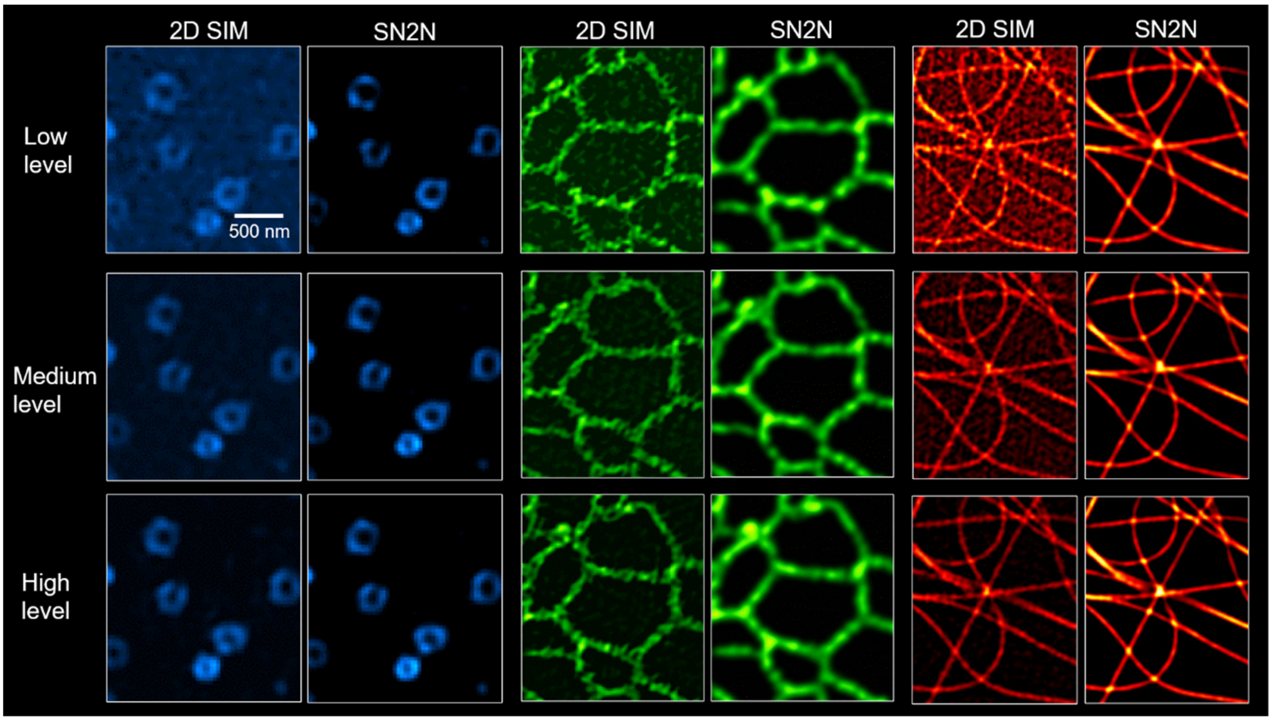


**Supplementary Video 9 | SN2N removes random, non-continuous artifacts in SIM reconstructions.** Part I shows the denoising results of SIM-SN2N (raw re-sampling method) on the BioSR SIM dataset under high, medium, and low SNR levels (*c.f.*, **Extended Data Fig. 8b-8g**). Part II compares the mitochondrial cristae structures in live COS-7 cells labeled with MitoTracker Green by 2D-SIM and SN2N-SIM (raw re-sampling method) (*c.f.*, **Extended Data Fig. 8h-8j**).
